# Engineered Whole Lungs for Tissue Biology

**DOI:** 10.1101/2024.10.02.616240

**Authors:** Allison Marie Greaney, Micha Sam Brickman Raredon, Tomohiro Obata, Juan Wang, Taylor Adams, Jonas Schupp, Satoshi Mizoguchi, Sophie Edelstein, Yifan Yuan, Pavlina Baevova, Nuoya Wang, Alex Engler, Katherine Leiby, Tomoshi Tsuchiya, Robert Homer, Naftali Kaminski, Robert Langer, Laura E. Niklason, Ruslan Medzhitov

## Abstract

End-stage lung disease and lung cancer significantly contribute to global mortality, necessitating new research strategies for studying pulmonary biology. Here, we present an engineered whole-lung tissue model used to evaluate the effects of cellular communities on tissue organization and alveolar barrier function. Engineered lungs were grown *ex vivo* on decellularized whole-lung matrices as structurally biomimetic, bioactive scaffolds. Histologic architecture of engineered lungs improved with the addition of alveolar macrophages, coming to resemble neonatal lung. Incorporating alveolar macrophages maximized the differentiation of native-like cellular communities, including alveolar type I-like epithelium, bronchioalveolar stem cells, microvascular endothelium, and pericytes. Cell-cell signaling in engineered lungs showed activation of developmental and inflammatory pathways, including WNT, Notch, and FGF signaling pathways. Engineered lungs containing alveolar macrophages showed a 668% improvement in measured alveolar barrier function. This work demonstrates the potential utility of engineered lung models for studying principles of tissue biology and pulmonary regeneration.

## Introduction

End-stage lung disease is the third leading cause of death worldwide, accounting for over 3 million deaths annually^1-3^. The only effective treatment is transplantation, which has a 10-year survival rate of about 25%, the worst long-term performance of all solid organ transplants^4^. Additionally, lung cancer is the deadliest form of cancer worldwide, accounting for 1.8 million deaths annually^5,6^. Such significant global health burden and patient morbidity calls for revolutionary modalities for studying lung dysfunction and potential routes for regeneration.

Pioneered in 2010, whole-lung tissue engineering utilizes organ decellularization and cellular repopulation to generate new tissues *in vitro*^7,8^. In this process, cells are removed from native lungs by detergent-enzymatic treatment, which preserves the composition and architecture of the extracellular matrix (ECM), then scaffolds are seeded with new cells and cultured in a bioreactor under controlled conditions to promote tissue-wide regeneration^9-11^. The initial goal for these engineered lungs was to function on transplantation, however early attempts were limited by incomplete recellularization and imperfect blood-gas barrier^8^. While transplantation remains a viable long-term goal, we have advanced this technology to serve as a powerful model system for studying tissue organization and regeneration *ex vivo*.

Here we present next-generation engineered whole lungs, and used them to study principles of tissue biology. We evaluated the effects of building co-culture on self-organization, thereby revealing a major role for alveolar macrophages in promoting tissue-level regeneration. We leveraged single-cell RNA sequencing (scRNAseq) to characterize phenotype and signaling landscapes within our engineered lungs, which further allowed us to observe patterns of functional delegation. Finally, we measured how multicellularity significantly improved alveolar barrier function *in vitro* and *in vivo*. Taken together, this work demonstrates the utility of engineered tissues as advanced *in vitro* model systems for answering new biologic questions.

## Results

### Engineered whole lungs as a model for studying tissue biology

We leveraged engineered whole rat lungs as model tissues for studying the effects of cellular community on tissue self-organization, with a particular goal of regenerating alveolar barrier function. Scaffolds were generated by decellularizing native rat lungs using detergents and enzymes to remove all cellular material, leaving behind intact extracellular matrix (ECM)^9^ (Fig. 1A). Decellularized ECM scaffolds preserve many localized properties of native tissues, including protein composition, architecture, and biomechanics, which have been shown to provide instructive cues to cells seeded into the environment^11-13^ (Fig. 1B). All starting cell populations were primary rat cells, including pharmacologically expanded basal cells (peBC) as epithelium^13^, rat lung microvascular endothelial cells (RLMVEC), pulmonary fibroblasts^14^, and alveolar macrophages from bronchoalveolar lavage (BAL) (Fig. 1C). Endothelial cells were seeded into the pulmonary artery and vein of the decellularized scaffold, whereas all other cell types were seeded into the air compartment via the trachea (Fig. 1D). Recellularized engineered lungs were then cultured in a bioreactor under controlled conditions for a week, undergoing continual pressure measurements and daily metabolic checks (Fig. 1E). For further details of the whole-lung tissue engineering protocol, see Methods.

**Figure 1.**
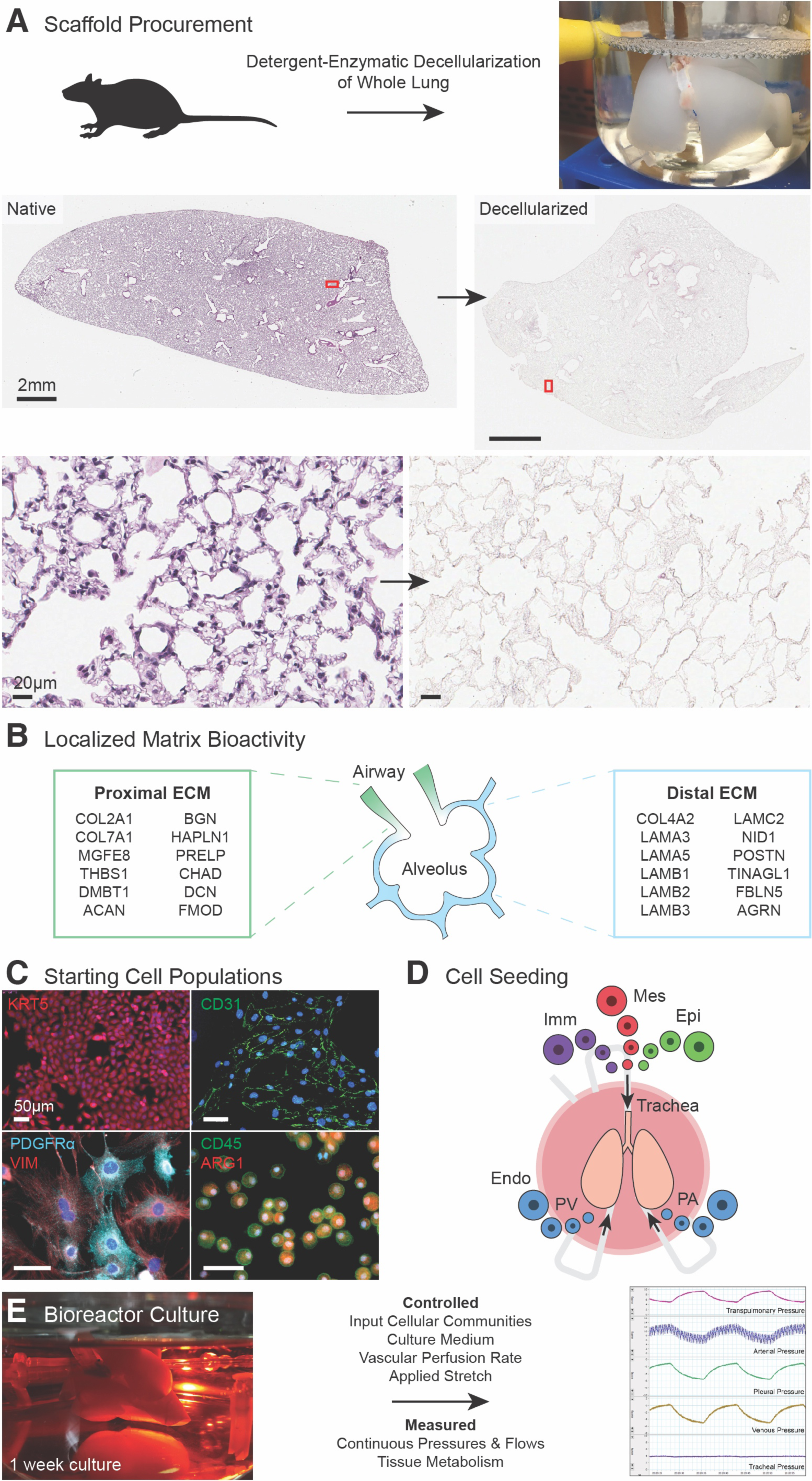
Engineered whole lungs as a model for studying tissue biology. A) Decellularization of whole native rat lungs to generate ECM scaffold for engineering. B) Localized protein enrichment in proximal and distal lung ECM 12. C) Immunofluorescent (IF) staining of starting cell populations: KRT5+ peBC, CD31+ RLMVEC, PDGFRα+/VIM+ pulmonary fibroblasts, CD45+/ARG1+ alveolar macrophages. D) Seeding each cell population into respective compartments of the lung while mounted in a bioreactor. E) Controlled and measured parameters during bioreactor culture of engineered lungs.

### Tissue organization changes with input cellular communities

To study the effects of building co-culture on cell phenotype and signaling, our experimental conditions included epithelial-only cultures (mono-culture), epithelium + endothelium (co-culture), epithelium + endothelium + mesenchyme (tri-culture), and epithelium + endothelium + mesenchyme + immune (quad-culture). By histology, mono-culture lungs displayed complete repopulation of the air compartment, with a mixture of cuboidal and thin cells in the alveoli (Fig. 2A). Co-culture lungs showed greater septal thinning, but also severe matrix degradation and debris (Fig. 2B). Tri-culture lungs displayed significant fibroblast overgrowth, which histologically resembled clinical fibroplasia (Fig. 2C). Notably, quad-culture lungs showed reduced fibroblast overgrowth and debris, enhanced septal thinning, and histologically resembled fetal lung tissue (Fig. 2D).

**Figure 2.**
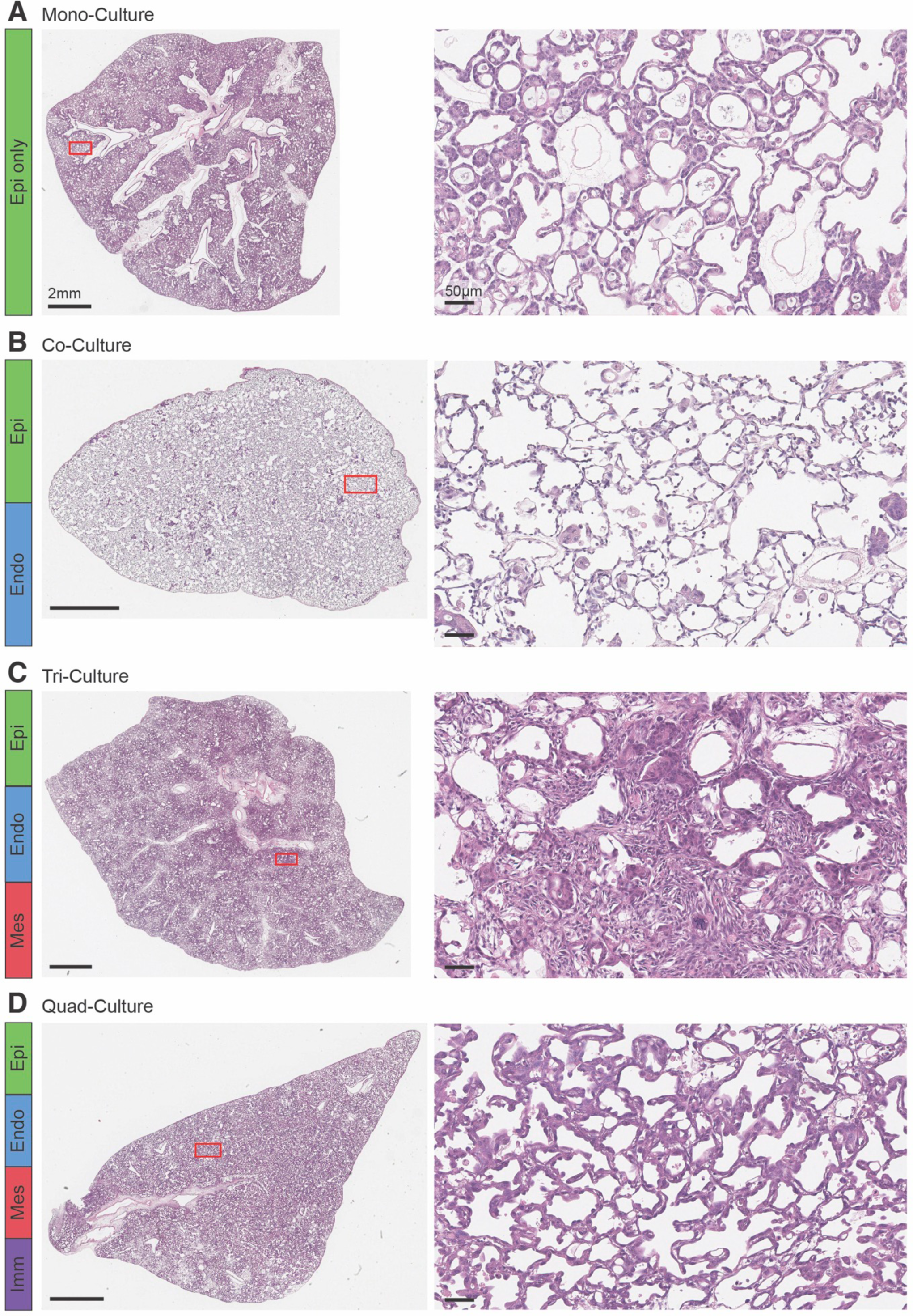
Tissue organization changes with input cellular communities. Slide scan and 20x H&E images of A) mono-culture engineered lung, B) co-culture engineered lung, C) tri-culture engineered lung, and D) quad-culture engineered lung.

### Heterogeneous alveolar archetypes arise in engineered lungs including rare cell types

We next evaluated cell phenotype and signaling in the engineered lungs using single-cell RNA sequencing (scRNAseq). We generated scRNAseq data of each experimental condition, including samples of starting cell populations (108,899 cells total). Following data cleaning (see Methods & Data Cleaning Supplemental Document), samples were divided by class for phenotype analysis. Cells in each class were integrated (Seurat CCA) to produce a unified embedding from which to perform cluster-based archetype identification. Engineered epithelium divided into two main archetypes: a *Krt5*+ “Basal-like” archetype, which mostly consisted of starting peBC, and a *Hopx*+ “ATI-like” archetype, which expressed genes associated with alveolar barrier formation (Fig. 3A-C). Endothelium displayed a “Microvascular” population, which gained expression of several key capillary genes in engineered lung cultures, including aCap/aerocyte markers *Kdr*, *Emcn*, and *Ednrb*, as well as gCap/general capillary markers *Gpihbp1*, *Vwf*, and *Vegfa^15,16^* (Fig. 3D-F). Engineered mesenchyme tended toward an alveolar fibroblast phenotype, expressing markers such as *Pdgfra* and *Tg@r3^17^* (Fig. 3G-I). Alveolar macrophages activated to a unique inflammatory state in engineered lungs, including expression of *Cxcl2* and *S100a9* (MRP-14)^18,19^ (Fig. 3J-L). We further identified rare cell types in engineered lungs, including *Sftpc*+/*Scgb1a1*+ bronchioalveolar stem cells (BASCs) and *Gucy1a1*+ pericytes, both of which closely resembled their native counterparts by top markers^17,20,21^. For additional analysis of engineered cell phenotypes, see Supp. Fig. 1-4.

**Figure 3.**
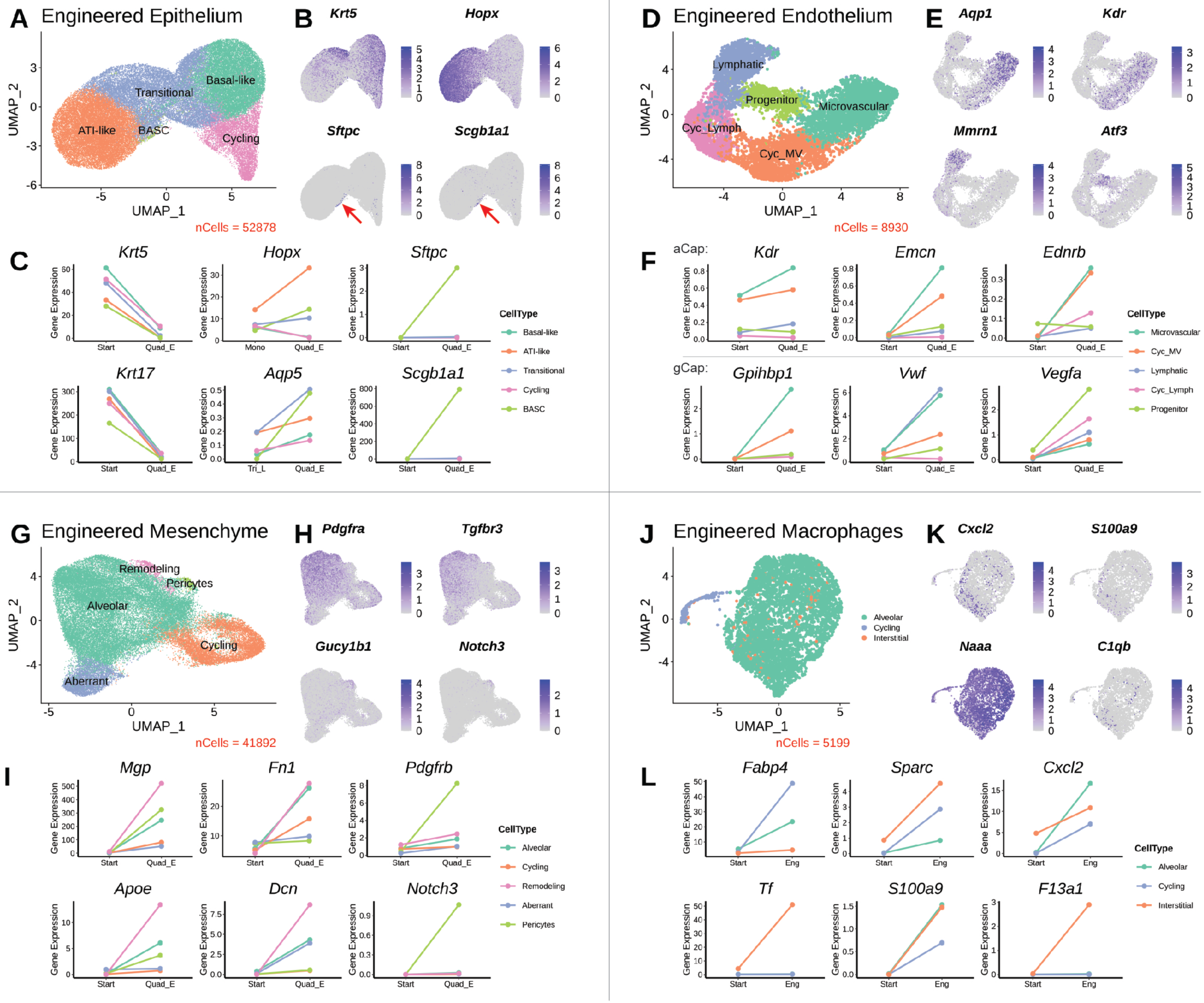
Heterogeneous alveolar archetypes arise in engineered lungs including rare cell types. A) Archetypes of engineered epithelium. B) Krt5, Hopx, Sftpc, and Scgb1a1 expression defining archetypes in engineered epithelium. C) Changes in relevant epithelial gene expression in engineered lungs and with increasing multicellularity. D) Archetypes of engineered endothelium. E) Aqp1, Kdr, Mmrn1, and Atf3 expression defining archetypes in engineered endothelium. F) Changes in relevant endothelial gene expression in engineered lungs. G) Archetypes of engineered mesenchyme. H) Pdgfra, Tgfbr3, Gucy1b1, and Notch3 expression defining archetypes in engineered mesenchyme. I) Changes in relevant mesenchymal gene expression in engineered lungs. J) Archetypes of engineered macrophages. K) Cxcl2, S100a9, Naaa, and C1qb expression defining archetypes in engineered macrophages. L) Changes in relevant macrophage gene expression in engineered lungs. See also Figures S1-S4.

### Alveolar macrophages support essential signaling for tissue self-organization

We performed ligand-receptor expression analysis using NICHES to capture cell-cell connectivity at single-interaction resolution^22^ (Fig. 4A). To measure the effects of adding alveolar macrophages on signaling in the system, we compared tri- and quad-culture conditions (Fig. 4B). In the epithelial niche, several mechanisms associated with development and stem cell regulation were increased with the addition of alveolar macrophages (Fig. 4C). This included *Wnt7a*—*Lrp6*, *Ereg*— *Egfr*, *Fgf2*—*Fgfr3*, and *Rgmb*—*Bmpr1b*. In the endothelial niche, mechanisms promoting microvascular maturation and stabilization were increased in quad-cultures, including *Dll4*— *Notch3*, *Sema3e*—*Plxnd1*, *Ebi3*—*Il27ra*, and *Mdk*—*Lrp2* (Fig. 4D). The immune niche demonstrated a pattern where certain inflammatory ligands were being expressed but not received in tri-culture lungs, essentially “orphan ligands”, signals which were then received by immune cells expressing the receptor in quad-cultures (Fig. 4E). *Cxcl6*—*Cxcr1* and *Il34*—*Csf1r* followed this type of pattern. In other instances, immune cells expressed a novel ligand, such as via *Il18*—*Cd48* and *Tnfsf13*—*Fas*. For additional details of NICHES analysis, see Supp. Fig. 5 and 6.

**Figure 4.**
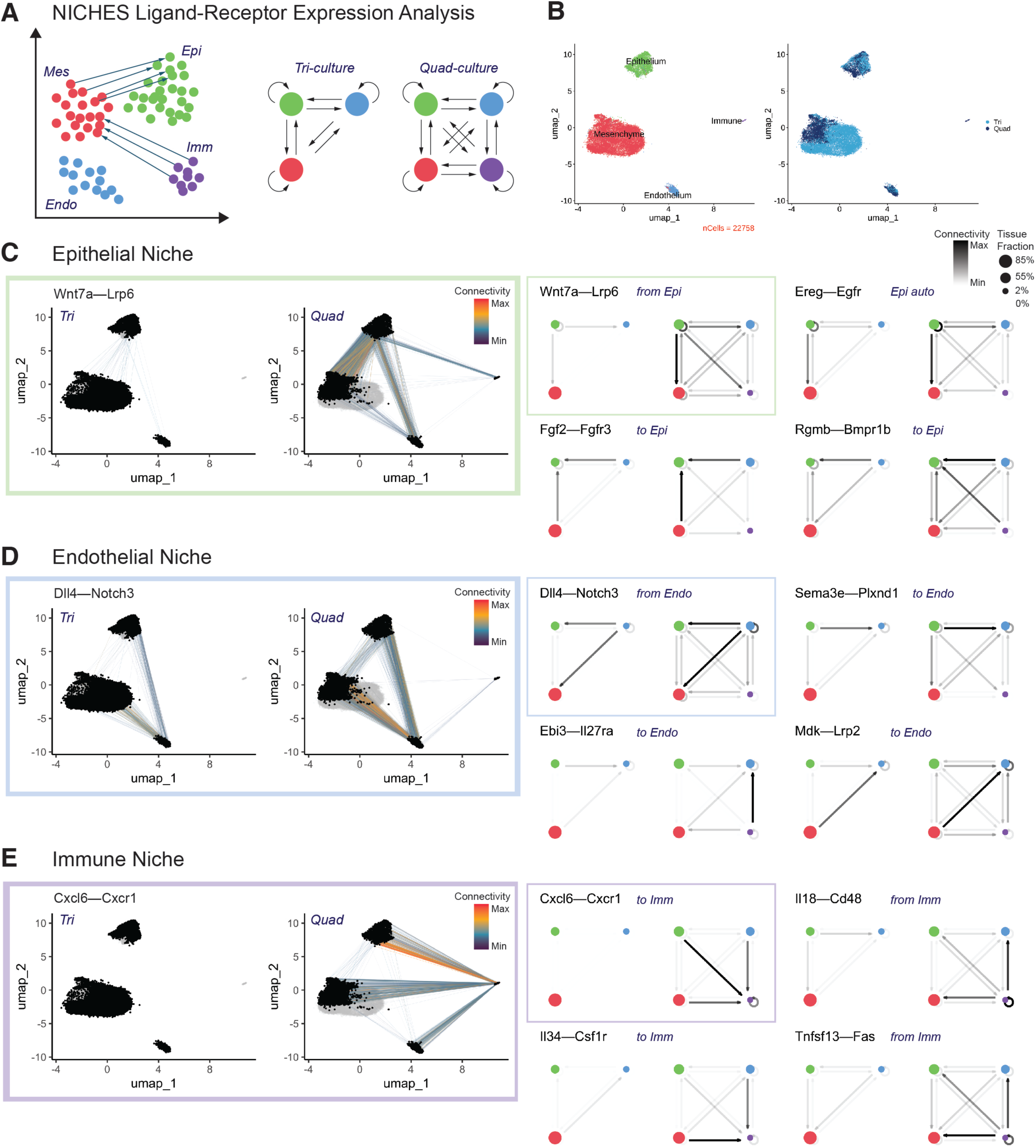
Alveolar macrophages support essential signaling for tissue self-organization. A) NICHES measures potential ligand-receptor interactions at single-cellular resolution 22, which was then applied to compare cell-cell signaling patterns in tri- and quad-culture engineered lungs. B) Embedding tri- and quad-culture lungs in the same phase space for analysis. C) Significant variation in epithelial niche signaling, including Wnt7a—Lrp6, Ereg—Egfr, Fgf2—Fgfr3, and Rgmb—Bmpr1b. D) Significant variation in endothelial niche signaling, including Dll4—Notch3, Sema3e—Plxnd1, Ebi3—Il27ra, and Mdk—Lrp2. E) Significant variation in immune niche signaling, including Cxcl6—Cxcr1, Il18—Cd48, Il34—Csf1r, and Tnfsf13—Fas. See also Figures S5-S6.

### Engineered tissue model demonstrates evidence of functional delegation

Functional delegation is a principle whereby the division of labor between cell types results in the delegation of some functions from highly specialized “client” cells (e.g., epithelial cells or neurons) to “accessory” cells (e.g., stromal and immune cells)^23^. The engineered lung model system allowed us to study this concept by comparing gene expression and cell-cell signaling in tissues with and without alveolar macrophages. By analyzing genes and mechanisms that were differentially expressed in a single class in tri-culture lungs, that are then differentially expressed in the immune cells of quad-culture lungs, we identified possible vestiges of evolutionary functional delegation to alveolar macrophages (Fig. 5A). We found that genes following this expression pattern grouped into functional categories associated with lipid metabolism and endocytosis (Fig. 5B). As these genes “moved” in top expression from one cell class to immune cells, most also increased in relative expression and specificity in immune cells compared to their top expressing class in tri-culture (Fig. 5C-F). At the cell-cell signaling level, one possible pattern of functional delegation could be immune cells stepping in to provide a needed ligand to the rest of the system (Fig. 5G). Pulling this pattern identified ligand expression specifically delegated to immune cells, including in *Lpl*—*Lrp2*, *Igf1*—*Insr*, and *Hgf*—*Met* pathways (Fig. 5H). Another pattern of functional delegation could be immune cells arriving to receive existing signals from the rest of the system (Fig. 5I). Investigating this pattern yielded several known inflammatory pathways, such as *Il6*—*Il6r*, *Il10*—*Il10rb*, and *Il24*—*Il20rb* (Fig. 5J).

**Figure 5.**
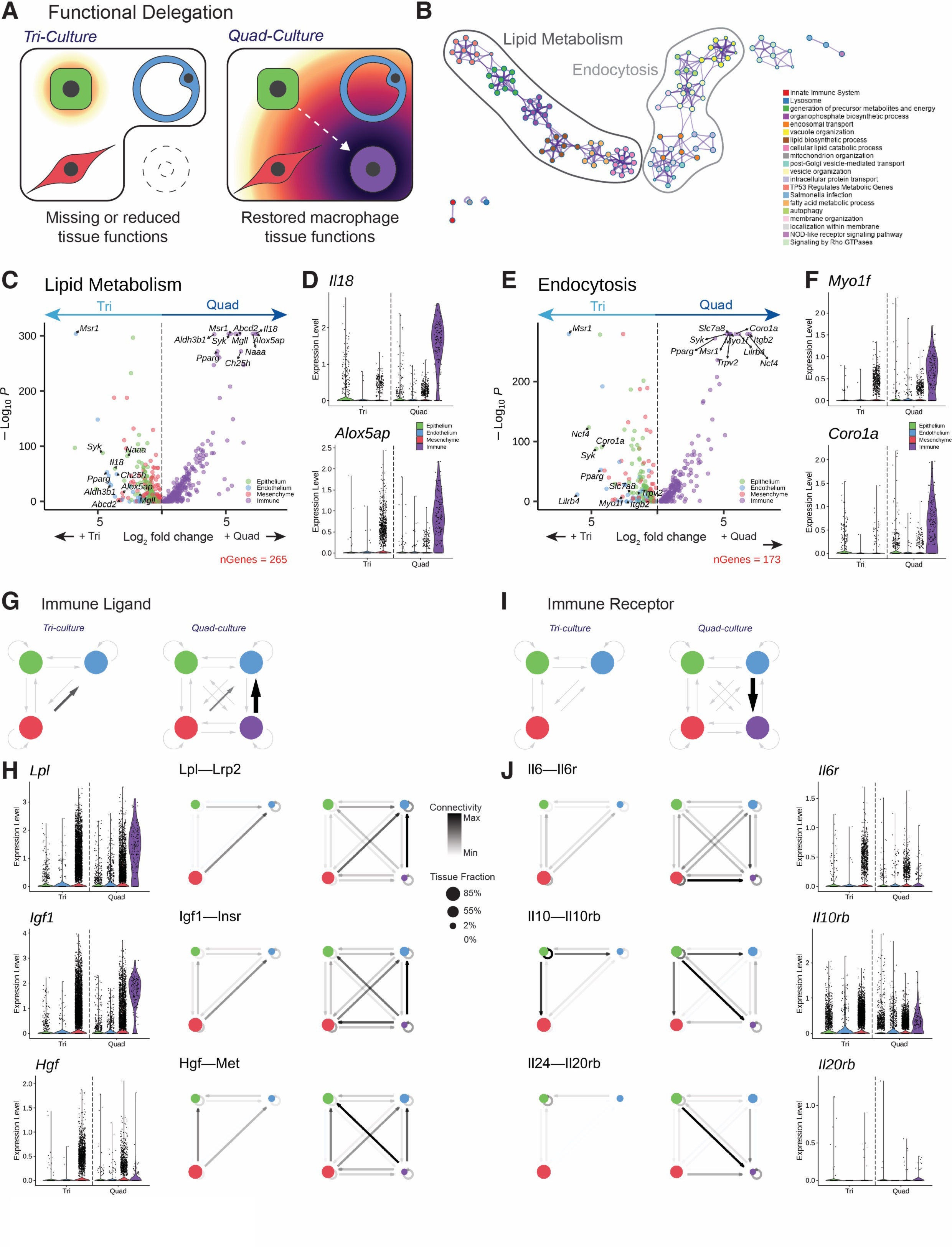
Engineered tissue model demonstrates evidence of functional delegation. A) Predicted gene expression pattern indicating functional delegation between tri- and quad-culture lungs. B) Groupings of “moved” genes indicating the functional delegation of lipid metabolism and endocytosis to immune cells. C) Modified volcano plot comparing expression of “moved” genes associated with lipid metabolism in tri- and quad-culture lungs. D) Violin plots of Il18 and Alox5ap expression between tri- and quad-culture lungs divided by class, demonstrating how these genes “moved” between conditions. E) Modified volcano plot comparing expression of “moved” genes associated with endocytosis in tri- and quad-culture lungs. F) Violin plots of Myo1f and Coro1a expression between tri- and quad-culture lungs divided by class, demonstrating how these genes “moved” between conditions. G) Predicted pattern of mechanism activation where immune cells provide a ligand more strongly in quad-culture lungs. H) Ligand and mechanism expression following this pattern, including Lpl—Lrp2, Igf1—Insr, and Hgf—Met. I) Predicted pattern of mechanism activation where immune cells provide a novel receptor in quad-culture lungs. J) Mechanism and receptor expression following this pattern, including Il6—Il6r, Il10—Il10rb, and Il24—Il20rb.

### Alveolar barrier function in quad-culture lungs in vitro and in vivo

Engineered lungs were evaluated for alveolar barrier function, including recapitulation of a functional mechanical barrier between air and blood compartments, as well as the potential for cellular alveoli to perform gas exchange *in vivo*. Fluid pressures measured continuously over the course of culture were extrapolated to flows in the system^24^, demonstrating a significant 668% increase in venous outflow in quad-culture lungs (Fig. 6A). By modeling microvascular flows from measured pressures^25^, quad-culture lungs displayed a significant 84% increase in vascular recruitment (Fig. 6B). Increasingly physiologic flow patterns in the tissue indicate enhanced organization and stability of the blood-gas barrier with the addition of alveolar macrophages (Fig. 6A,B). We visualized this barrier by TEM, which showed alveolar thickness approaching native values (900 nm^26^) (Fig. 6C). Gene expression associated with cell-cell junctions in engineered epithelium and endothelium increased 577% and 1050% respectively, in quad-culture lungs over starting cells (Fig. 6D-E). To benchmark these constructs to previous studies, we evaluated quad-culture engineered lungs *in vivo* by left lung pneumonectomy transplant. These lungs displayed functional blood-gas barrier in the short term (1 hour), including functional capillary barrier and evidence of potential gas exchange (Fig. 6F,G).

**Figure 6.**
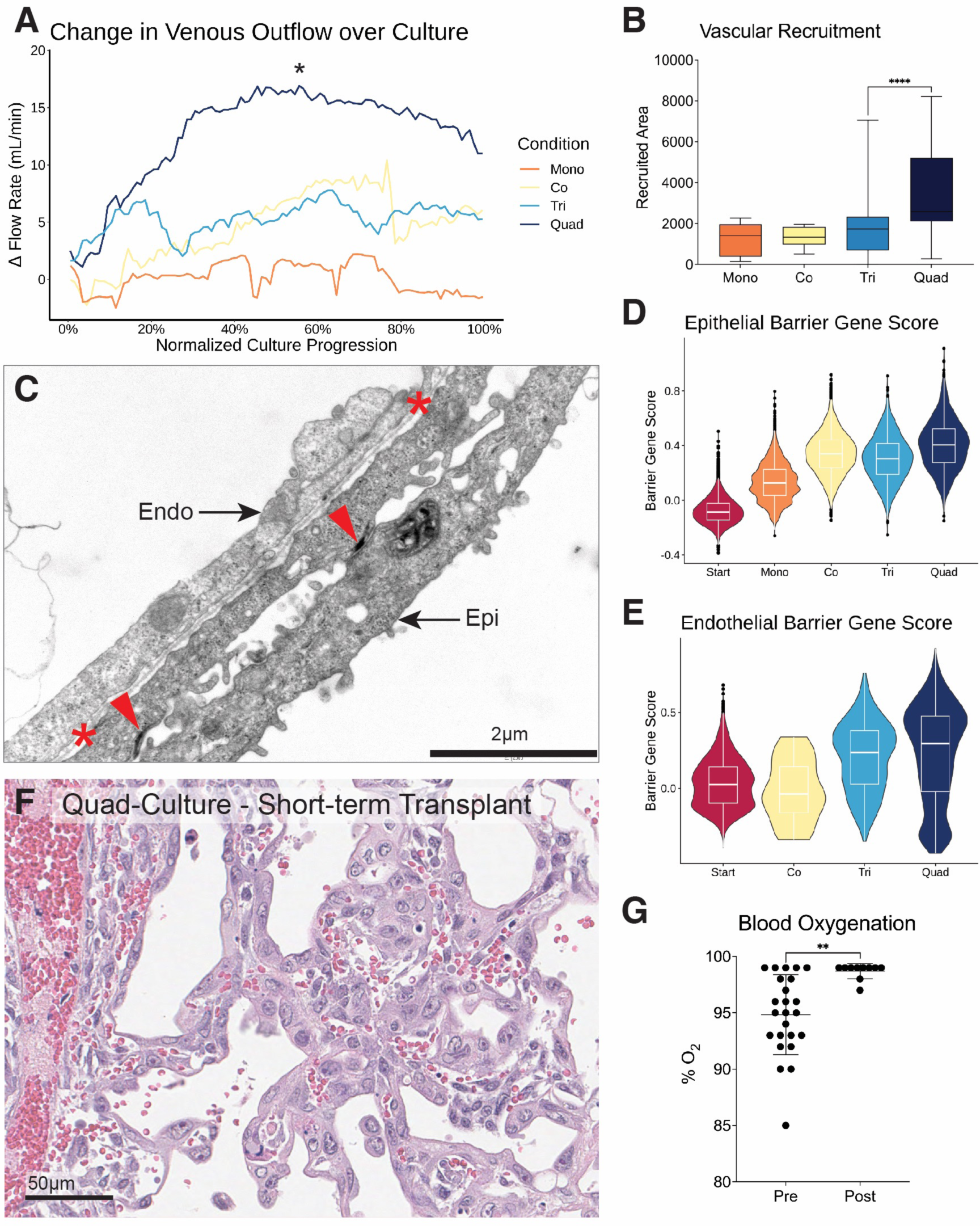
Alveolar barrier function in quad-culture lungs in vitro and in vivo. A) Change in measured venous outflow over the course of culture across each engineered lung condition (*p < 0.05). B) Measured vascular recruitment over the course of culture across each engineered lung condition (****p < 0.0001). C) TEM of engineered lung demonstrating alveolar barrier approaching native thickness (900nm); basement membrane labeled between *, and arrowheads indicate epithelial tight junctions. D) Epithelial and E) endothelial gene expression scores by scRNAseq between conditions. F) H&E of blood in large vessels and capillaries with little leakage into the air compartment in a short-term transplant of quad-culture lungs in vivo. G) Measured blood oxygenation by pulse oximeter during engineered left lung pneumonectomy transplant; “Pre” = left lung clamped off, “Post” = engineered left lung unclamped to vascular perfusion; significant increase in blood oxygenation indicates possible gas exchange by engineered lung in vivo (**p < 0.005).

## Discussion

Here we present an engineered whole-lung tissue model unprecedented in cell coverage, viability, biomimicry, function, and methodologic replicability. We leveraged this model system to study principles of tissue biology, including self-organization, constructive regeneration, and functional delegation^23,27^. Engineered lungs are the most biomimetic *in vitro* system available to explore these concepts given the ability to culture cells in a native ECM environment under physiologic mechanical conditions^13^. In this system we can control many culture parameters, such as source cells, metabolic input, and physical stimuli, which are difficult or impossible to modulate *in vivo*. Our study of cellular communities in regeneration was only possible by using such an *in vitro* model system.

The most prominent finding of this work is the central role of alveolar macrophages in orchestrating tissue-wide regeneration. This is evidenced by histology, cell phenotype, cell-cell signaling, and mechanical measurements of tissue function. The addition of alveolar macrophages precipitated a stigmergic shift from a disorganized to organized tissue state^27,28^, which coincided with an upregulation of known developmental and regenerative signaling pathways. These patterns of cellular differentiation and signaling in quad-cultures comprise a landscape of “constructive” regeneration in the lung. Disease states, such as fibrosis and cancer, are often characterized as the failure of normal regenerative processes and could be described as “frustrated” regeneration^29-31^. Interestingly, several significant mechanisms in our analyses, such as those involving *Itgb6* and *Cxcr1*, have been implicated in disease states^32,33^. However, since our quad-culture tissues do not display other hallmarks of disease, we believe our engineered tissues exemplify the activity of these pathways in their constructive state. This provides an interesting and potentially useful counterpoint to clinical data, and future work could be devoted to perturbing such marker pathways in engineered tissues to produce a new class of *in vitro* disease models.

The inflammatory state of alveolar macrophages underwent a distinct shift when introduced to engineered lung cultures, likely contributing to their regenerative effects. Our NICHES analysis demonstrated that this cell state introduced new direct inflammatory communication to the system, through novel expression of ligands and receptors in immune cells. In other cases, the presence of alveolar macrophages had indirect effects on the system, causing other populations to change how they communicated with each other. This demonstrates how expression of most genes and mechanisms are interdependent with other parts of the system and adding or removing a cell type drastically shifts the entire communication landscape.

This type of systemic shift is exemplified by patterns of functional delegation we observed going from tri- to quad-culture lungs. Functional delegation is best studied in this biomimetic *in vitro* system for its complete flexibility in input cellular communities, including conditions that would This type of systemic shift is exemplified by patterns of functional delegation we observed going from tri- to quad-culture lungs. Functional delegation is best studied in this biomimetic *in vitro* system for its complete flexibility in input cellular communities, including conditions that would not be consistent with life in an animal model. If a client cell with a specialized tissue-level function can delegate some of its non-cell-autonomous functions to an accessory cell, the client cell can optimize the performance of its primary specialized functions^23^. This explains why quad- culture tissues functioned better as a whole, with each cell type able to play its respective role. Our results suggest that lipid metabolism and endocytosis are two supportive functions which were historically delegated to alveolar macrophages from other cell types in the lung. Other cells still possess the machinery to perform these functions to a degree in their absence, though at this point alveolar macrophages are known specialists of lipid handling and endocytosis in lung*^34^*. These functions may have been delegated by repurposing the machinery of canonical immune functions, such as recognizing and engulfing pathogens. In contrast, most inflammatory functions of alveolar macrophages were completely absent in tri-culture lungs, suggesting the inflammatory role of macrophages is more specialized and may have split off from the original client cell earlier in multicellular diversification. Further exploration is required to determine which aspects of endocytosis and lipid handling have been delegated to alveolar macrophages, as well as methods for estimating the cellular phylogenic tree of functional delegation.

One of the goals of this work was to achieve functional blood-gas barrier in engineered lungs. We were able to reach about 50% venous outflow in quad-culture lungs, representing a 668% improvement in venous outflow over epithelial mono-cultures. Imperfect mechanical barrier does not limit the model’s utility *in vitro*, but further improvements would be required for the tissue to perform well long-term *in vivo*. Future work to improve barrier in engineered lungs could include seeding additional cell types to support barrier formation and stability, as well as developing a protocol of growth factor delivery to more actively direct cellular differentiation and organization.

We have shown that engineered lungs serve as a potentially valuable model system for investigating questions in biology and disease. Our work highlights the importance of cellular communities in tissue organization, particularly the central role of alveolar macrophages in cell-cell signaling during tissue regeneration. Furthermore, we have illustrated how multicellularity in engineered tissues contributes to tissue-level function. This platform and these methodologies could offer a robust foundation for anyone performing *in vitro* modeling studies in the future.

## Acknowledgements

This work was supported by grants from the National Institutes of Health: F32HL162428 (A.M.G.), F30HL143906 (M.S.B.R.), T32GM086287 (M.S.B.R.), T32GM136651 (M.S.B.R. and K.L.), F30HL143880 (K.L.), and R00HL159261 (Y.Y.). The opinions expressed are those of the authors and do not necessarily represent the thoughts or opinions of NHLBI, NIGMS, NIH, or the United States government. This work was also supported by an unrestricted research gift from Humacyte Inc. (L.E.N.). We thank the Yale Center for Genome Analysis (YCGA), particularly Christopher Castaldi; Medicine Center for Cellular and Molecular Imaging (CCMI) Electron Microscopy Facility, particularly Dr. Xinran Liu; the Yale Stem Cell Center Genomics Core, particularly Dr. Mei Zhong; the Yale Scientific Glassblowing Laboratory, particularly Daryl Smith and Preston Smith; and the Yale Animal Resources Center (YARC). We thank the following people for useful discussions that have shaped this project: Dr. Themis Kyriakides, Dr. Jay Humphrey, Dr. Marie Egan, Dr. Daniel Greif, Dr. Andre Levchenko, Dr. Yuval Kluger, Junchen Yang, Dr. Lisa Leffert, Dr. Helene Benveniste, Dr. Robert Schonberger, and Dr. Carlos Fernandez-Hernando.

## Methods

### Lung Decellularization

All animal procedures in this study were approved by the Yale University Institutional Animal Care and Use Committee and complied with the NIH Guidelines for the Care and Use of Laboratory Animals. Male WT Sprague Dawley rats (*Rattus norvegicus*) were obtained from Charles River (8 weeks old, 300g) and used for lung decellularization. Decellularization of whole adult rat lungs was performed using an established protocol^9^. Detergent and enzyme reagents include triton X-100 (Sigma), benzonase nuclease (Sigma), and sodium deoxycholate (SDC, Sigma).

### Starting cell culture

Male WT Sprague Dawley rats (*Rattus norvegicus*) were obtained from Charles River (8-12 weeks old, 300-350 g) and used for isolation of peBC and BAL. All cell populations used in this study were primary cells derived from Sprague-Dawley rat lungs. Pharmacologically expanded basal cells (peBC) were isolated and cultured as previously described^13^. peBC were isolated into and expanded in EpiX medium (Propagenix) on collagen-coated (Advanced Biomatrix) tissue culture flasks. Primary rat lung microvascular endothelial cells (RLMVECs) were purchased and expanded in MCDB-131 complete medium (VEC Technologies) on fibronectin-coated (Sigma) tissue culture flasks. Neonatal rat lung fibroblasts were isolated and cultured as previously described^14^. Lung fibroblasts were isolated from 7-9-day-old WT Sprague Dawley rat pups where pups were selected randomly without bias for males or females. Fibroblasts were expanded in DMEM-HG (Gibco) + 10% FBS. Alveolar macrophages were isolated from adult rats by bronchoalveolar lavage (BAL) as previously described^35^. Bulk isolates were rinsed briefly in antibiotics/antimycotics (10% pen-strep, 4% amphotericin-B, 2% gentamycin) prior to direct seeding into engineered lungs.

### Engineered Lung Seeding

Decellularized lungs were mounted in a custom glass bioreactor for culture (Yale Scientific Glassblowing Laboratory). Scaffolds were connected to the system via cannulated pulmonary artery, pulmonary vein, and trachea. 100 million endothelial cells were seeded into the pulmonary artery and vein (50 million each side) as previously described^36^. Epithelium (40 million cells), mesenchyme (5 million), and immune cells (8 million) were seeded into the air compartment as previously described^13^.

### Engineered Lung Culture

Lungs were then cultured for seven days by arterial medium perfusion at 40mL/min and applied cyclic stretch resembling breathing by pulling negative pressure within the bioreactor with the trachea exposed to atmospheric pressure. Culture medium was composed of 50:50 growth factor-rich epithelial and endothelial media, Propagenix Airway Differentiation Medium (PADM, Propagenix) and MCDB-131 without serum (VEC Technologies), respectively. Epithelial mono-culture lungs only contained PADM. Culture medium was tested for glucose, lactate, and pH daily, and continuous medium exchange rates were titrated to maintain acceptable levels of metabolites. Additionally, growth factor-rich medium was titrated to DMEM-LG (Gibco) over the course of culture to discourage cellular dependence on exogenous growth factors and encourage cell-cell support. Pressures were measured at all inlets and outlets of the organ for the duration of culture as previously described^24^, which allowed for subsequent mechanical barrier measurements.

### Engineered Lung Histology

Engineered lungs were fixed by vascular perfusion of formalin for 30 minutes and static incubation at room temperature overnight. Paraffin-embedding, sectioning, H&E staining, and slide scanning was performed by Yale Pathology Tissue Services (YPTS). Slide scans were viewed and imaged using QuPath 0.5.1.

### Engineered Lung Dissociation

Engineered lungs were dissociated to single-cell suspensions as previously described ^37^. Efficient temperature control and minimization of time from tissue to single-cell suspension was critical in this protocol to maximize cell viability and downstream data quality. Briefly, enzyme solution was made using collagenase/dispase (1 mg/ml; Roche), elastase (3 U/ml; Worthington), and deoxyribonuclease (DNase) (20 U/ml; Worthington) in DMEM, and preheated to 37°C for use. Upon bioreactor take-down, the lower right lobe of the engineered lung was tied off and removed to a clean petri dish for singe-cell enzymatic dissociation. 25ml warm enzyme solution was repeatedly instilled into the tissue using a syringe with a 25G needle, taking care to saturate the entire tissue with enzyme. Tissue and enzyme effluent were pooled in a 50ml conical and incubated in a 37°C water bath with shaking at 40rpm for 20 minutes. Reagents for 10x GEM preparation were removed from the freezer during enzyme incubation. The softened tissue was then forced through a clean wire strainer to remove large collagenous structures, the strainer was rinsed with 20ml ice-cold DMEM containing 10% fetal bovine serum (FBS), and pooled with strained cells. Cell solution was centrifuged at 300g and 4°C for 5 minutes. Pellet was resuspended in 10ml 0.01% bovine serum albumin (BSA) in PBS without pipetting and spun down a second time. Pellet was resuspended in 10ml 0.01% BSA in PBS, passed through a 70μm cell strainer (Falcon), and centrifuged at 300g and 4°C for 3 minutes. Pellet was resuspended in 10ml 0.01% BSA in PBS and passed through two sequential 40μm cell strainers (Falcon). Cells were then counted and diluted in 0.01% BSA in PBS to a concentration of 1 million cells/ml for GEM formation. Starting cell suspensions were also prepared at 1 million cells/ml in 0.01% BSA in PBS for GEM formation and sequencing.

### scRNAseq library preparation, sequencing & alignment

Libraries were generated for single-cell RNA sequencing (scRNAseq) using Chromium Single Cell 3’ v2, v3, and v3.1 (10x Genomics). Libraries were sequenced by the Yale Center for Genomic Analysis (YCGA) at a target recovery of 5000-10000 cells per sample, and a target depth of 50,000 reads/cell on an Illumina Hiseq4000 or NovaSeq 6000. Cutadapt 1.17 was used to preprocess FASTQ files as necessary ^38^. Data was aligned and extracted using STARsolo (STAR-2.7.3a) ^39^ with appropriate barcode whitelists and parameters for each chemistry. Ensembl Rnor6.0 release 92 was used to align the data. The top 30,000 cell barcodes per sample, ranked by nUMI, were imported into R for further processing.

### scRNAseq data cleaning

scRNAseq data was cleaned following established best practices^40^. First, data were loaded into Seurat (4.1.1)^41^ and filter cutoffs were manually selected for nCount_RNA, nFeature_RNA, and percent mitochondrial reads for each sample. Samples were then carried through to an initial round of embedding and clustering to ensure expected cell populations were still present post-filtration. Samples were then manually cleaned of partial cells and doublets through iterative embedding and cluster removal. Where possible, to improve statistical power in identifying clusters to be removed, samples similar in chemistry and cell type diversity were combined by ComBat^42^ or Seurat CCA integration^43^. Clusters containing low-information cells were removed from individual and combined samples. Final cluster-based cleaning was performed on individual samples. Cell class was assigned by cluster-based expression of lineage markers *Epcam*, *Cdh5*, *Col1a1*, or *Ptprc*, which were found to be highly specific to epithelium, endothelium, mesenchyme, and immune populations, respectively. High-information cells co-expressing multiple lineage markers were deemed likely doublets and removed. For data cleaning code, see our Github repository. For quality control plots demonstrating the data cleaning process, see Document S1.

### scRNAseq phenotype analysis

Samples were divided by cell class and class objects integrated by sample (Seurat CCA)^43^. Class objects were embedded and clustered to show relevant archetype segregation. Analysis packages include Slingshot (2.2.1)^44^, Velocyto-R (0.6)^45^, Complex Heatmap (2.10.0)^46^, and Gene Ontology^47,48^. For code to replicate phenotype analysis or figure plots, see our Github repository.

### scRNAseq NICHES analysis

Ligand-receptor expression analysis was performed at single-cell resolution using the R package, NICHES (1.0.0)^22^. Differential expression analysis was performed to identify mechanisms that varied significantly between tri- and quad-culture lungs. For code to replicate NICHES analysis or figure plots, see our Github repository.

### scRNAseq functional delegation analysis

Differential expression lists were generated to identify potentially delegated genes and mechanisms. Differential gene expression was performed in tri-culture and quad-culture lungs independently, by cell class. Genes were identified as “moved” if they were top-expressed in different classes between tri- and quad-culture lungs. The gene list was further filtered to identify genes that were top expressed in immune cells in quad-culture lungs, or those that may have been “delegated” from other cell types. A similar method was performed for identifying signaling mechanisms of interest. Mechanisms that were highly expressed in both tri- and quad-culture lungs were identified using NICHES data. Mechanisms were then identified as “moved” to or from immune cells if the ligand or receptor were expressed by the immune class in quad-culture lungs. Results were plotted using Enhanced Volcano (1.12.0) or NICHES. For code to replicate functional delegation analysis or figure plots, see our Github repository.

### Alveolar barrier measurements

Pressures were measured at the inlets and outlets of the engineered organ for the duration of culture. These pressures could then be extrapolated to flows, resistances, and barrier and vascular recruitment measurements in the system as previously described^24,25^. TEM of engineered lungs was performed by the Yale School of Medicine Center for Cellular and Molecular Imaging (CCMI) Electron Microscopy Facility.

### Transplant surgery

Left-lung transplant of quad-culture engineered lungs was performed on adult male rats (n=4) as previously described^49-51^. In brief, a healthy, adult male rat between 9-14 weeks of age was anesthetized, intubated, and placed on continuous inhaled anesthesia with blood pressure, O2, and temperature monitoring within a sterile surgical field. The animal was given a dose of bupivacaine and buprenorphine. The engineered lung was removed from the bioreactor in sterile fashion, instilled with 100ul of Curosurf (Chiesi) into the airways, and attached to a rodent ventilator set to pressure-driven ventilation. The left lobe was dissected and prepared with cuffs for surgical anastomosis^49^. A left thoracotomy was then performed and the engineered lung was implanted as described^49^, with the PV, PA, and bronchus unclamped in this order. Observations were made of organ reperfusion, airway patency, and systemic oxygen saturation. A custom-fabricated silicone chest tube was inserted and placed on regulated low vacuum. The chest wall and skin were then closed layer-by-layer. The animal was given a dose of meloxicam, switched to room air, and allowed to passively recover until self-extubation. The animal was monitored continuously until sacrifice, with buprenorphine administered every 6-12 hours and meloxicam administered every 24 hours.

---

**Figure S1.**
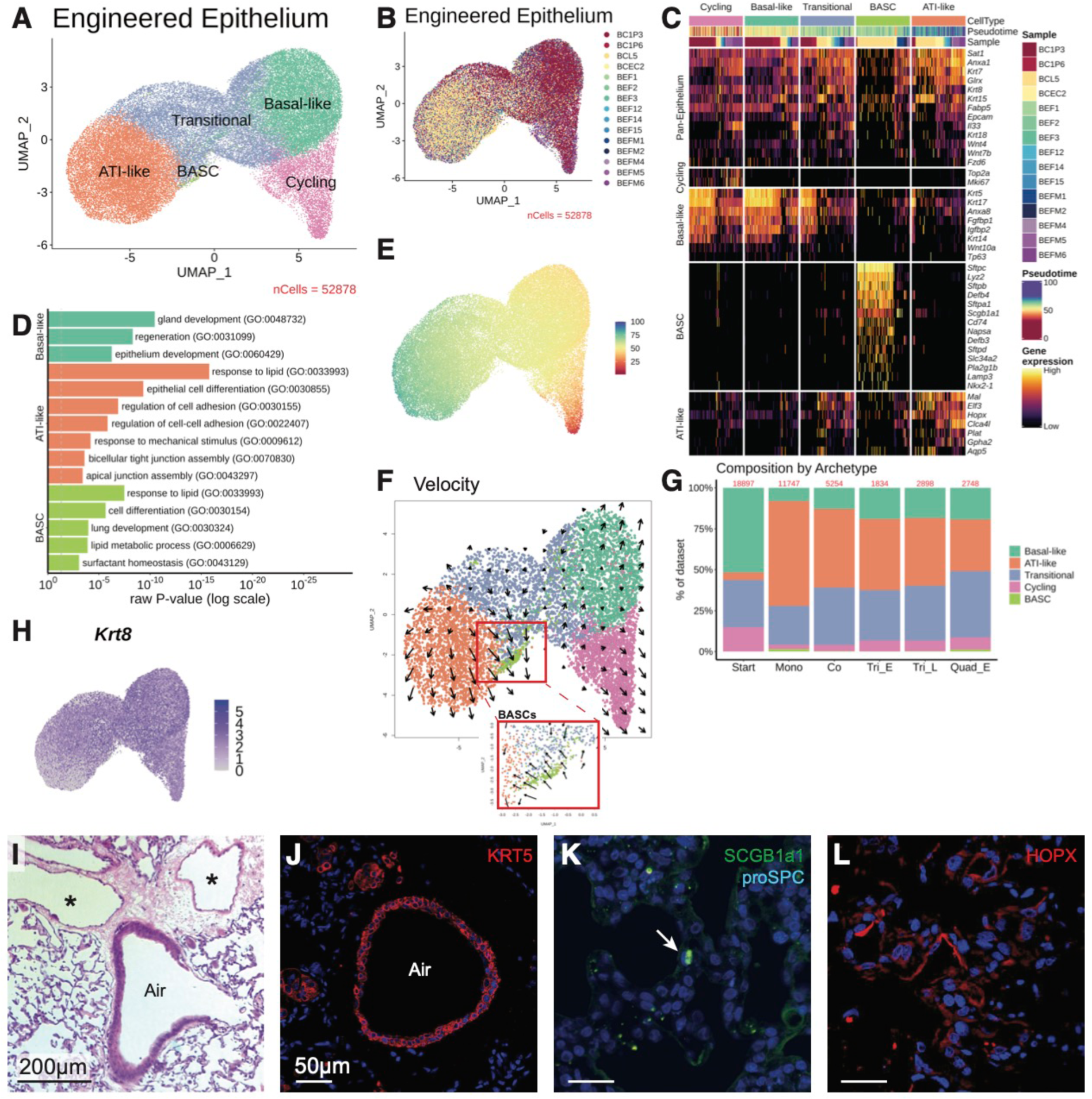
Epithelium differentiates into ATI-like barrier-forming cells and BASCs, related to Figure 3. A) UMAP of engineered epithelial archetypes. B) UMAP of integrated epithelium by sample. C) Heatmap of defining genes of engineered epithelial archetypes, normalized by gene across archetypes. D) Top GO terms of DEGs defining select archetypes, plotted by significance. E) UMAP of engineered epithelium colored by pseudotime (Slingshot). F) UMAP of engineered epithelium annotated by RNA velocity (Velocyto). G) Plot of cellular proportions by archetype across datasets; number of cells per dataset at top of columns in red. H) Feature plot of Krt8 expression. I) H&E of engineered lung demonstrating airway (“Air”) and alveolar repopulation, and vascular repopulation (* indicates large vessels). IF staining of engineered lung for J) KRT5 in the airway, K) a proSPC+/SCGB1A1+ BASC (arrow), and L) HOPX in alveoli.

**Figure S2.**
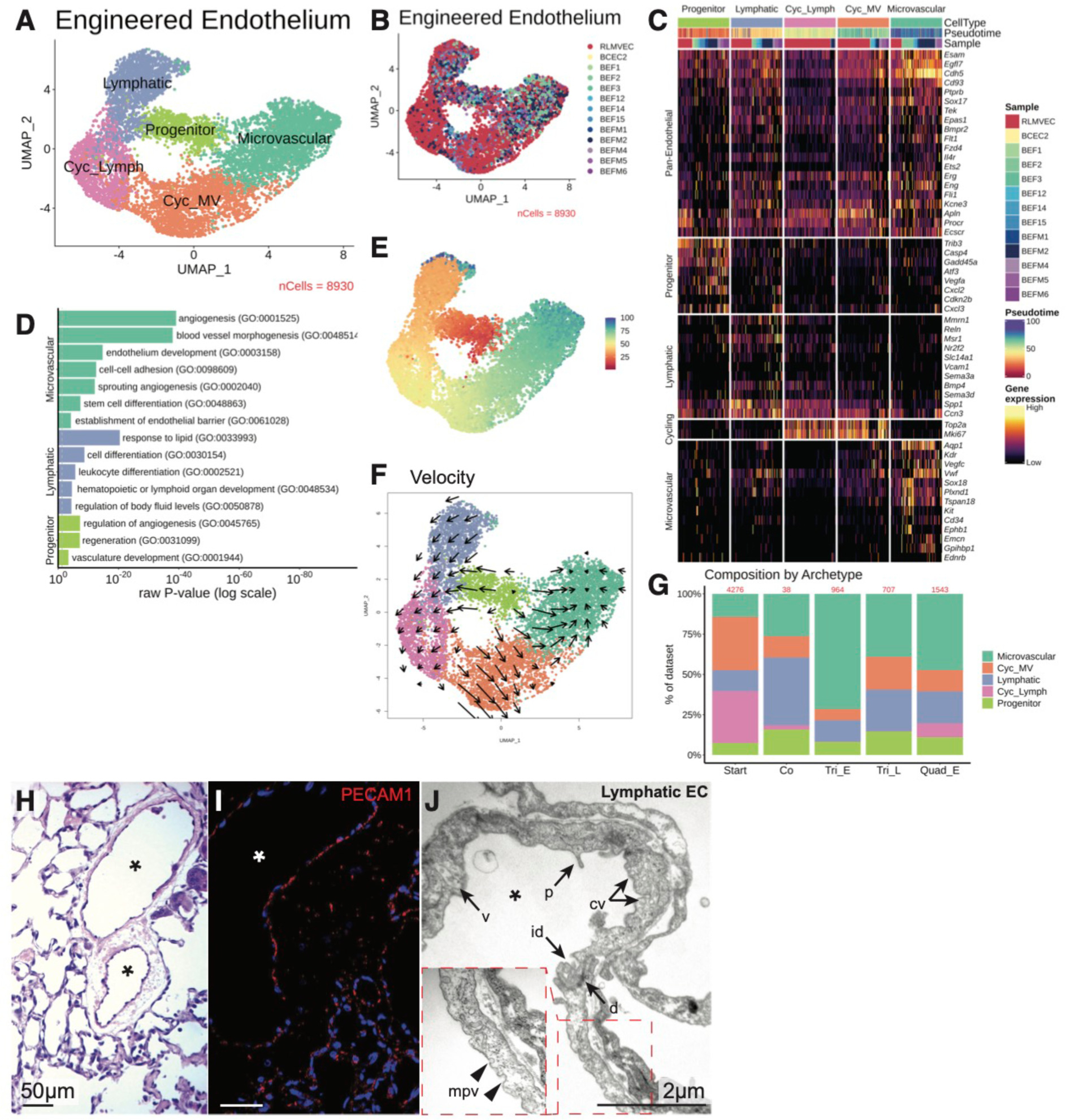
Endothelium adopts lymphatic and microvascular character, related to Figure 3. A) UMAP of engineered endothelial archetypes. B) UMAP of integrated endotheium by sample. C) Heatmap of defining genes of engineered endothelial archetypes, normalized by gene across archetypes. D) Top GO terms of DEGs defining select archetypes, plotted by significance. E) UMAP of engineered endothelium colored by pseudotime (Slingshot). F) UMAP of engineered endothelium annotated by RNA velocity (Velocyto). G) Plot of cellular proportions by archetype across datasets; number of cells per dataset at top of columns in red. H) H&E of engineered lung demonstrating vascular repopulation (* indicates large vessels). I) IF staining of engineered lung for PECAM1+ endothelium in large vessels and capillaries. J) TEM of lymphatic endothelial cells in engineered lung, as identified by ultrastructural features.

**Figure S3.**
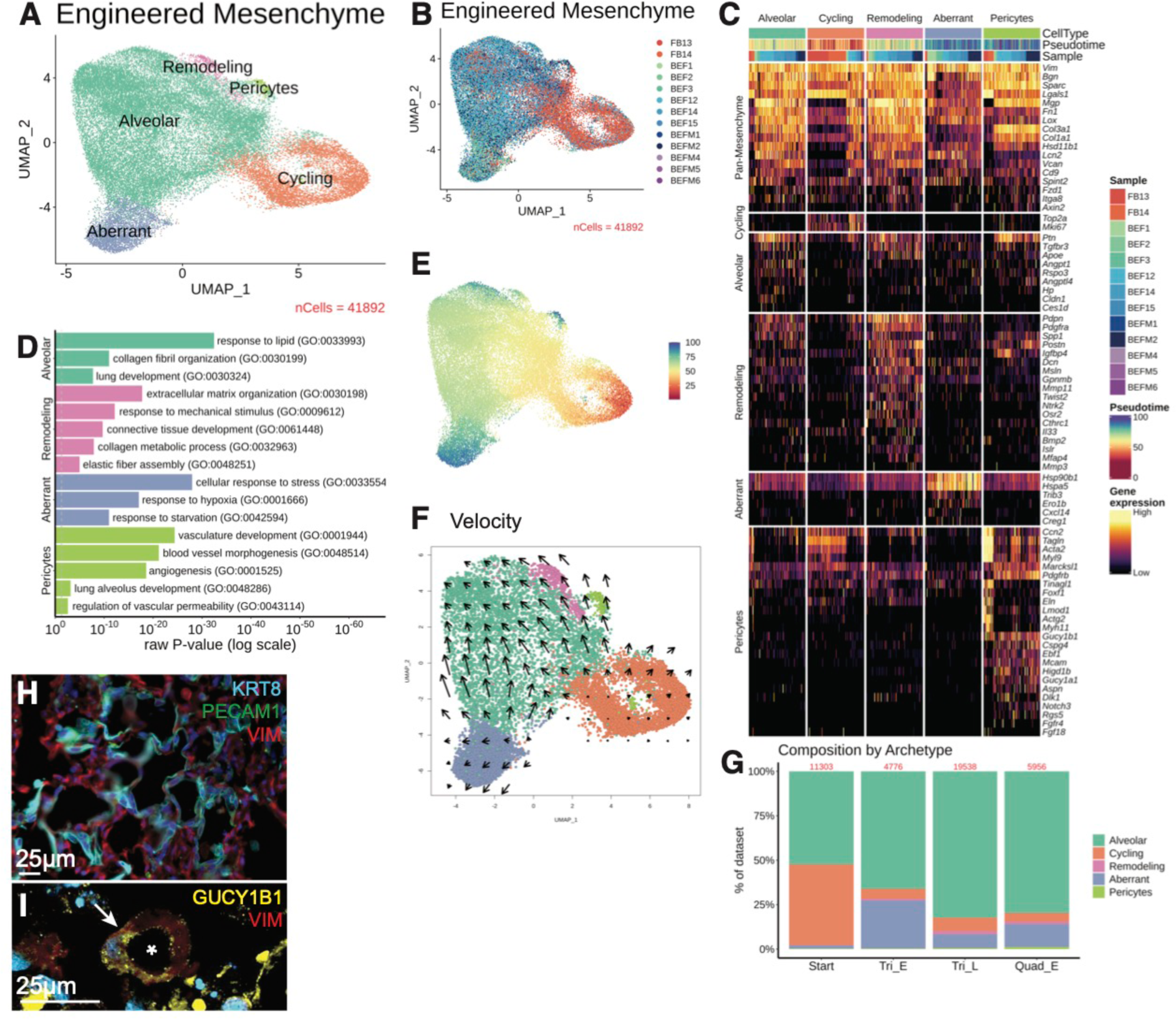
Mesenchyme differentiates into pericytes and supports alveolar formation in the presence of alveolar macrophages, related to Figure 3. A) UMAP of engineered mesenchymal archetypes. B) UMAP of integrated mesenchyme by sample. C) Heatmap of defining genes of engineered mesenchymal archetypes, normalized by gene across archetypes. D) Top GO terms of DEGs defining select archetypes, plotted by significance. E) UMAP of engineered mesenchyme colored by pseudotime (Slingshot). F) UMAP of engineered mesenchyme annotated by RNA velocity (Velocyto). G) Plot of cellular proportions by archetype across datasets; number of cells per dataset at top of columns in red. H) IF staining of engineered tri-culture lung for KRT8+ epithelium, PECAM1+ endothelium, and VIM+ mesenchyme demonstrating fibroblast overgrowth. I) IF staining in engineered lung of a GUCY1B1+/VIM+ pericyte (arrow, * indicates capillary lumen).

**Figure S4.**
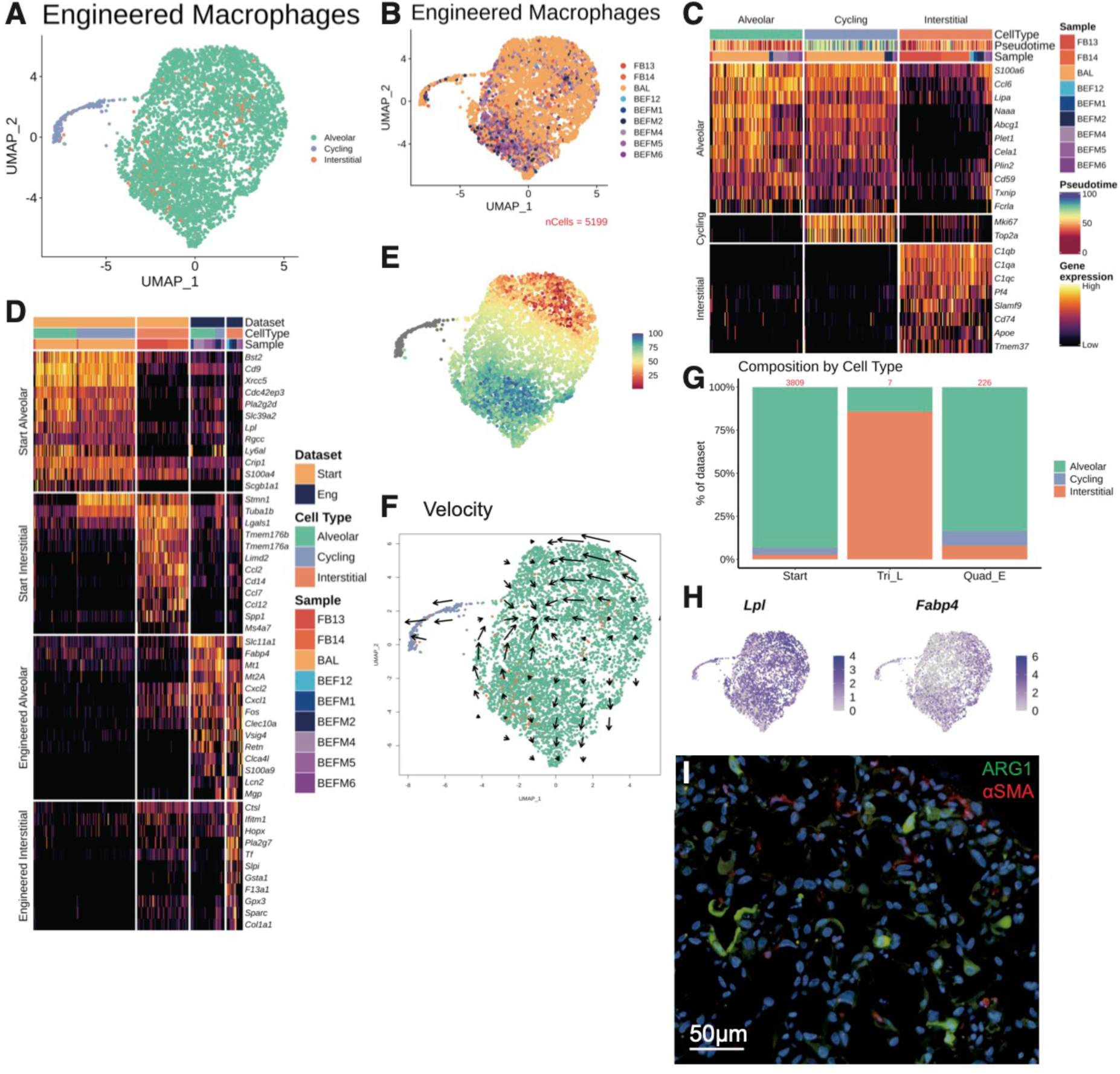
Alveolar macrophages activate and promote constructive tissue organization in the engineered alveolus, related to Figure 3. A) UMAP of engineered macrophage types. B) UMAP of integrated macrophages by sample. C) Heatmap of defining genes of engineered macrophage types, normalized by gene across types. D) Heatmap of activation markers in engineered macrophages before seeding (‘Start’) and post-culture (‘Eng’), normalized by gene across types. E) UMAP of engineered macrophages colored by pseudotime (Slingshot). F) UMAP of engineered macrophages annotated by RNA velocity (Velocyto). G) Plot of cellular proportions by type across datasets; number of cells per dataset at top of columns in red. H) Feature plot of Lpl and Fabp4 expression. I) IF staining of engineered lung for ARG1+ alveolar macrophages in the alveoli.

**Figure S5.**
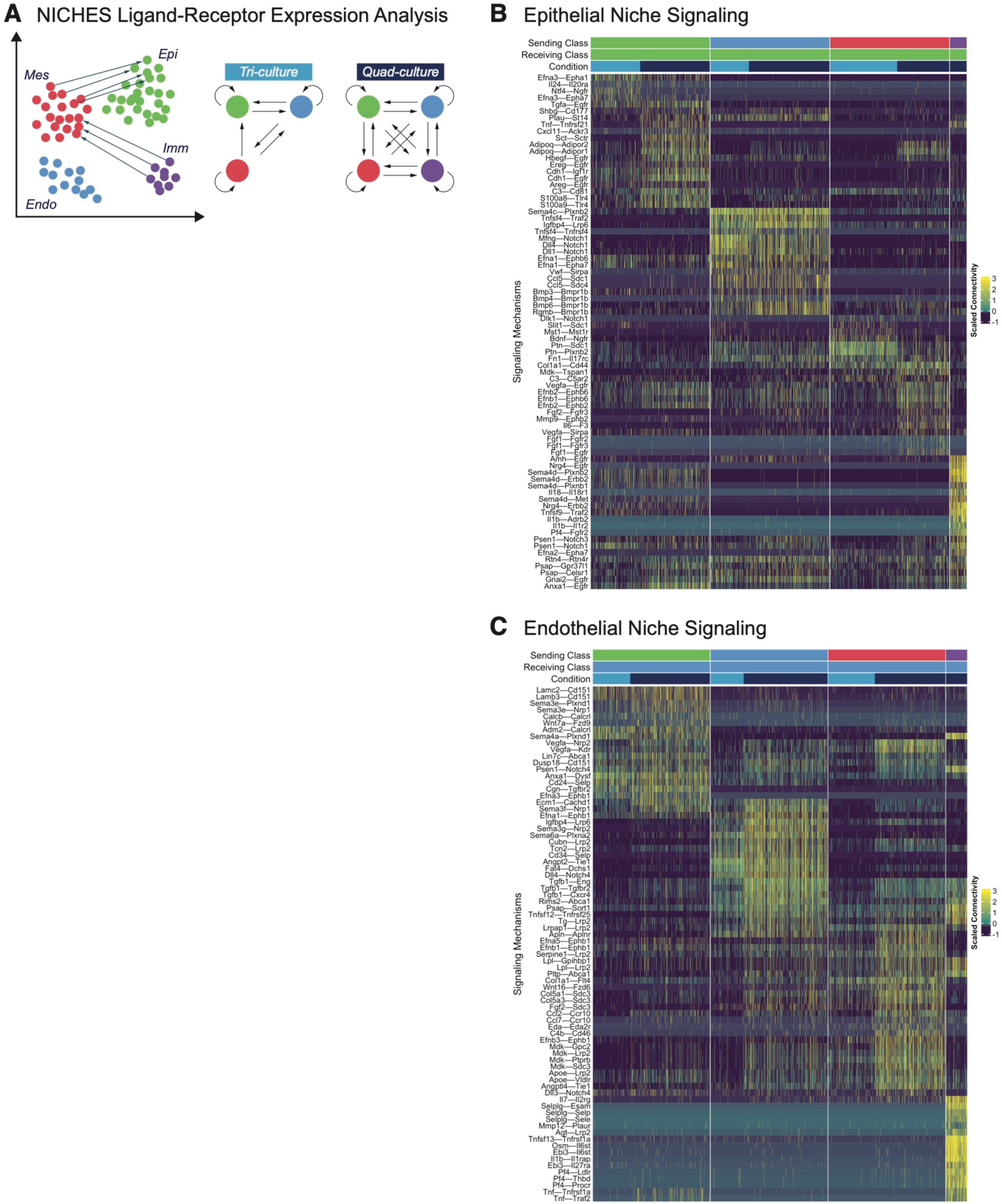
Epithelial and endothelial niche signaling, related to Figure 4. A) Schematic of NICHES ligand-receptor expression analysis, specifically comparing tri- and quad-culture engineered lungs. Heatmaps of top signaling mechanisms in the B) epithelial and C) endothelial niches, plotted by sending cell and comparing mechanism expression in tri- and quad-culture lungs.

**Figure S6.**
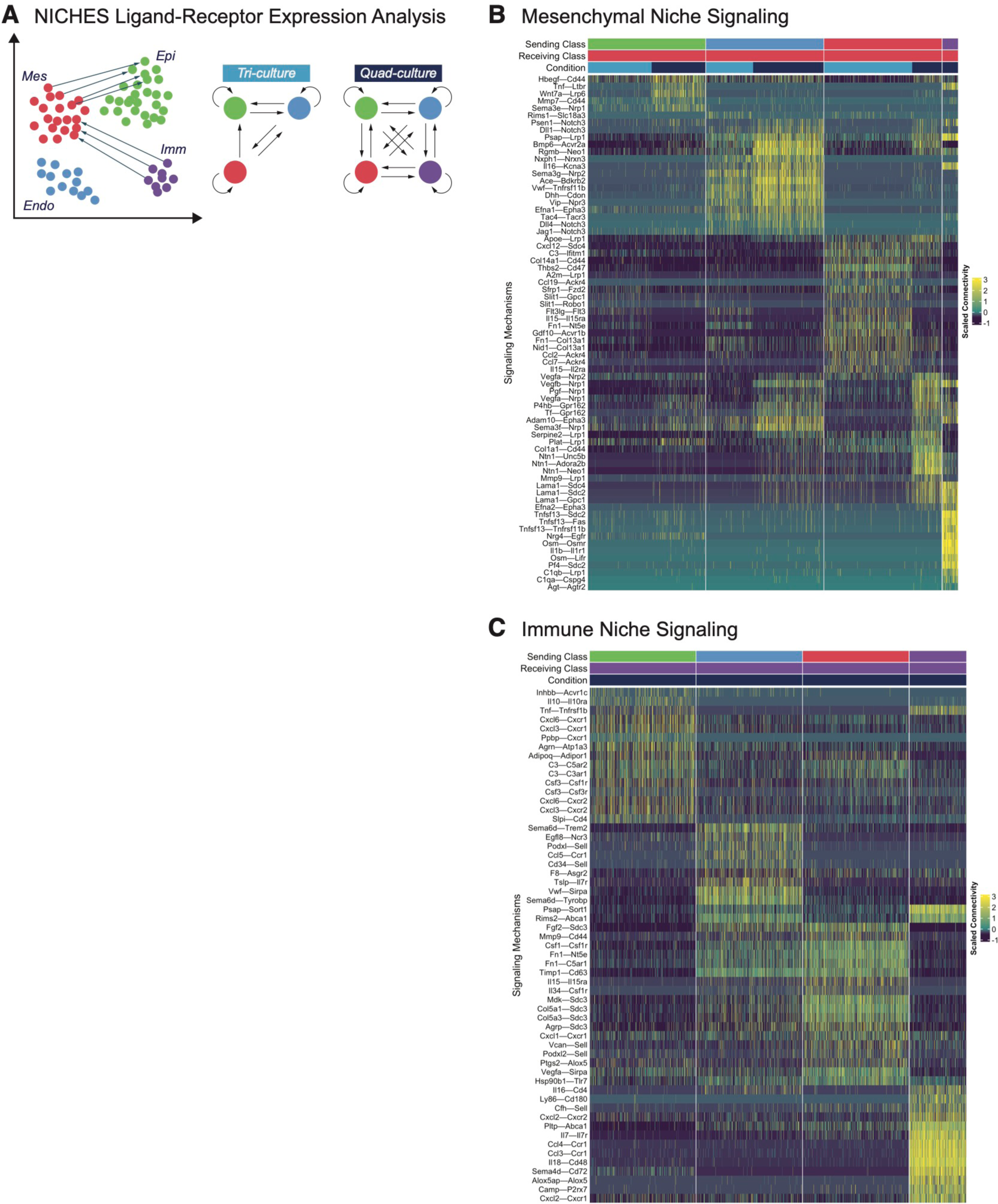
Mesenchymal and immune niche signaling, related to Figure 4. A) Schematic of NICHES ligand-receptor expression analysis, specifically comparing tri- and quad-culture engineered lungs. Heatmaps of top signaling mechanisms in the B) mesenchymal and C) immune cell niches, plotted by sending cell and comparing mechanism expression in tri- and quad-culture lungs.

## Notes

### Competing Interest Statement

The authors have declared no competing interest.

