## Supplementary material for "Engineered Whole Lungs for Tissue Biology": Document S1

I. Filtering individual objects from DGEs

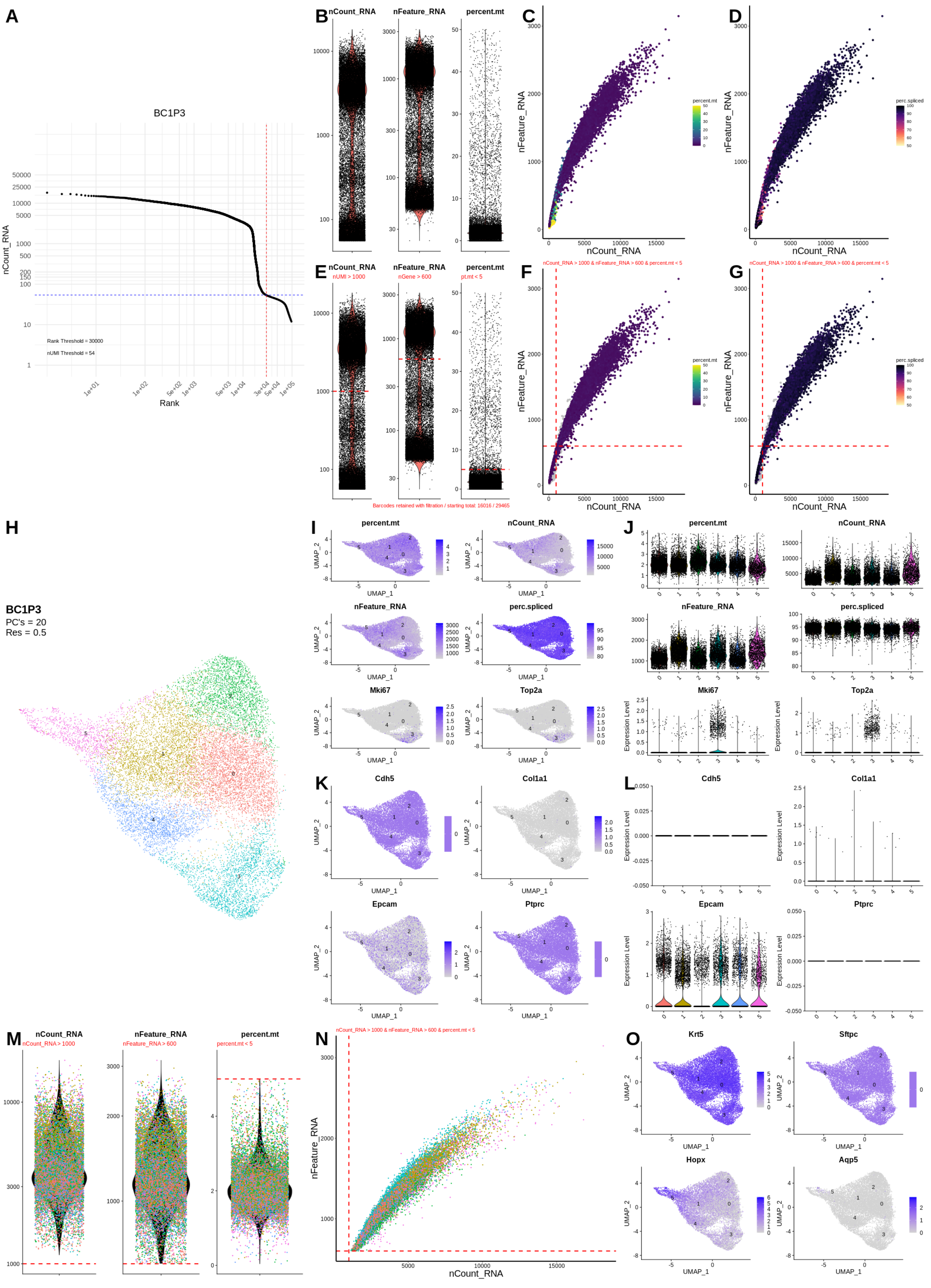

A
BC1P3
Rank Threshold = 30000
nUMI Threshold = 54
nCount_RNA
Rank
B
nCount_RNA
nFeature_RNA
percent.mt
C
D
nFeature_RNA
nCount_RNA
percent.mt
perc.spliced
E
nUMI > 1000
nGene > 600
pt.mt < 5
F
G
nCount_RNA > 1000 & nFeature_RNA > 600 & percent.mt < 5
Barcodes retained with filtration / starting total: 16016 / 29465
H
BC1P3
PC's = 20
Res = 0.5
I
J
Mki67
Top2a
UMAP_1
UMAP_2
Expression Level
K
L
Cdh5
Col1a1
Epcam
Ptprc
M
nCount_RNA > 1000
nFeature_RNA > 600
percent.mt < 5
N
O
Krt5
Sftpc
Hopx
Aqp5

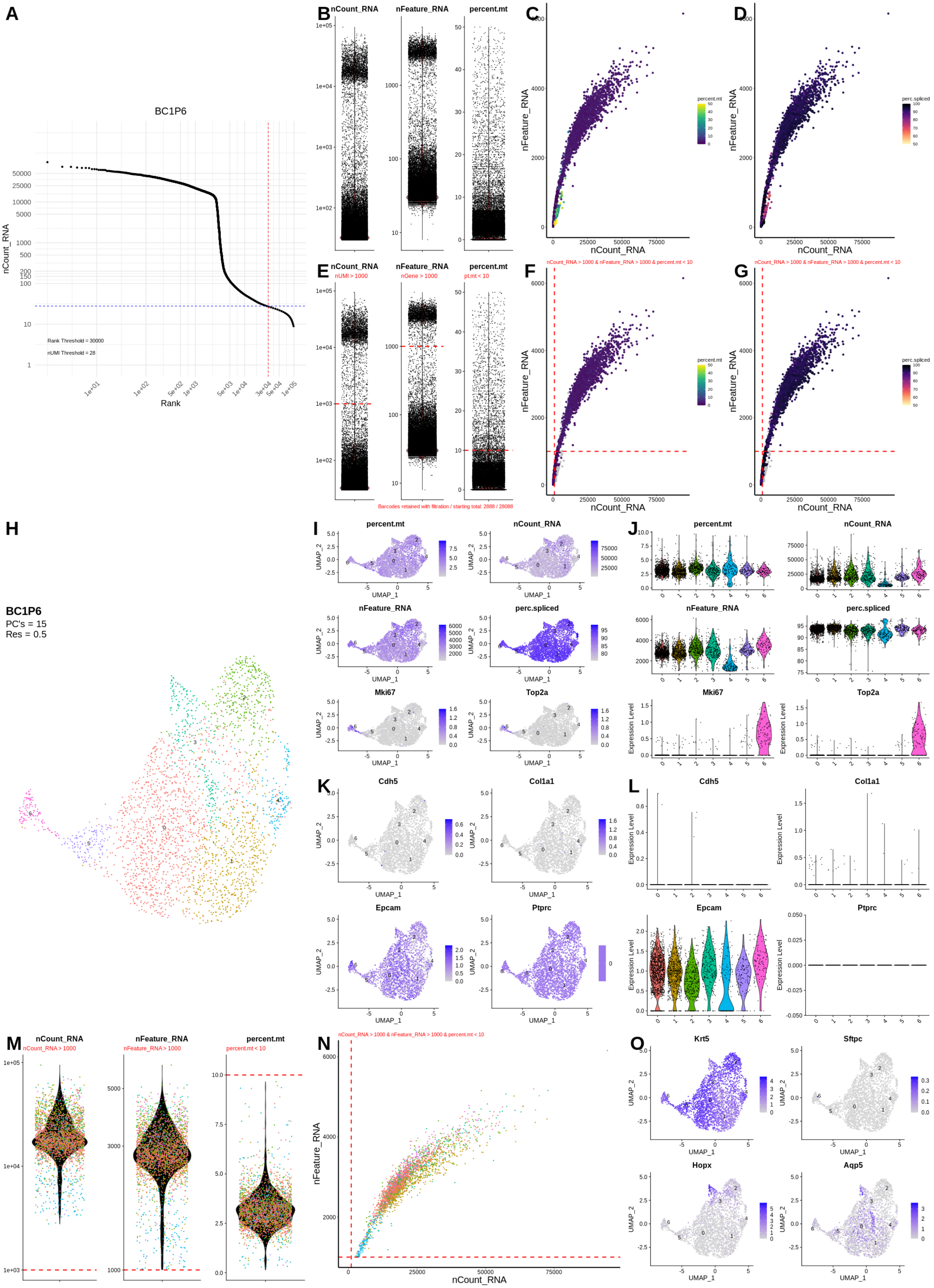
A
BC1P6
nCount_RNA
Rank
Rank Threshold = 30000
nUMI Threshold = 28
B
nCount_RNA
nFeature_RNA
percent.mt
C
nFeature_RNA
nCount_RNA
percent.mt
D
perc.spliced
E
nUMI > 1000
nGene > 1000
pt.mt < 10
Barcodes retained with filtration / starting total: 2888 / 28088
F
nCount_RNA > 1000 & nFeature_RNA > 1000 & percent.mt < 10
G
H
BC1P6
PC's = 15
Res = 0.5
I
J
Mki67
Top2a
UMAP_1
UMAP_2
Expression Level
K
Cdh5
Col1a1
L
Epcam
Ptprc
M
nCount_RNA > 1000
nFeature_RNA > 1000
percent.mt < 10
N
O
Krt5
Sftpc
Hopx
Aqp5

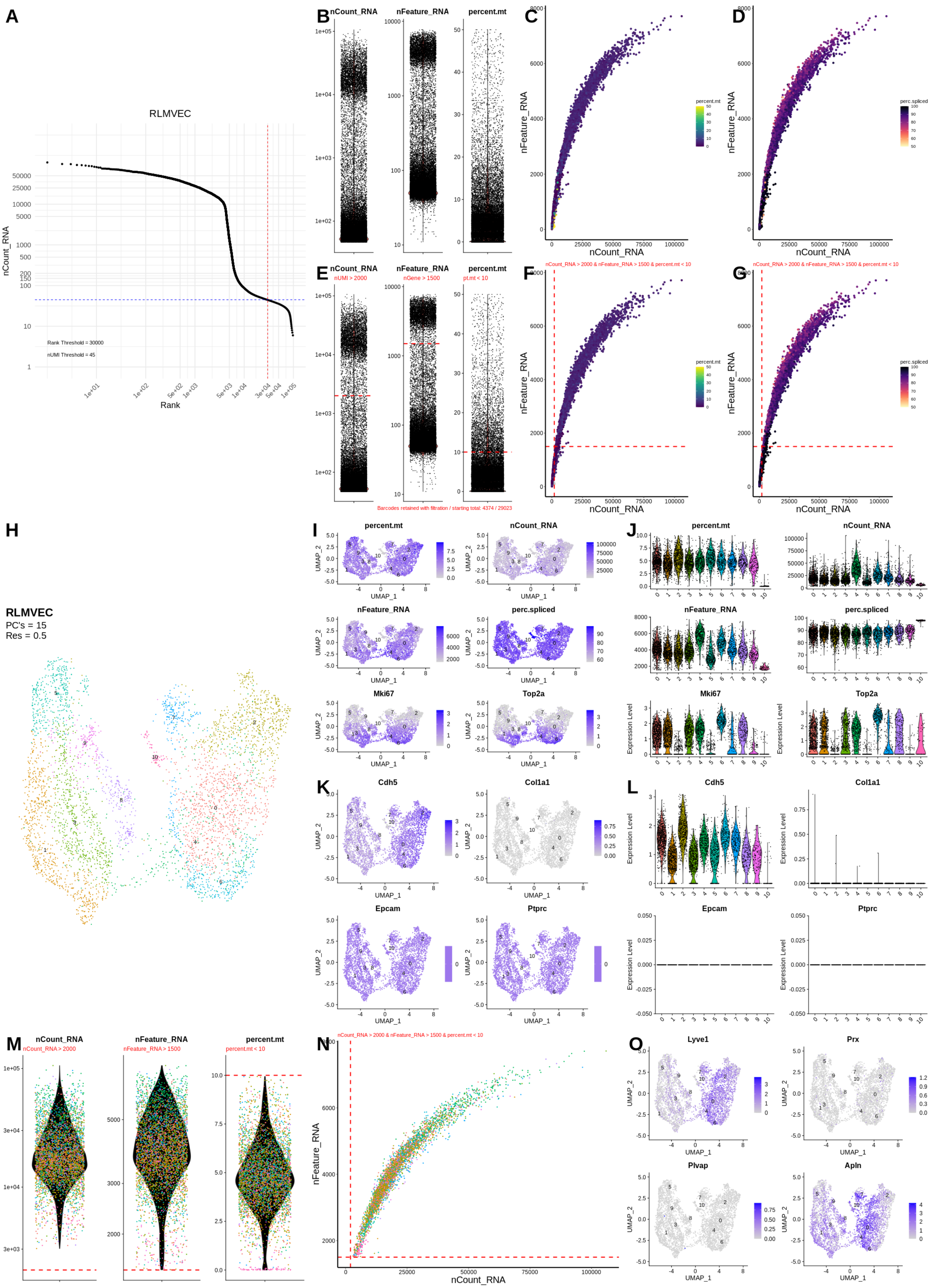

A
RLMVEC
nCount_RNA
Rank
Rank Threshold = 30000
nUMI Threshold = 45
B
C
D
E
nUMI > 2000
nGene > 1500
pt.mt < 10
F
G
nCount_RNA > 2000 & nFeature_RNA > 1500 & percent.mt < 10
Barcodes retained with filtration / starting total: 4374 / 29023
nFeature_RNA
percent.mt
perc.spliced
H
RLMVEC
PC's = 15
Res = 0.5
I
J
K
L
Mki67
Top2a
Cdh5
Col1a1
Epcam
Ptprc
UMAP_1
UMAP_2
Expression Level
M
N
O
Lyve1
Prx
Plvap
Apln

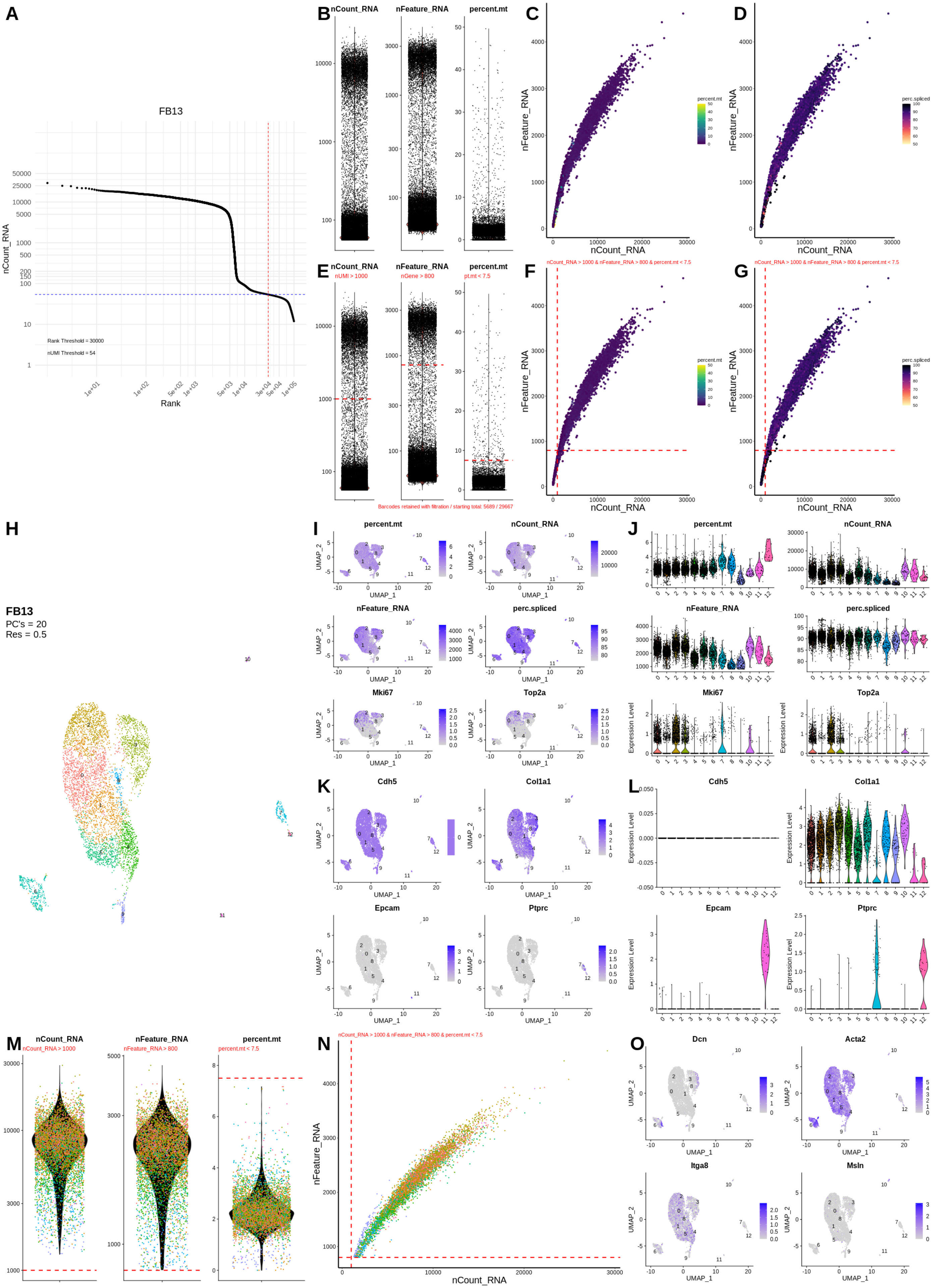

A
FB13
nCount_RNA
Rank
Rank Threshold = 30000
nUMI Threshold = 54
B
nCount_RNA
nFeature_RNA
percent.mt
C
D
nFeature_RNA
nCount_RNA
percent.mt
perc.spliced
E
nUMI > 1000
nGene > 800
pt.mt < 7.5
Barcodes retained with filtration / starting total: 5689 / 29667
F
G
nCount_RNA > 1000 & nFeature_RNA > 800 & percent.mt < 7.5
H
FB13
PC's = 20
Res = 0.5
I
J
Mki67
Top2a
Expression Level
UMAP_1
UMAP_2
K
L
Cdh5
Col1a1
Epcam
Ptprc
M
nCount_RNA > 1000
nFeature_RNA > 800
percent.mt < 7.5
N
O
Dcn
Acta2
Itga8
Msln

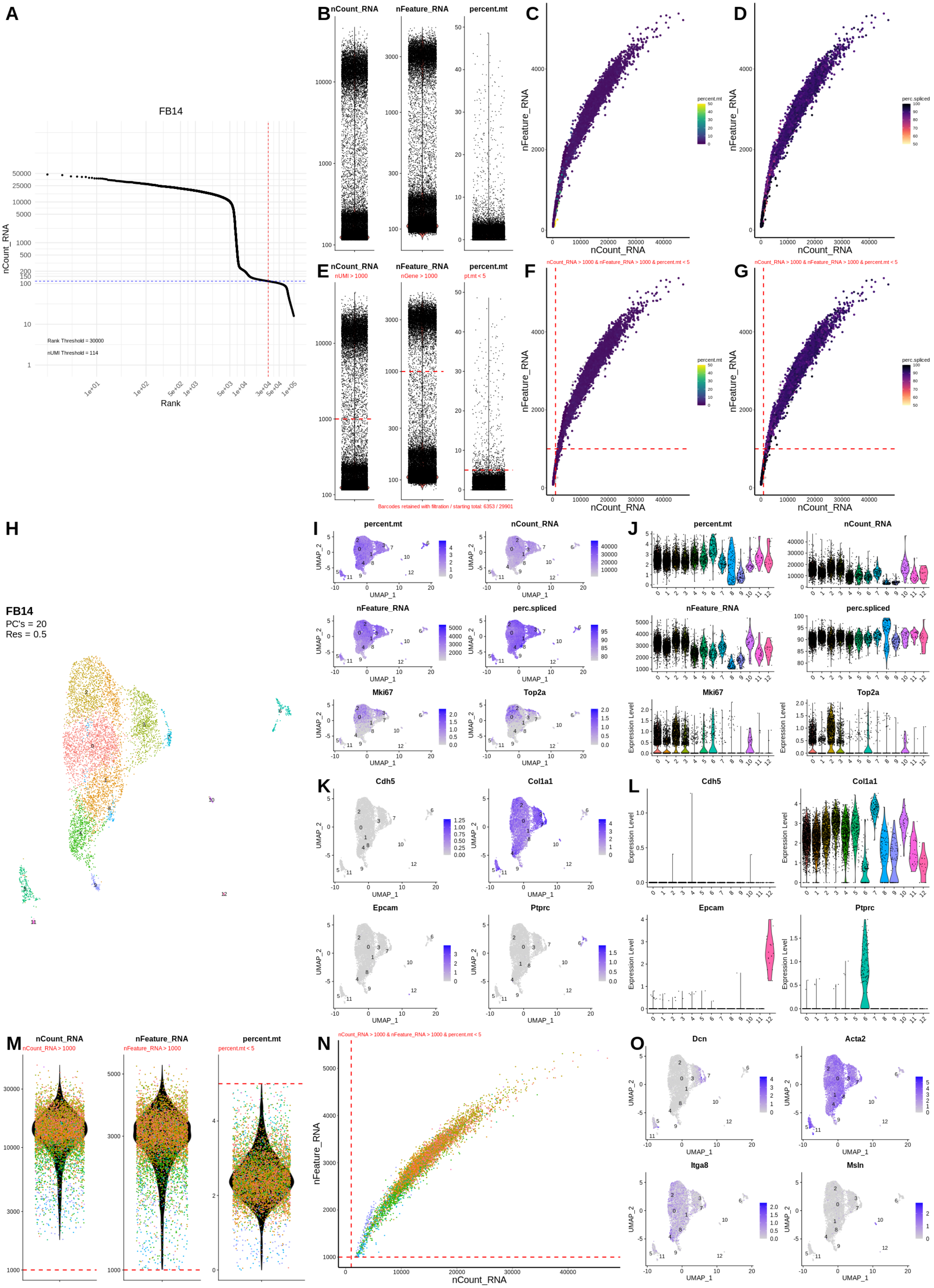
A
FB14
nCount_RNA
Rank
Rank Threshold = 30000
nUMI Threshold = 114
B
nCount_RNA
nFeature_RNA
percent.mt
C
nFeature_RNA
nCount_RNA
percent.mt
D
perc.spliced
E
nUMI > 1000
nGene > 1000
pt.mt < 5
Barcodes retained with filtration / starting total: 6353 / 29901
F
nCount_RNA > 1000 & nFeature_RNA > 1000 & percent.mt < 5
G
H
FB14
PC's = 20
Res = 0.5
I
J
nFeature_RNA
perc.spliced
Mki67
Top2a
UMAP_1
UMAP_2
Expression Level
K
L
Cdh5
Col1a1
Epcam
Ptprc
M
nCount_RNA > 1000
nFeature_RNA > 1000
percent.mt < 5
N
O
Dcn
Acta2
Itga8
Msln

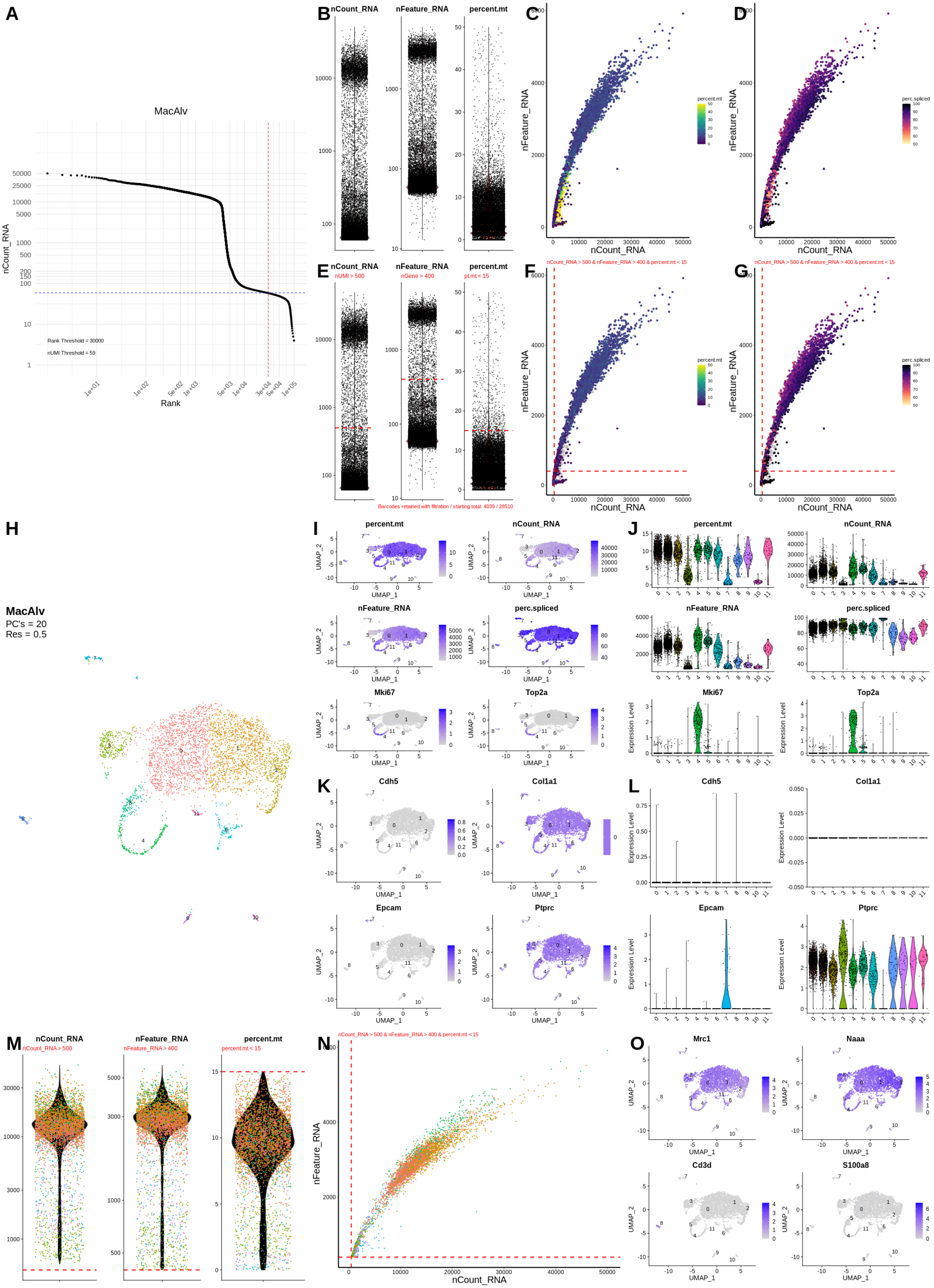
A
MacAlv
nCount_RNA
Rank
Rank Threshold = 30000
nUMI Threshold = 59
B
nCount_RNA
nFeature_RNA
percent.mt
C
D
nFeature_RNA
nCount_RNA
percent.mt
perc.spliced
E
nUMI > 500
nGene > 400
pt.mt < 15
Barcodes retained with filtration / starting total: 4039 / 28510
F
G
nCount_RNA > 500 & nFeature_RNA > 400 & percent.mt < 15
H
MacAlv
PC's = 20
Res = 0.5
I
J
Mki67
Top2a
UMAP_1
UMAP_2
Expression Level
K
L
Cdh5
Col1a1
Epcam
Ptprc
M
nCount_RNA > 500
nFeature_RNA > 400
percent.mt < 15
N
O
Mrc1
Naaa
Cd3d
S100a8

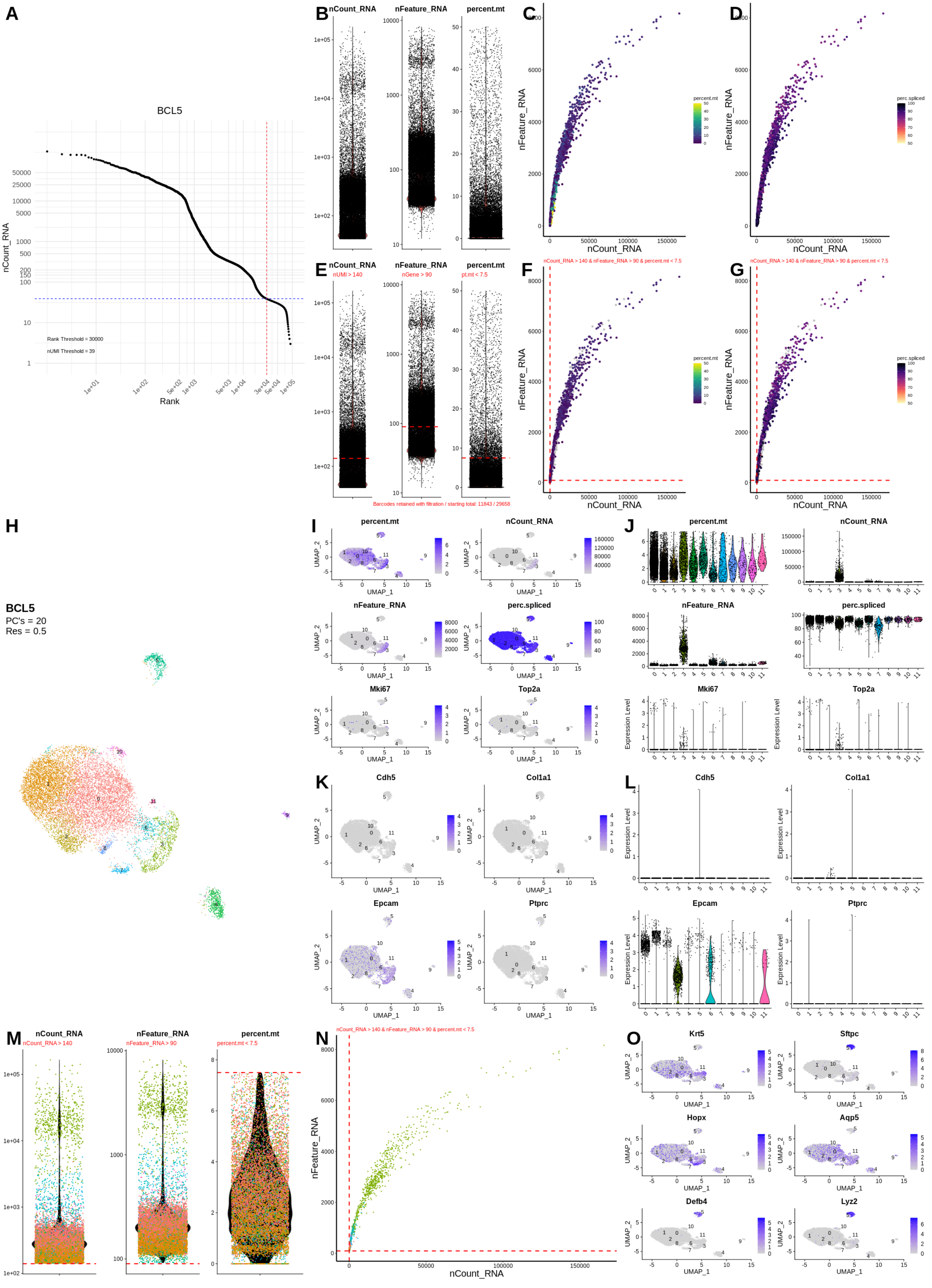

A
BCL5
nCount_RNA
Rank
Rank Threshold = 30000
nUMI Threshold = 39
B
nCount_RNA
nFeature_RNA
percent.mt
C
nFeature_RNA
nCount_RNA
percent.mt
D
perc.spliced
E
nUMI > 140
nGene > 90
pt.mt < 7.5
Barcodes retained with filtration / starting total: 11843 / 29658
F
nCount_RNA > 140 & nFeature_RNA > 90 & percent.mt < 7.5
G
H
BCL5
PC's = 20
Res = 0.5
I
UMAP_1
UMAP_2
J
Mki67
Top2a
Expression Level
K
Cdh5
Col1a1
Epcam
Ptprc
L
M
nCount_RNA > 140
nFeature_RNA > 90
percent.mt < 7.5
N
O
Krt5
Sftpc
Hopx
Aqp5
Defb4
Lyz2

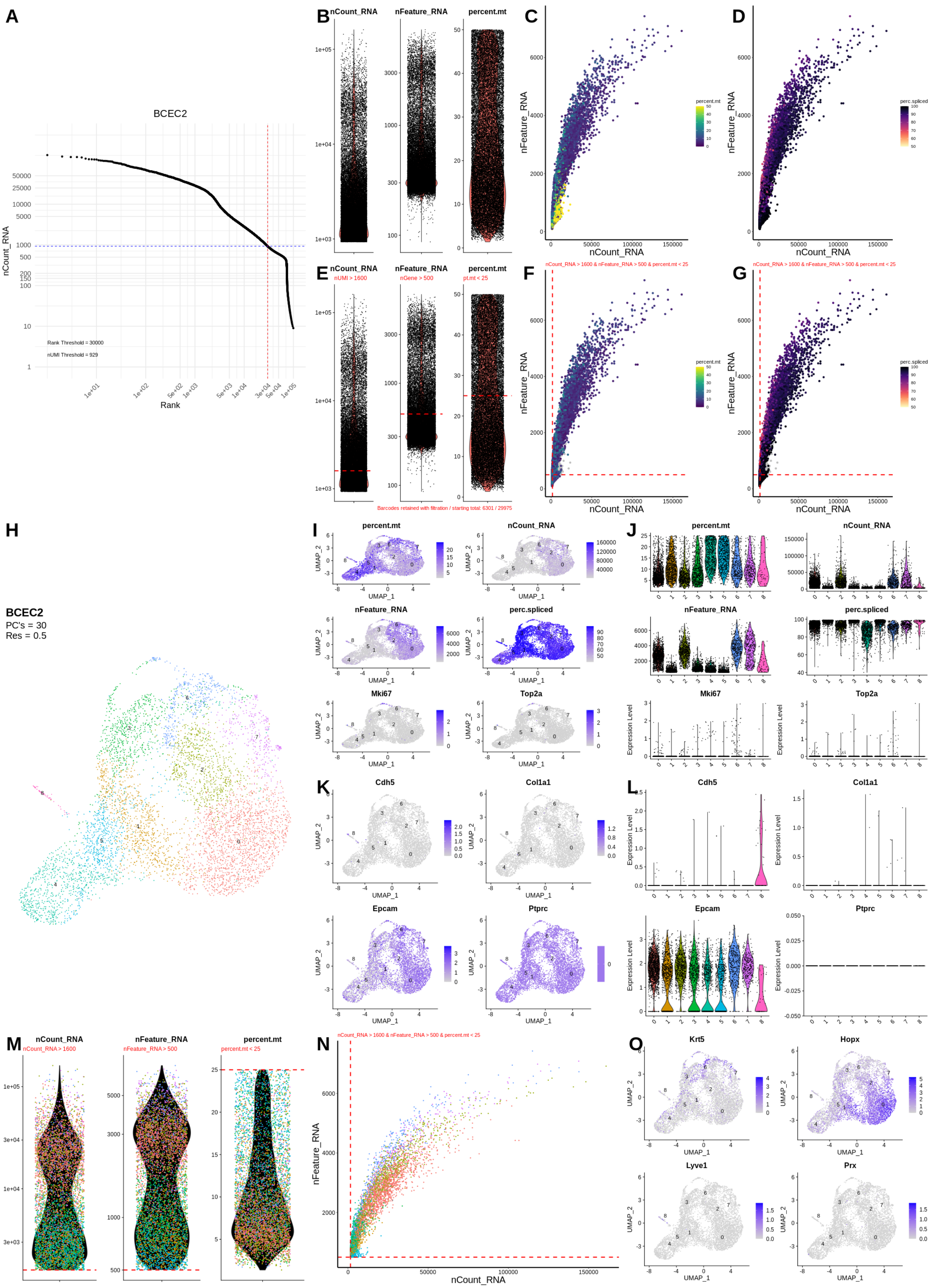
A
BCEC2
nCount_RNA
Rank
Rank Threshold = 30000
nUMI Threshold = 929
B
nCount_RNA
nFeature_RNA
percent.mt
C
nFeature_RNA
nCount_RNA
percent.mt
D
perc.spliced
E
nUMI > 1600
nGene > 500
pt.mt < 25
Barcodes retained with filtration / starting total: 6301 / 29975
F
nCount_RNA > 1600 & nFeature_RNA > 500 & percent.mt < 25
G
H
BCEC2
PC's = 30
Res = 0.5
I
UMAP_1
UMAP_2
Mki67
Top2a
J
Expression Level
K
Cdh5
Col1a1
Epcam
Ptprc
L
M
nCount_RNA > 1600
nFeature_RNA > 500
percent.mt < 25
N
O
Krt5
Hopx
Lyve1
Prx

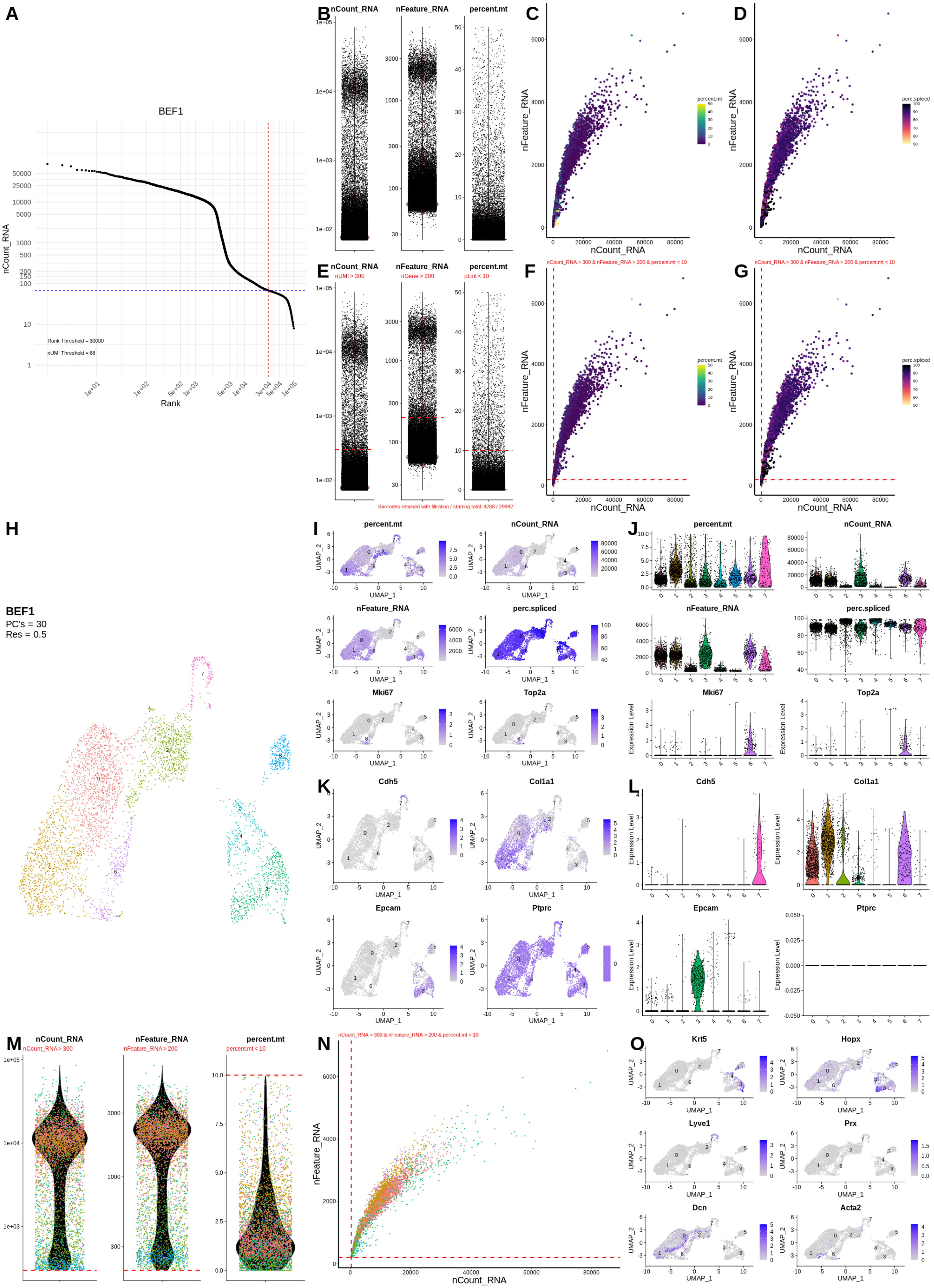
A
BEF1
nCount_RNA
Rank
Rank Threshold = 30000
nUMI Threshold = 68
B
nCount_RNA
nFeature_RNA
percent.mt
C
nFeature_RNA
nCount_RNA
percent.mt
D
perc.spliced
E
nUMI > 300
nGene > 200
pt.mt < 10
Barcodes retained with filtration / starting total: 4288 / 29992
F
nCount_RNA > 300 & nFeature_RNA > 200 & percent.mt < 10
G
H
BEF1
PC's = 30
Res = 0.5
I
J
Mki67
Top2a
UMAP_1
UMAP_2
Expression Level
K
Cdh5
Col1a1
Epcam
Ptprc
L
M
nCount_RNA > 300
nFeature_RNA > 200
percent.mt < 10
N
O
Krt5
Hopx
Lyve1
Prx
Dcn
Acta2

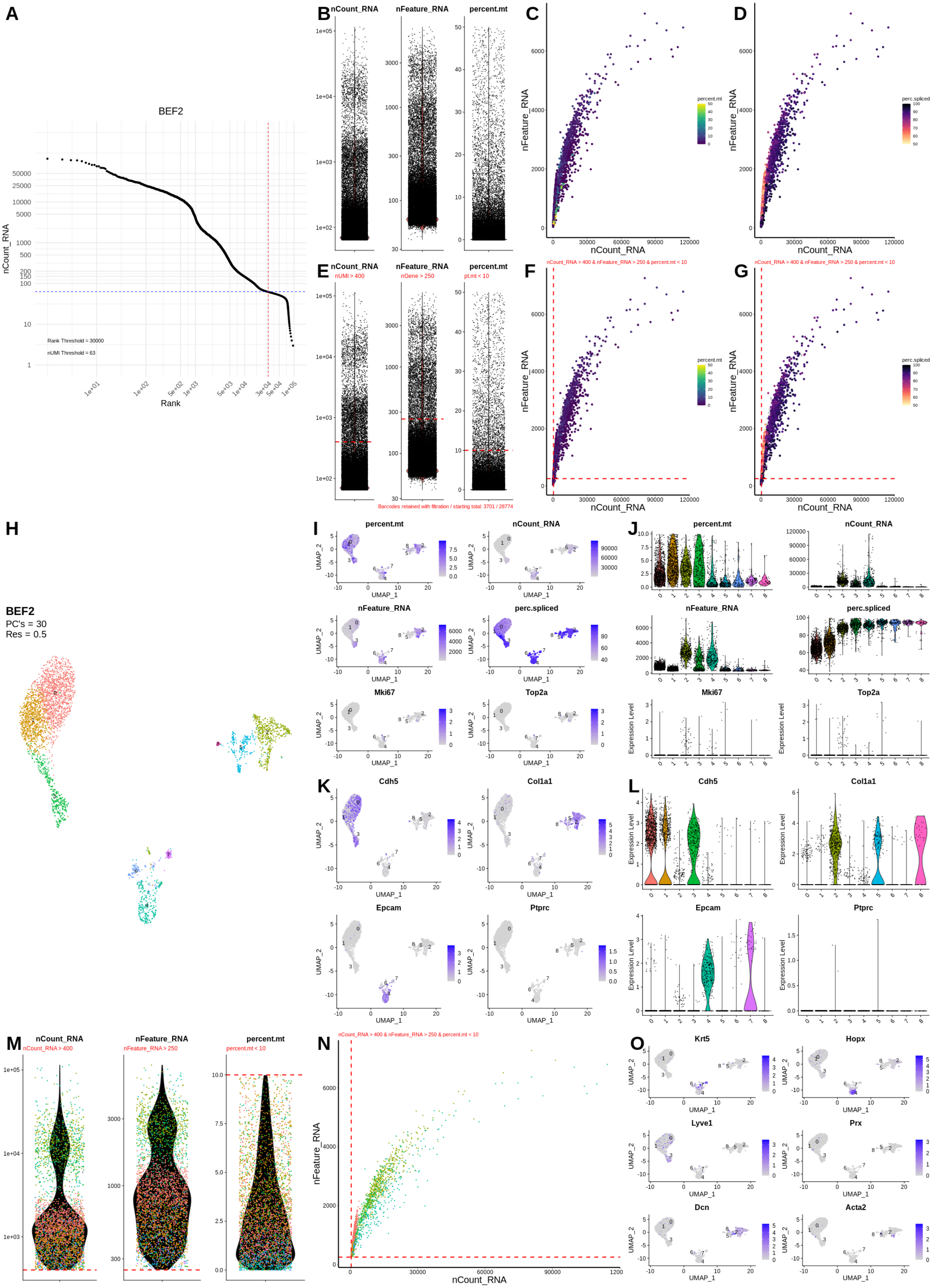
A
BEF2
nCount_RNA
Rank
Rank Threshold = 30000
nUMI Threshold = 63
B
nCount_RNA
nFeature_RNA
percent.mt
C
D
nFeature_RNA
nCount_RNA
percent.mt
perc.spliced
E
nUMI > 400
nGene > 250
pt.mt < 10
Barcodes retained with filtration / starting total: 3701 / 28774
F
G
nCount_RNA > 400 & nFeature_RNA > 250 & percent.mt < 10
H
BEF2
PC's = 30
Res = 0.5
I
J
Mki67
Top2a
UMAP_1
UMAP_2
Expression Level
K
L
Cdh5
Col1a1
Epcam
Ptprc
M
nCount_RNA > 400
nFeature_RNA > 250
percent.mt < 10
N
O
Krt5
Hopx
Lyve1
Prx
Dcn
Acta2

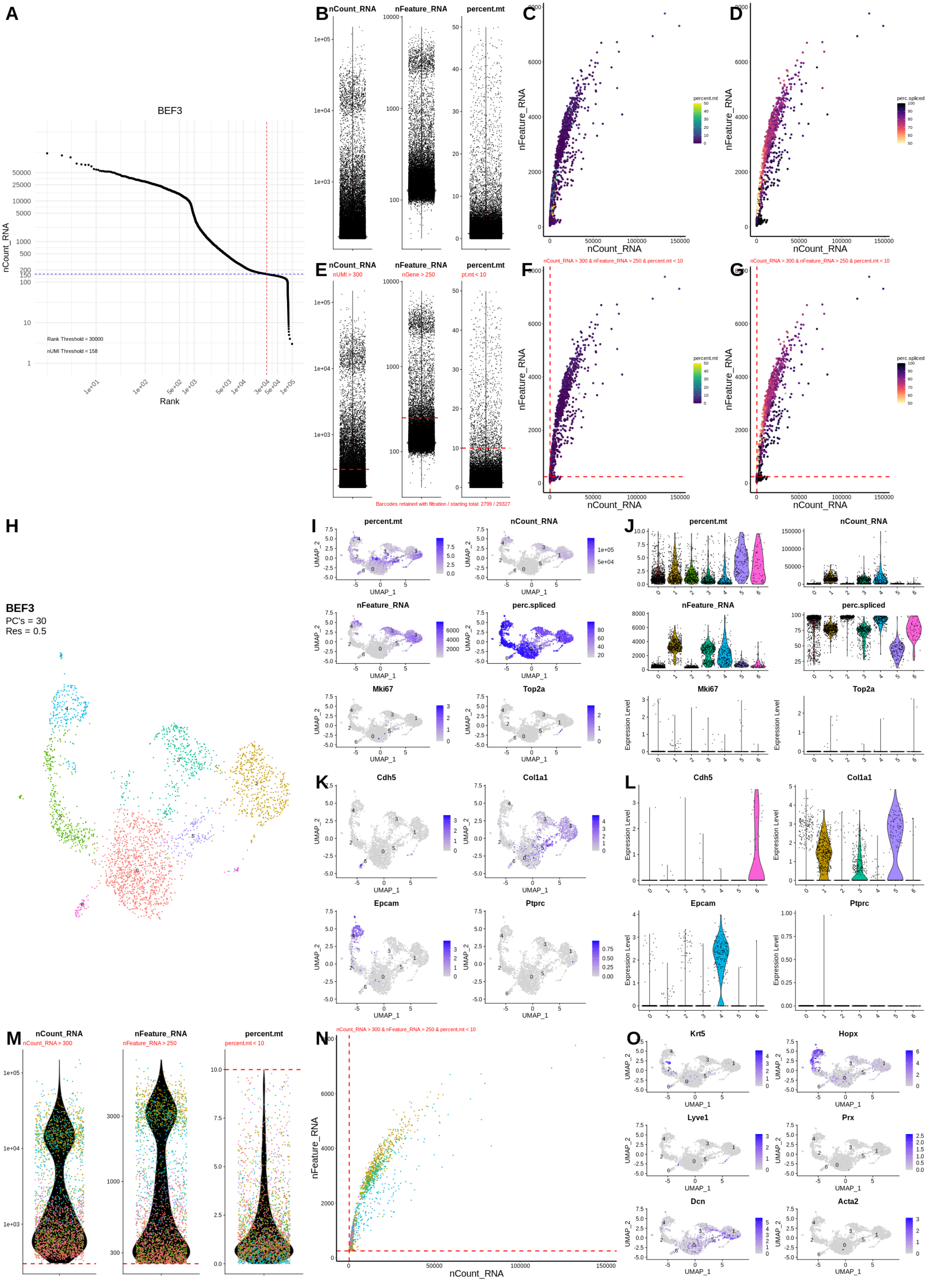

A
BEF3
nCount_RNA
Rank
Rank Threshold = 30000
nUMI Threshold = 158
B
nCount_RNA
nFeature_RNA
percent.mt
C
nFeature_RNA
nCount_RNA
percent.mt
D
perc.spliced
E
nUMI > 300
nGene > 250
pt.mt < 10
Barcodes retained with filtration / starting total: 2799 / 29327
F
nCount_RNA > 300 & nFeature_RNA > 250 & percent.mt < 10
G
H
BEF3
PC's = 30
Res = 0.5
I
UMAP_1
UMAP_2
J
Mki67
Top2a
Expression Level
K
Cdh5
Col1a1
L
Epcam
Ptprc
M
nCount_RNA > 300
nFeature_RNA > 250
percent.mt < 10
N
O
Krt5
Hopx
Lyve1
Prx
Dcn
Acta2

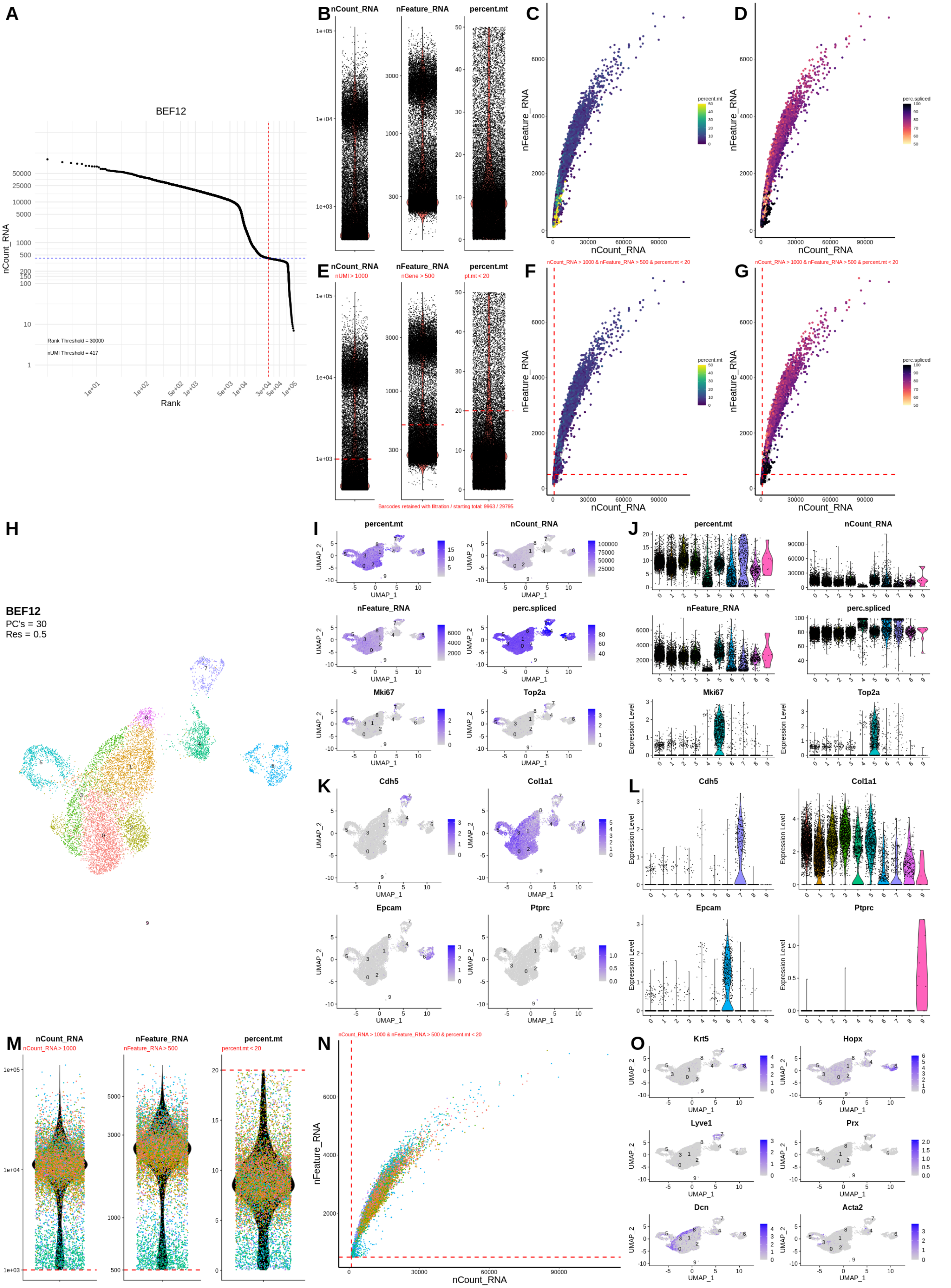

A
BEF12
nCount_RNA
Rank
Rank Threshold = 30000
nUMI Threshold = 417
B
nCount_RNA
nFeature_RNA
percent.mt
C
nFeature_RNA
nCount_RNA
percent.mt
D
perc.spliced
E
nUMI > 1000
nGene > 500
pt.mt < 20
F
nCount_RNA > 1000 & nFeature_RNA > 500 & percent.mt < 20
G
Barcodes retained with filtration / starting total: 9963 / 29795
H
BEF12
PC's = 30
Res = 0.5
I
UMAP_1
UMAP_2
perc.spliced
Mki67
Top2a
J
Expression Level
K
Cdh5
Col1a1
Epcam
Ptprc
L
M
nCount_RNA > 1000
nFeature_RNA > 500
percent.mt < 20
N
O
Krt5
Hopx
Lyve1
Prx
Dcn
Acta2

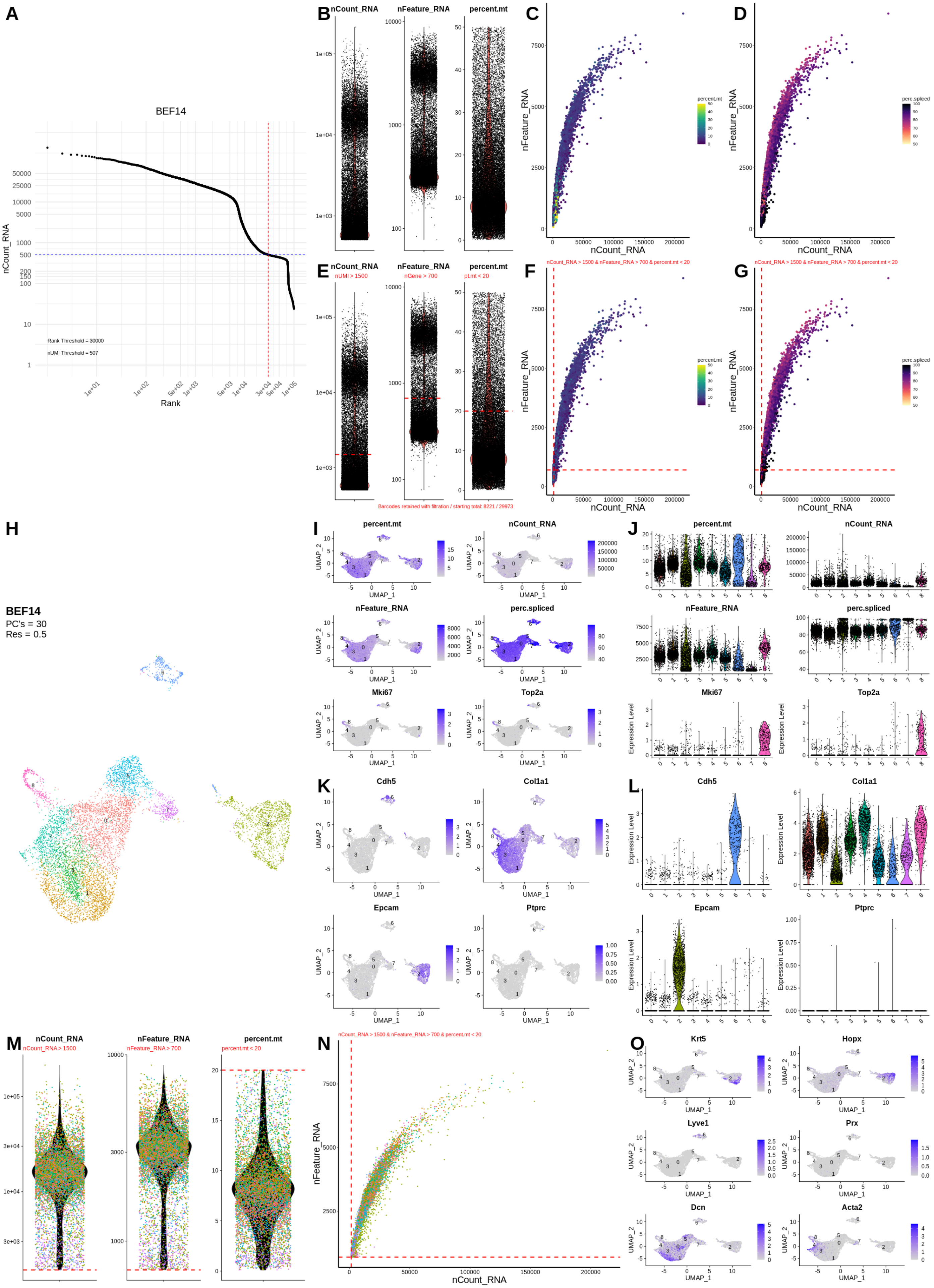
A
BEF14
nCount_RNA
Rank
Rank Threshold = 30000
nUMI Threshold = 507
B
nCount_RNA
nFeature_RNA
percent.mt
C
nFeature_RNA
nCount_RNA
percent.mt
D
perc.spliced
E
nUMI > 1500
nGene > 700
pt.mt < 20
Barcodes retained with filtration / starting total: 8221 / 29973
F
nCount_RNA > 1500 & nFeature_RNA > 700 & percent.mt < 20
G
H
BEF14
PC's = 30
Res = 0.5
I
UMAP_1
UMAP_2
Mki67
Top2a
J
Expression Level
K
Cdh5
Col1a1
Epcam
Ptprc
L
M
nCount_RNA > 1500
nFeature_RNA > 700
percent.mt < 20
N
O
Krt5
Hopx
Lyve1
Prx
Dcn
Acta2

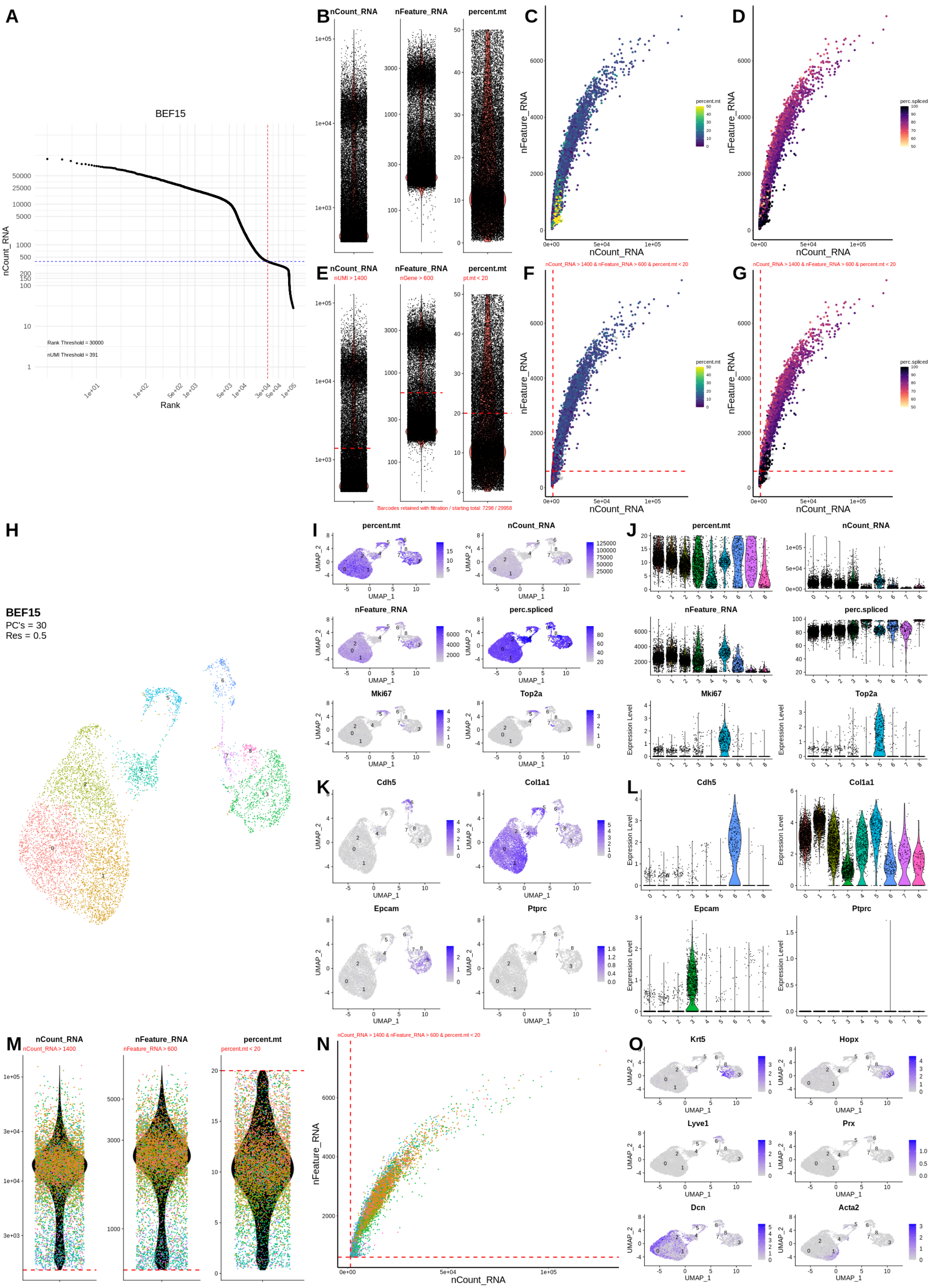

A
BEF15
Rank Threshold = 30000
nUMI Threshold = 391
nCount_RNA
Rank
B
nCount_RNA
nFeature_RNA
percent.mt
C
D
E
nUMI > 1400
nGene > 600
pt.mt < 20
Barcodes retained with filtration / starting total: 7298 / 29958
F
G
nCount_RNA > 1400 & nFeature_RNA > 600 & percent.mt < 20
H
BEF15
PC's = 30
Res = 0.5
I
J
Mki67
Top2a
perc.spliced
K
L
Cdh5
Col1a1
Epcam
Ptprc
M
nCount_RNA > 1400
nFeature_RNA > 600
percent.mt < 20
N
O
Krt5
Hopx
Lyve1
Prx
Dcn
Acta2
UMAP_1
UMAP_2
Expression Level

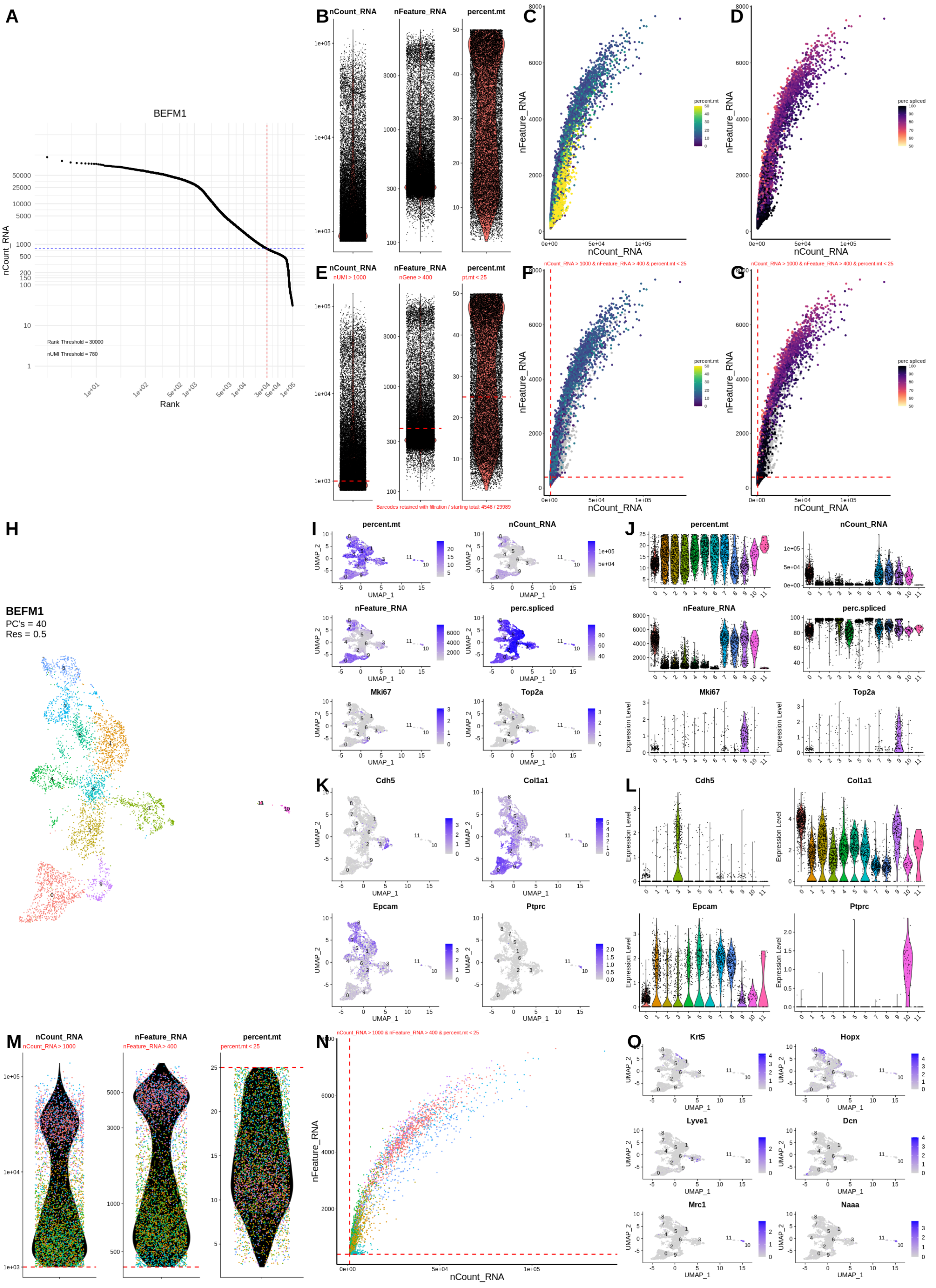
A
BEFM1
nCount_RNA
Rank
Rank Threshold = 30000
nUMI Threshold = 780
B
nCount_RNA
nFeature_RNA
percent.mt
C
D
nFeature_RNA
nCount_RNA
percent.mt
perc.spliced
E
nUMI > 1000
nGene > 400
pt.mt < 25
F
G
nCount_RNA > 1000 & nFeature_RNA > 400 & percent.mt < 25
Barcodes retained with filtration / starting total: 4548 / 29989
H
BEFM1
PC's = 40
Res = 0.5
I
J
Mki67
Top2a
UMAP_1
UMAP_2
Expression Level
K
L
Cdh5
Col1a1
Epcam
Ptprc
M
nCount_RNA > 1000
nFeature_RNA > 400
percent.mt < 25
N
O
Krt5
Hopx
Lyve1
Dcn
Mrc1
Naaa

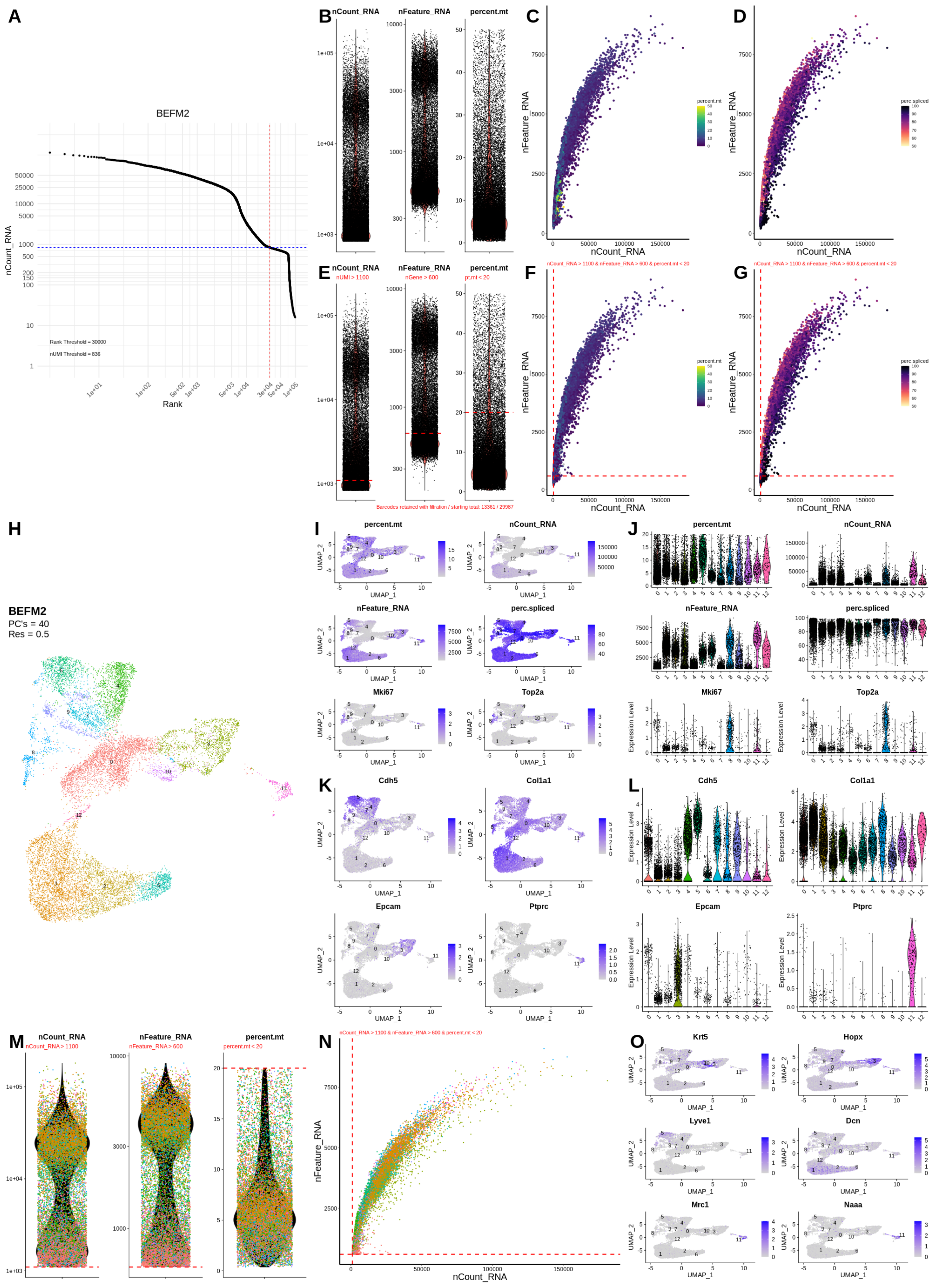
A
BEFM2
nCount_RNA
Rank
Rank Threshold = 30000
nUMI Threshold = 836
B
nCount_RNA
nFeature_RNA
percent.mt
C
D
nFeature_RNA
nCount_RNA
percent.mt
perc.spliced
E
nUMI > 1100
nGene > 600
pt.mt < 20
F
G
nCount_RNA > 1100 & nFeature_RNA > 600 & percent.mt < 20
Barcodes retained with filtration / starting total: 13361 / 29987
H
BEFM2
PC's = 40
Res = 0.5
I
J
UMAP_1
UMAP_2
Mki67
Top2a
Expression Level
K
L
Cdh5
Col1a1
Epcam
Ptprc
M
nCount_RNA > 1100
nFeature_RNA > 600
percent.mt < 20
N
O
Krt5
Hopx
Lyve1
Dcn
Mrc1
Naaa

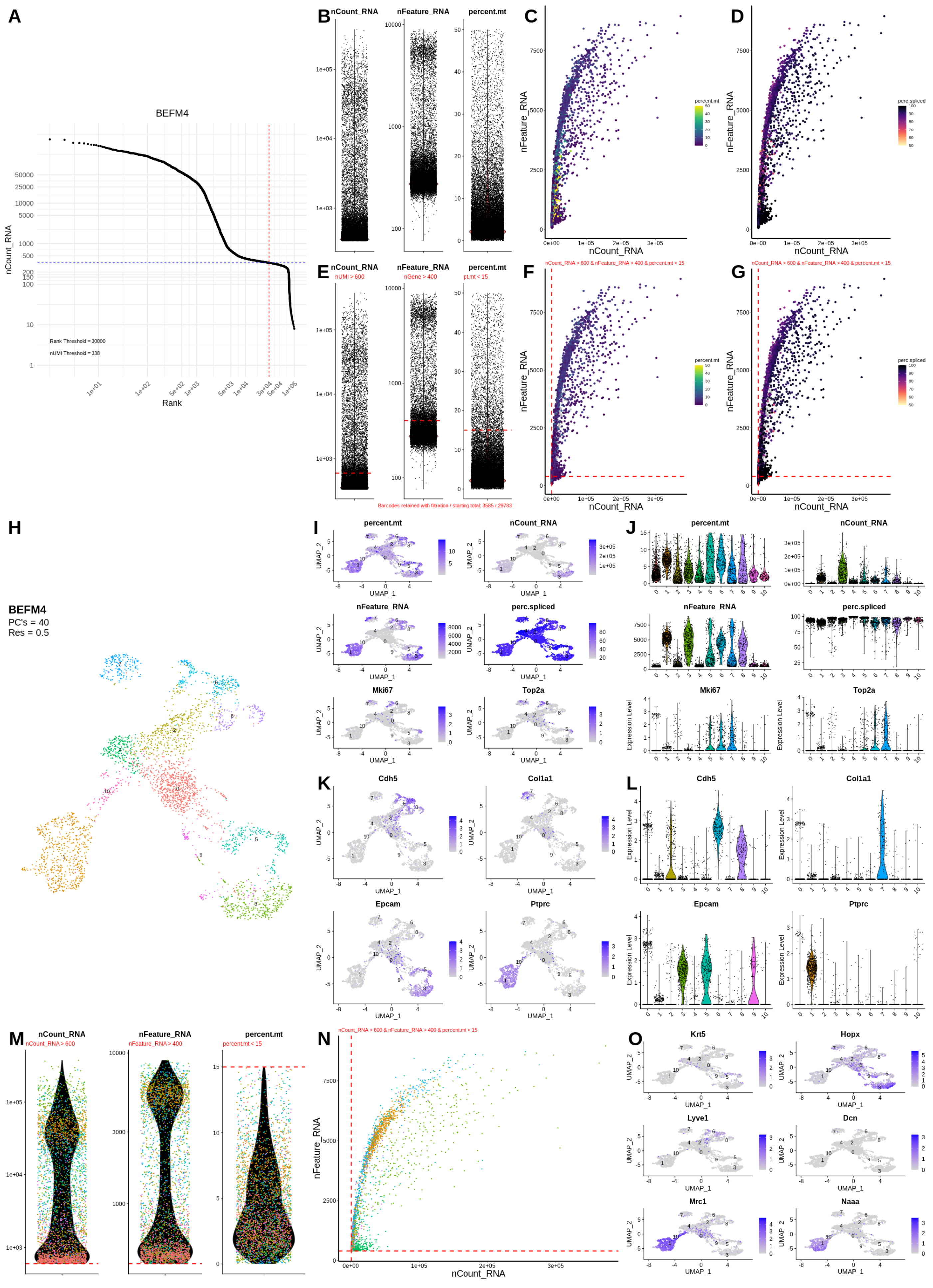
A
BEFM4
nCount_RNA
Rank
Rank Threshold = 30000
nUMI Threshold = 338
B
nCount_RNA
nFeature_RNA
percent.mt
C
D
nFeature_RNA
nCount_RNA
percent.mt
perc.spliced
E
nUMI > 600
nGene > 400
pt.mt < 15
Barcodes retained with filtration / starting total: 3585 / 29783
F
G
nCount_RNA > 600 & nFeature_RNA > 400 & percent.mt < 15
H
BEFM4
PC's = 40
Res = 0.5
I
J
Mki67
Top2a
UMAP_1
UMAP_2
Expression Level
K
L
Cdh5
Col1a1
Epcam
Ptprc
M
nCount_RNA > 600
nFeature_RNA > 400
percent.mt < 15
N
O
Krt5
Hopx
Lyve1
Dcn
Mrc1
Naaa

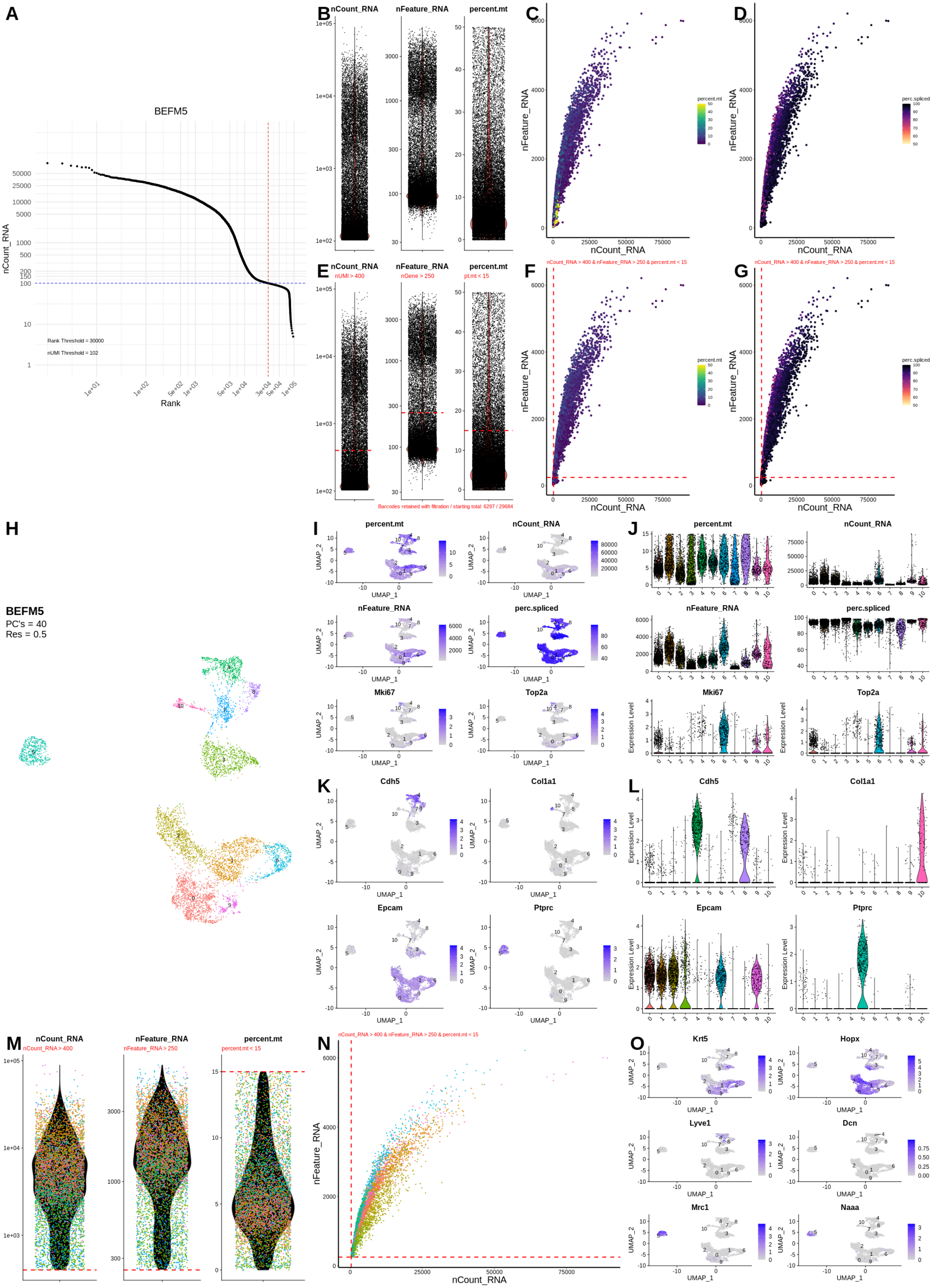
A
BEFM5
nCount_RNA
Rank
Rank Threshold = 30000
nUMI Threshold = 102
B
nCount_RNA
nFeature_RNA
percent.mt
C
nFeature_RNA
nCount_RNA
percent.mt
D
perc.spliced
E
nUMI > 400
nGene > 250
pt.mt < 15
Barcodes retained with filtration / starting total: 6297 / 29684
F
nCount_RNA > 400 & nFeature_RNA > 250 & percent.mt < 15
G
H
BEFM5
PC's = 40
Res = 0.5
I
UMAP_1
UMAP_2
Mki67
Top2a
J
Expression Level
K
Cdh5
Col1a1
Epcam
Ptprc
L
M
nCount_RNA > 400
nFeature_RNA > 250
percent.mt < 15
N
O
Krt5
Hopx
Lyve1
Dcn
Mrc1
Naaa

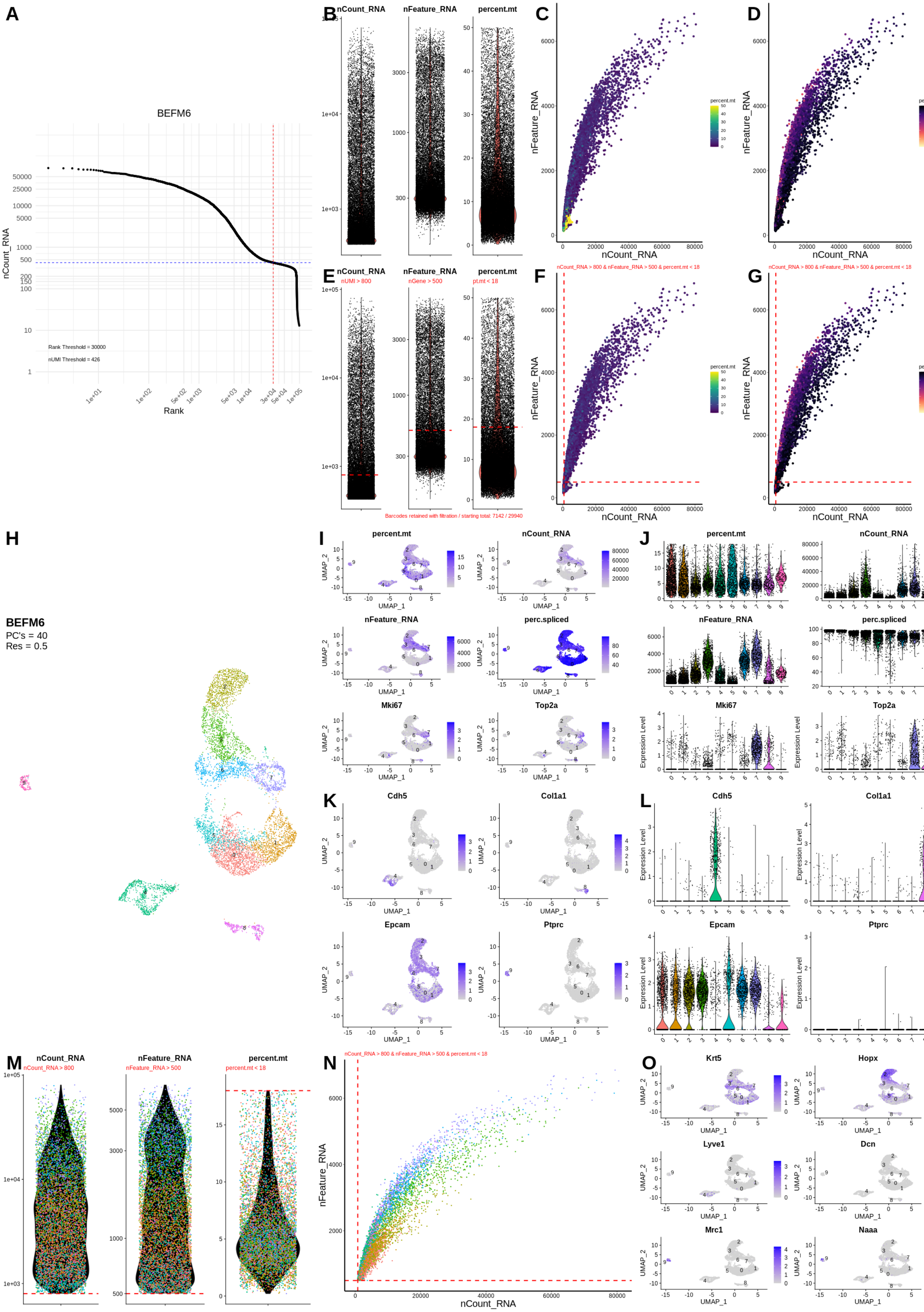
A
BEFM6
nCount_RNA
Rank
Rank Threshold = 30000
nUMI Threshold = 426
B
nCount_RNA
nFeature_RNA
percent.mt
C
nFeature_RNA
nCount_RNA
percent.mt
D
perc.spliced
E
nUMI > 800
nGene > 500
pt.mt < 18
F
nCount_RNA > 800 & nFeature_RNA > 500 & percent.mt < 18
G
Barcodes retained with filtration / starting total: 7142 / 29940
H
BEFM6
PC's = 40
Res = 0.5
I
UMAP_1
UMAP_2
J
Mki67
Top2a
Expression Level
K
Cdh5
Col1a1
Epcam
Ptprc
L
M
nCount_RNA > 800
nFeature_RNA > 500
percent.mt < 18
N
O
Krt5
Hopx
Lyve1
Dcn
Mrc1
Naaa

II. Cleaning sample objects grouped by chemistry & cell type similarity

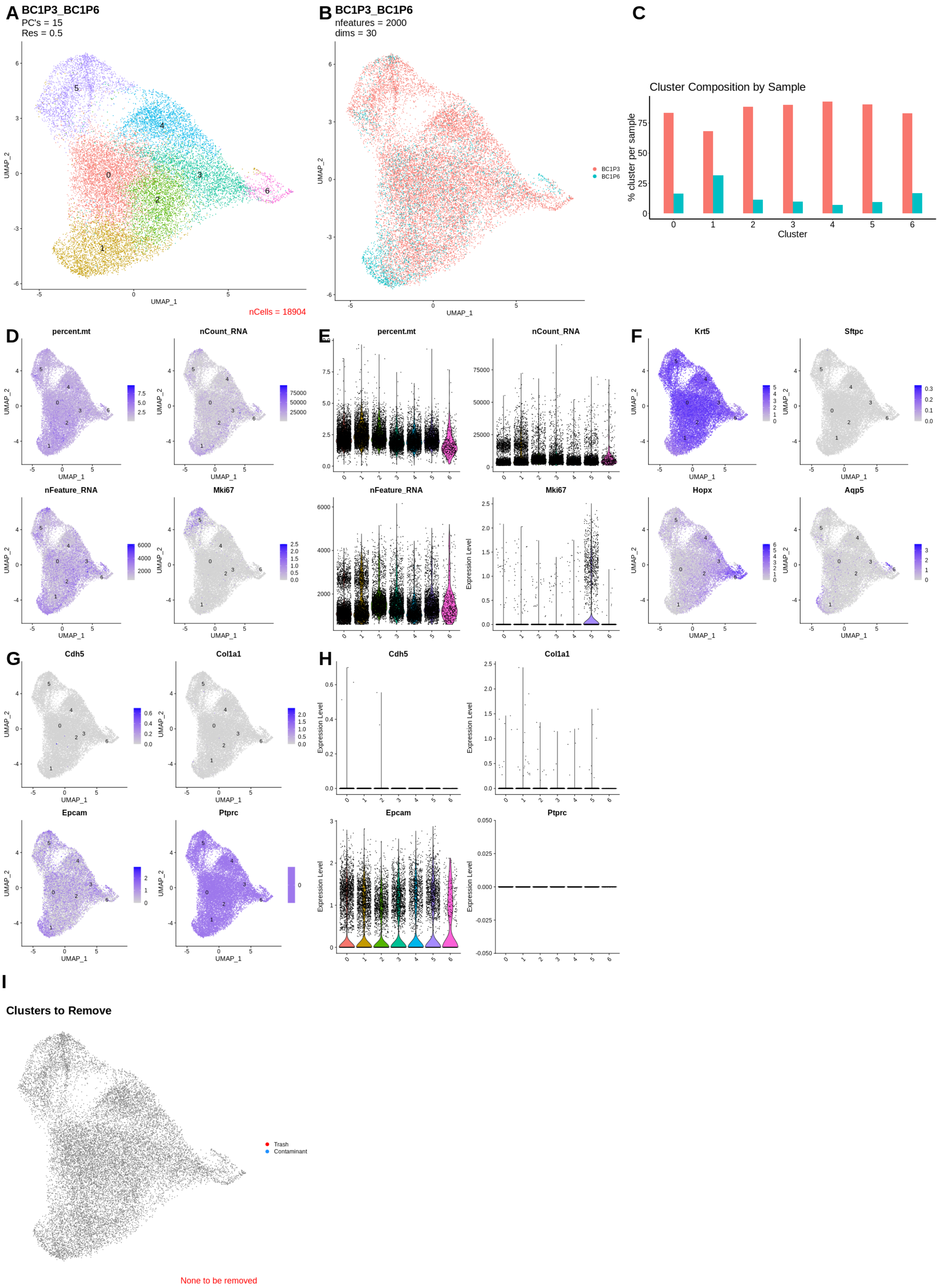

A
BC1P3_BC1P6
PC's = 15
Res = 0.5
UMAP_1
UMAP_2
nCells = 18904
B
BC1P3_BC1P6
nfeatures = 2000
dims = 30
BC1P3
BC1P6
C
Cluster Composition by Sample
% cluster per sample
Cluster
D
percent.mt
nCount_RNA
nFeature_RNA
Mki67
E
percent.mt
nCount_RNA
nFeature_RNA
Mki67
Expression Level
F
Krt5
Sftpc
Hopx
Aqp5
G
Cdh5
Col1a1
Epcam
Ptprc
H
Cdh5
Col1a1
Epcam
Ptprc
I
Clusters to Remove
Trash
Contaminant
None to be removed

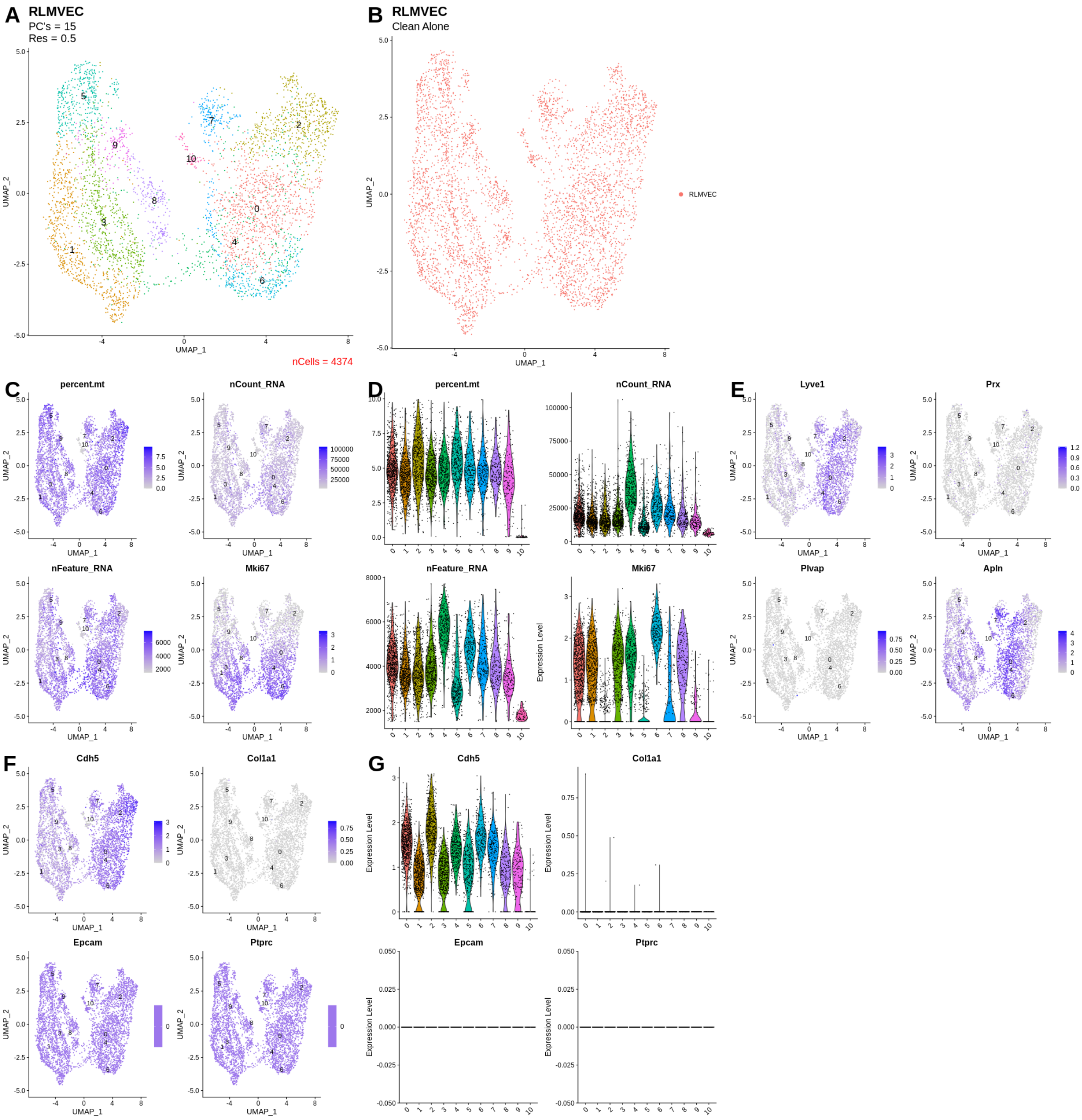


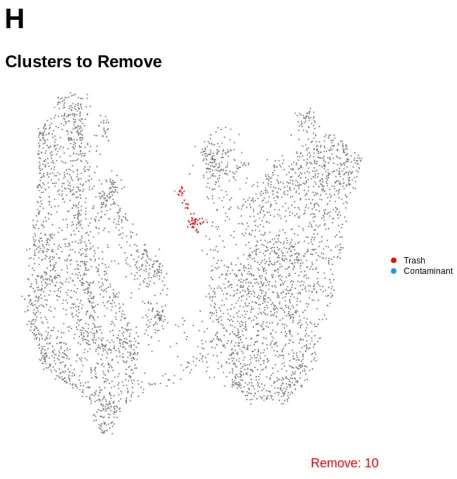


I

| | p_val | avg_log2FC | pct.1 | pct.2 | p_val_adj | ratio | cluster |
|---|---|---|---|---|---|---|---|
| *Mrps10* | 0.0263 | 0.4155 | 0.2 | 0.128 | 1 | 1.5625 | 10 |
| *Sgcg* | 0.0271 | 0.4065 | 0.217 | 0.145 | 1 | 1.4966 | 10 |
| *LOC103690013* | 0.0494 | 0.3484 | 0.217 | 0.156 | 1 | 1.391 | 10 |
| *Cxcl3* | 0.0019 | 0.6095 | 0.467 | 0.356 | 1 | 1.3118 | 10 |
| *Fabp5* | 0 | 1.4664 | 0.733 | 0.612 | 0.1283 | 1.1977 | 10 |
| *Basp1* | 8e-04 | 0.6985 | 0.55 | 0.474 | 1 | 1.1603 | 10 |
| *LOC684762* | 0.2906 | 0.3261 | 0.233 | 0.218 | 1 | 1.0688 | 10 |
| *Apln* | 1e-04 | 0.4207 | 0.9 | 0.85 | 1 | 1.0588 | 10 |
| *Casp4* | 0.0013 | 0.6827 | 0.517 | 0.492 | 1 | 1.0508 | 10 |
| *Trib3* | 0.0523 | 0.5147 | 0.417 | 0.398 | 1 | 1.0477 | 10 |
| *Atf3* | 0.2765 | 0.4073 | 0.267 | 0.255 | 1 | 1.0471 | 10 |
| *Smim3* | 0.3008 | 0.316 | 0.233 | 0.224 | 1 | 1.0402 | 10 |
| *Tubb4b* | 0 | 0.4779 | 0.9 | 0.873 | 1 | 1.0309 | 10 |
| *Ptprq* | 0 | 0.5162 | 1 | 0.978 | 0.461 | 1.0225 | 10 |
| *Psmd12* | 0 | 0.6847 | 0.95 | 0.93 | 0.0017 | 1.0215 | 10 |
| *Eif4ebp1* | 0 | 0.9137 | 0.983 | 0.963 | 0 | 1.0208 | 10 |
| *Arf4* | 0 | 0.4356 | 1 | 0.981 | 0.2482 | 1.0194 | 10 |
| *Rab33a* | 0.5564 | 0.2791 | 0.167 | 0.164 | 1 | 1.0183 | 10 |
| *Tubb6* | 1e-04 | 0.5431 | 0.95 | 0.934 | 1 | 1.0171 | 10 |
| *Nabp1* | 0.0422 | 0.6338 | 0.383 | 0.377 | 1 | 1.0159 | 10 |

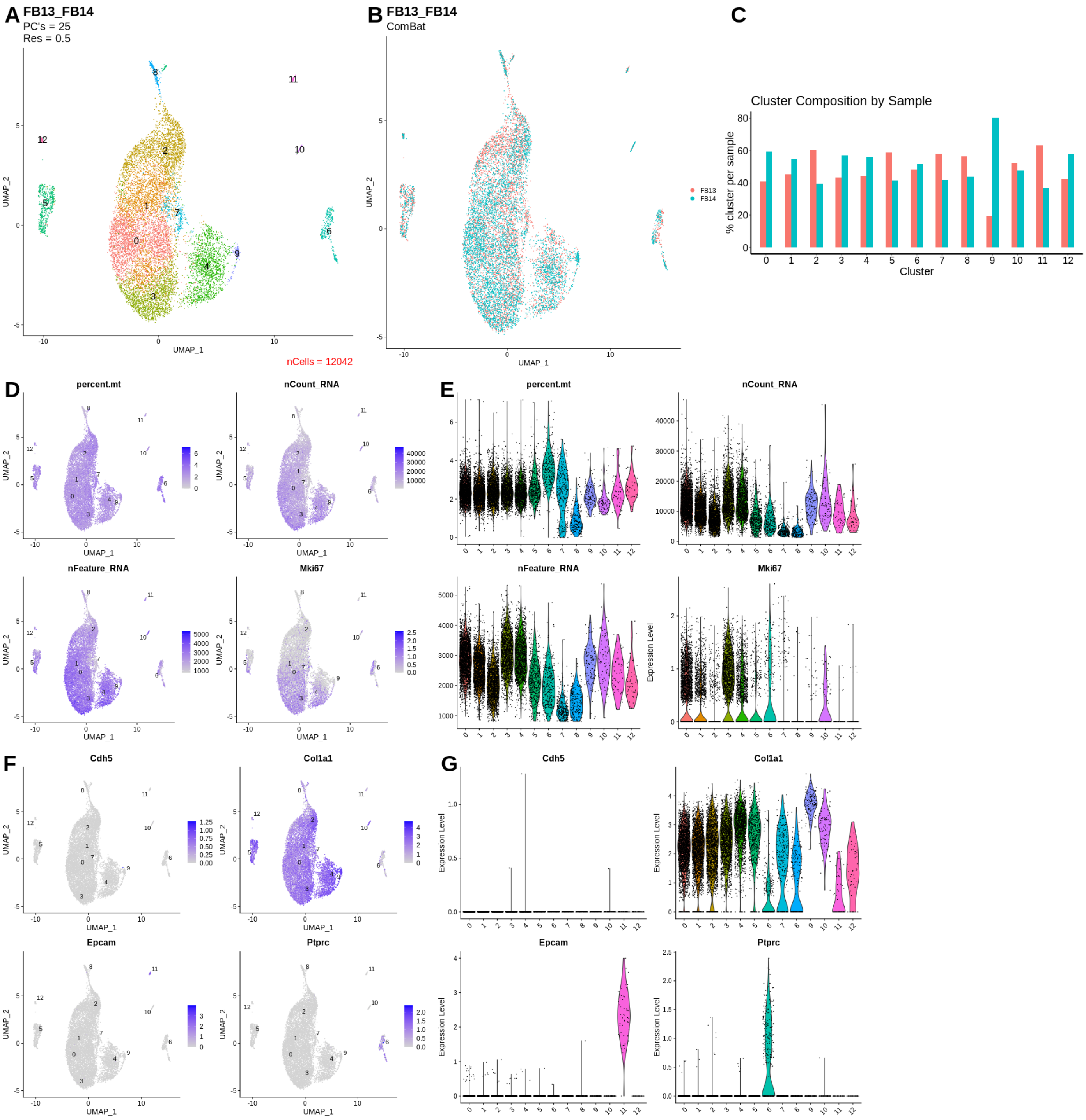


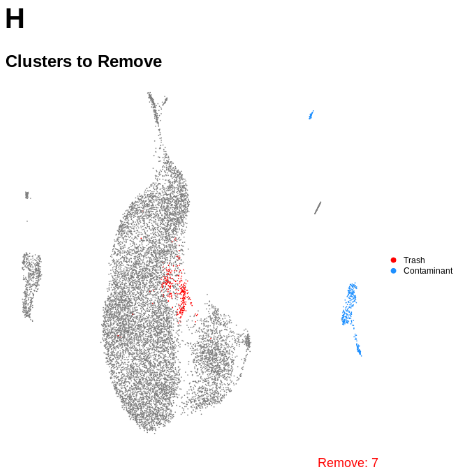


I

| | p_val | avg_log2FC | pct.1 | pct.2 | p_val_adj | ratio | cluster |
|---|---|---|---|---|---|---|---|
| *Cwf19l1* | 0 | 0.6973 | 0.222 | 0.077 | 0 | 2.8831 | 7 |
| *Nipsnap2* | 0 | 0.6913 | 0.239 | 0.124 | 0 | 1.9274 | 7 |
| *AABR07006894.1* | 2e-04 | 0.3823 | 0.102 | 0.056 | 1 | 1.8214 | 7 |
| *Slc39a1* | 0 | 1.8056 | 0.775 | 0.457 | 0 | 1.6958 | 7 |
| *Jund* | 0 | 1.2115 | 0.465 | 0.288 | 0 | 1.6146 | 7 |
| *Abcd3* | 0 | 0.5558 | 0.236 | 0.178 | 1 | 1.3258 | 7 |
| *Hs6st1* | 0 | 0.8017 | 0.366 | 0.292 | 5e-04 | 1.2534 | 7 |
| *Ccn1* | 0 | 1.8046 | 0.697 | 0.593 | 0 | 1.1754 | 7 |
| *Poldip3* | 0 | 0.8571 | 0.489 | 0.417 | 0 | 1.1727 | 7 |
| *Fitm2* | 0.1493 | 0.3486 | 0.123 | 0.108 | 1 | 1.1389 | 7 |
| *AABR07029605.1* | 0.2211 | 0.2834 | 0.102 | 0.09 | 1 | 1.1333 | 7 |
| *Rap1gds1* | 0 | 0.6998 | 0.317 | 0.281 | 0.7706 | 1.1281 | 7 |
| *Ube4b* | 0.0337 | 0.4995 | 0.197 | 0.18 | 1 | 1.0944 | 7 |
| *Tuba1b* | 0 | 0.2734 | 0.993 | 0.99 | 3e-04 | 1.003 | 7 |
| *Rps19l2* | 0 | 0.2897 | 0.965 | 0.994 | 0.057 | 0.9708 | 7 |
| *Vcl* | 0 | 1.4315 | 0.817 | 0.842 | 0 | 0.9703 | 7 |
| *Hnrnpa1* | 1e-04 | 0.2819 | 0.933 | 0.972 | 1 | 0.9599 | 7 |
| *LOC100359539* | 0 | 0.6155 | 0.588 | 0.659 | 0.0725 | 0.8923 | 7 |
| *Kpna1* | 0.7267 | 0.4197 | 0.18 | 0.204 | 1 | 0.8824 | 7 |
| *Cks1b* | 8e-04 | 0.2624 | 0.764 | 0.868 | 1 | 0.8802 | 7 |

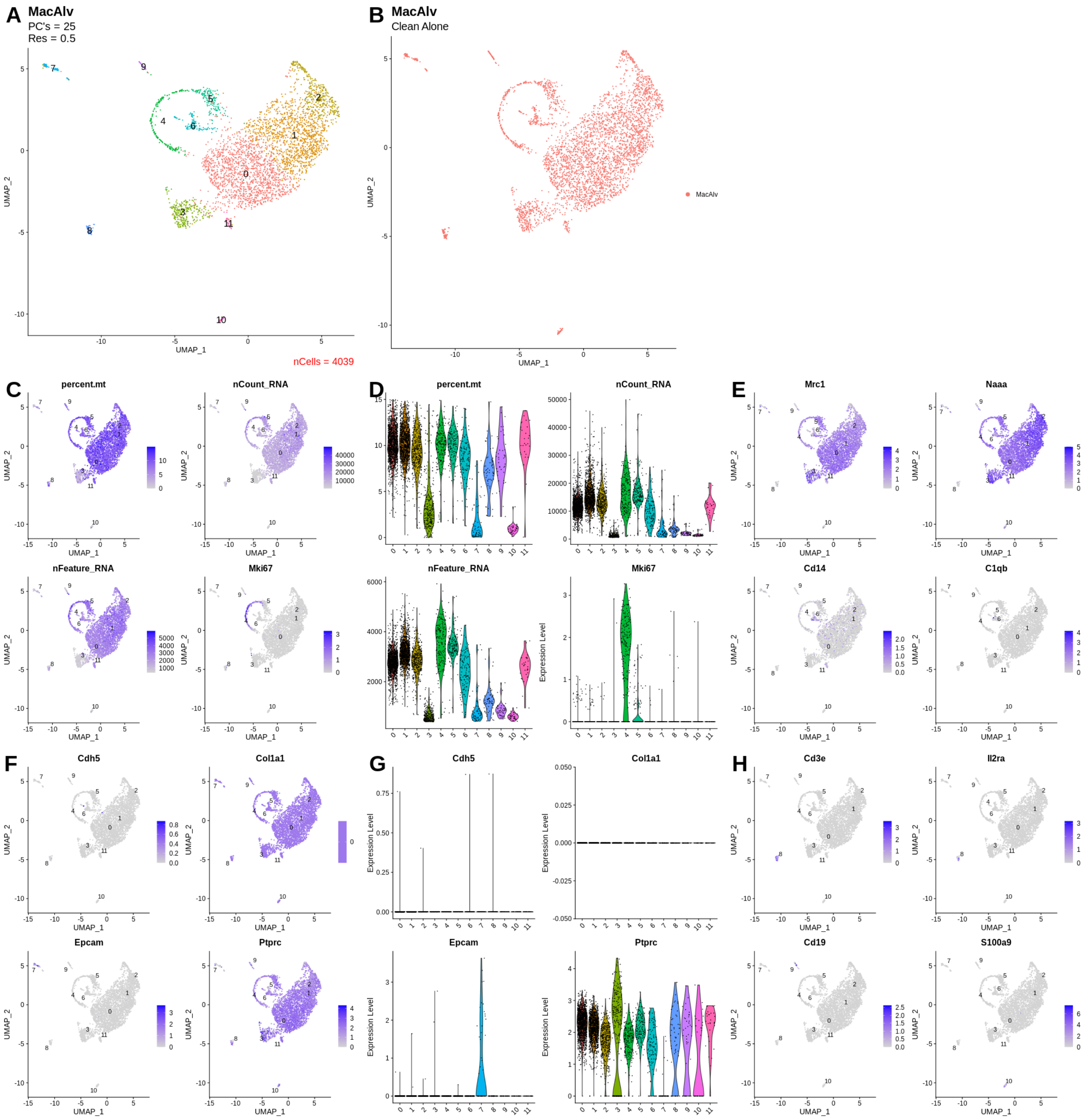


I

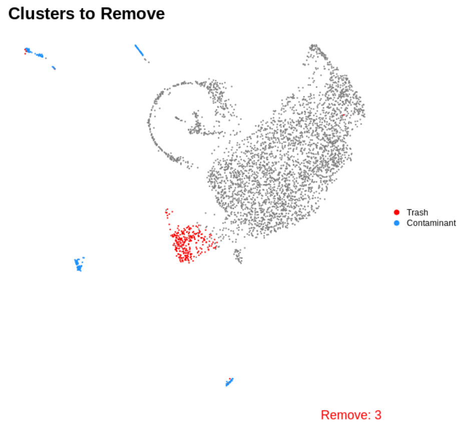


J

| | p_val | avg_log2FC | pct.1 | pct.2 | p_val_adj | ratio | cluster |
|---|---|---|---|---|---|---|---|
| *Hba-a1* | 0.038 | 2.311 | 0.134 | 0.107 | 1 | 1.2523 | 3 |
| *LOC100134871* | 0.1527 | 2.2435 | 0.13 | 0.116 | 1 | 1.1207 | 3 |
| *Cd9* | 0 | 0.7192 | 0.975 | 0.966 | 0 | 1.0093 | 3 |
| *S100a6* | 0 | 0.3953 | 0.982 | 0.974 | 0 | 1.0082 | 3 |
| *Lgals3* | 0 | 0.5443 | 0.971 | 0.974 | 0 | 0.9969 | 3 |
| *Lyz2* | 0 | 0.4079 | 0.989 | 0.993 | 0 | 0.996 | 3 |
| *B2m* | 0 | 0.4294 | 0.986 | 0.997 | 0 | 0.989 | 3 |
| *Rpl23* | 0 | 0.5341 | 0.982 | 0.996 | 0 | 0.9859 | 3 |
| *Ctsd* | 0.0037 | 0.2818 | 0.957 | 0.971 | 1 | 0.9856 | 3 |
| *LOC100911847* | 0 | 0.3079 | 0.971 | 0.997 | 0 | 0.9739 | 3 |
| *Rps16* | 0 | 0.5783 | 0.968 | 0.996 | 0 | 0.9719 | 3 |
| *Cstb* | 0 | 0.3238 | 0.949 | 0.982 | 0.0023 | 0.9664 | 3 |
| *Cd63* | 0 | 0.3076 | 0.939 | 0.975 | 1 | 0.9631 | 3 |
| *Rps11* | 0 | 0.2724 | 0.953 | 0.994 | 0 | 0.9588 | 3 |
| *Taldo1* | 0 | 0.4869 | 0.935 | 0.977 | 0 | 0.957 | 3 |
| *Rpl35* | 0 | 0.4534 | 0.949 | 0.995 | 0 | 0.9538 | 3 |
| *LOC108352650* | 0 | 0.3715 | 0.949 | 0.995 | 0.1962 | 0.9538 | 3 |
| *AY172581.9* | 0 | 2.0822 | 0.942 | 0.992 | 0 | 0.9496 | 3 |
| *AY172581.24* | 0 | 1.995 | 0.942 | 0.992 | 0 | 0.9496 | 3 |
| *Crip1* | 0 | 0.3116 | 0.928 | 0.978 | 1 | 0.9489 | 3 |

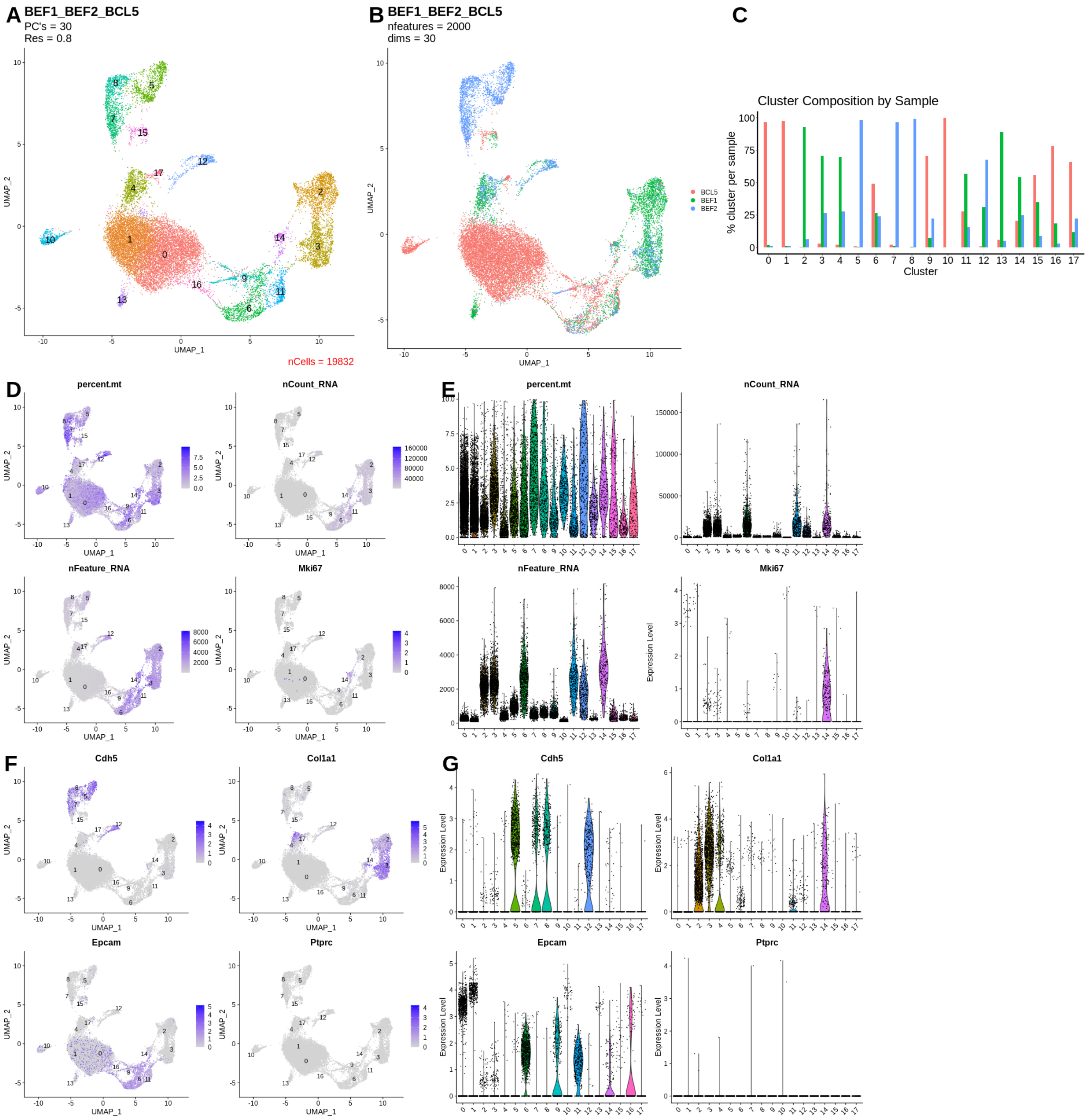


H

Clusters to Remove

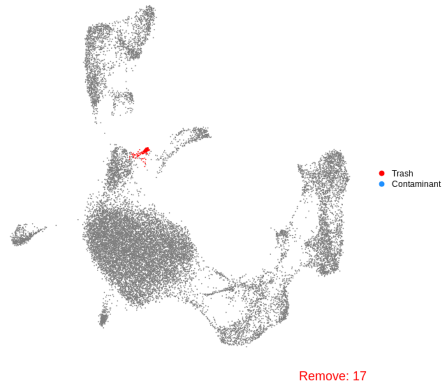


I

| | p_val | avg_log2FC | pct.1 | pct.2 | p_val_adj | ratio | cluster |
|---|---|---|---|---|---|---|---|
| *LOC100361025* | 0 | 4.4686 | 0.933 | 0.117 | 0 | 7.9744 | 17 |
| *Prmt1* | 0 | 4.422 | 0.904 | 0.114 | 0 | 7.9298 | 17 |
| *Ccn1* | 0.0136 | 1.1994 | 0.133 | 0.082 | 1 | 1.622 | 17 |
| *Ly6al* | 0.2864 | 0.4917 | 0.126 | 0.101 | 1 | 1.2475 | 17 |
| *LOC100360449* | 0.004 | 0.3148 | 0.807 | 0.701 | 1 | 1.1512 | 17 |
| *Rps3* | 0.0122 | 0.2586 | 0.815 | 0.708 | 1 | 1.1511 | 17 |
| *Rpl32* | 0 | 0.4214 | 0.726 | 0.641 | 1 | 1.1326 | 17 |
| *Rps11* | 0.0014 | 0.3703 | 0.756 | 0.668 | 1 | 1.1317 | 17 |
| *LOC100360087* | 0.0033 | 0.417 | 0.667 | 0.593 | 1 | 1.1248 | 17 |
| *S100a11* | 7e-04 | 0.3387 | 0.785 | 0.699 | 1 | 1.123 | 17 |
| *Gstt3* | 0.4129 | 0.9851 | 0.133 | 0.12 | 1 | 1.1083 | 17 |
| *Rps15a* | 2e-04 | 0.3159 | 0.844 | 0.766 | 1 | 1.1018 | 17 |
| *LOC100911847* | 0.0021 | 0.2764 | 0.911 | 0.828 | 1 | 1.1002 | 17 |
| *S100a6* | 8e-04 | 0.2639 | 0.97 | 0.889 | 1 | 1.0911 | 17 |
| *Rps17* | 1e-04 | 0.4667 | 0.733 | 0.672 | 1 | 1.0908 | 17 |
| *Clca4l* | 0.7204 | 0.2961 | 0.104 | 0.097 | 1 | 1.0722 | 17 |
| *Aqp5* | 0.5559 | 0.2902 | 0.23 | 0.215 | 1 | 1.0698 | 17 |
| *Rps7* | 8e-04 | 0.3838 | 0.763 | 0.714 | 1 | 1.0686 | 17 |
| *Rpl35* | 7e-04 | 0.2688 | 0.904 | 0.848 | 1 | 1.066 | 17 |
| *Rps27a* | 0.0044 | 0.6115 | 0.548 | 0.515 | 1 | 1.0641 | 17 |

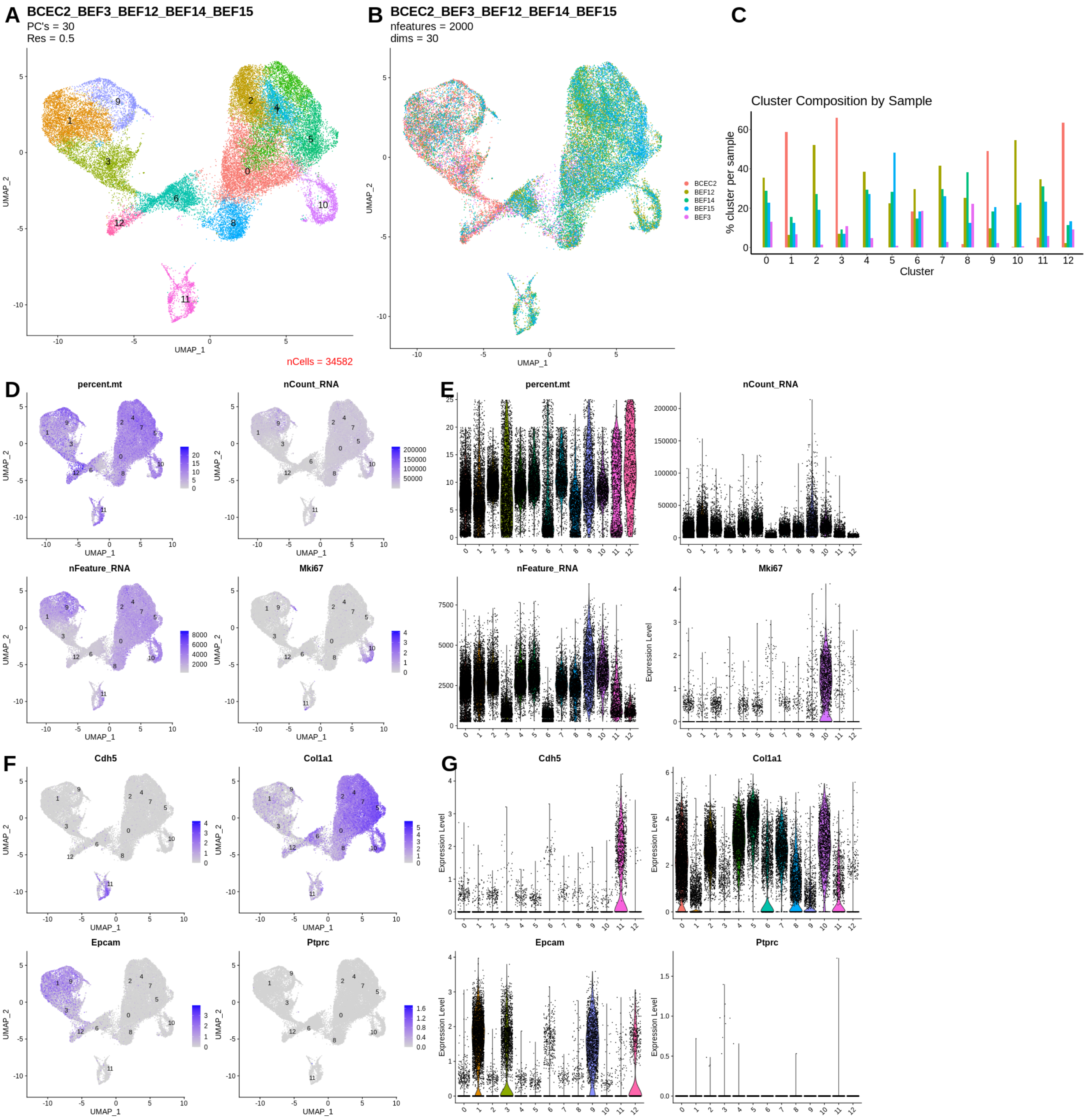


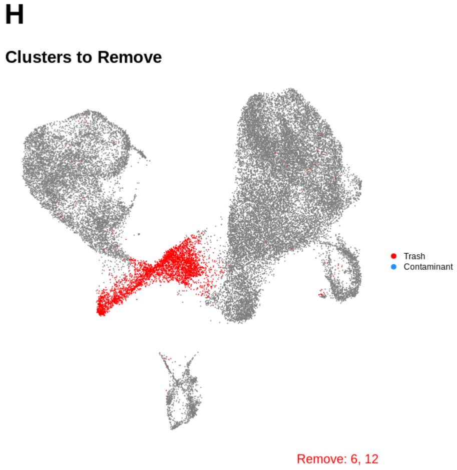


**I**

| | p_val | avg_log2FC | pct.1 | pct.2 | p_val_adj | ratio | cluster |
|---|---|---|---|---|---|---|---|
| *Mgp* | 0 | 0.3432 | 0.817 | 0.818 | 0 | 0.9988 | 6 |
| *AY172581.24* | 0 | 2.2122 | 0.97 | 0.998 | 0 | 0.9719 | 6 |
| *Mt2A* | 0 | 0.2999 | 0.736 | 0.786 | 5e-04 | 0.9364 | 6 |
| *Rpl26* | 0 | 0.2726 | 0.857 | 0.966 | 0.259 | 0.8872 | 6 |
| *Rpl36a* | 0 | 0.288 | 0.849 | 0.958 | 3e-04 | 0.8862 | 6 |
| *AY172581.9* | 0 | 1.6294 | 0.871 | 0.983 | 0 | 0.8861 | 6 |
| *Rpl21.3* | 0 | 0.3236 | 0.841 | 0.964 | 0 | 0.8724 | 6 |
| *Rpl32* | 0 | 0.3903 | 0.833 | 0.956 | 0 | 0.8713 | 6 |
| *Lgals1* | 0.5621 | 0.2572 | 0.611 | 0.726 | 1 | 0.8416 | 6 |
| *Rps27a* | 0.2206 | 0.2974 | 0.788 | 0.957 | 1 | 0.8234 | 6 |
| *Rpl30* | 0.3223 | 0.3006 | 0.77 | 0.951 | 1 | 0.8097 | 6 |
| *AABR07000398.1* | 0 | 0.769 | 0.603 | 0.843 | 0.0028 | 0.7153 | 6 |
| *LOC103693375* | 0.0028 | 0.2732 | 0.601 | 0.873 | 1 | 0.6884 | 6 |
| *Cst3* | 0 | 0.3455 | 0.518 | 0.764 | 3e-04 | 0.678 | 6 |
| *Rps19l2* | 0 | 0.321 | 0.578 | 0.86 | 0.1279 | 0.6721 | 6 |
| *LOC100359951* | 0 | 0.3311 | 0.594 | 0.903 | 0.1152 | 0.6578 | 6 |
| *AC128960.1* | 0 | 0.3168 | 0.547 | 0.866 | 0 | 0.6316 | 6 |
| *Rpl37* | 0 | 0.3484 | 0.503 | 0.826 | 0 | 0.609 | 6 |
| *Cryab* | 0 | 0.3013 | 0.125 | 0.217 | 0 | 0.576 | 6 |
| *LOC102549726* | 0 | 0.2889 | 0.421 | 0.786 | 0 | 0.5356 | 6 |

**J**

| | p_val | avg_log2FC | pct.1 | pct.2 | p_val_adj | ratio | cluster |
|---|---|---|---|---|---|---|---|
| *7SK.48* | 0 | 0.5557 | 0.102 | 0.002 | 0 | 51 | 12 |
| *Mroh6* | 0 | 0.6626 | 0.134 | 0.003 | 0 | 44.6667 | 12 |
| *7SK.261* | 0 | 0.6789 | 0.133 | 0.003 | 0 | 44.3333 | 12 |
| *Prnd* | 0 | 1.3392 | 0.128 | 0.003 | 0 | 42.6667 | 12 |
| *7SK.22* | 0 | 0.6315 | 0.123 | 0.003 | 0 | 41 | 12 |
| *7SK.242* | 0 | 0.6154 | 0.116 | 0.003 | 0 | 38.6667 | 12 |
| *Flvcr1* | 0 | 0.5289 | 0.109 | 0.003 | 0 | 36.3333 | 12 |
| *Arc* | 0 | 0.5219 | 0.104 | 0.003 | 0 | 34.6667 | 12 |
| *Ntng2* | 0 | 0.6328 | 0.126 | 0.004 | 0 | 31.5 | 12 |
| *LOC108348112* | 0 | 0.8868 | 0.187 | 0.006 | 0 | 31.1667 | 12 |
| *Plxnb1* | 0 | 0.7181 | 0.142 | 0.005 | 0 | 28.4 | 12 |
| *U6.82* | 0 | 1.2565 | 0.267 | 0.011 | 0 | 24.2727 | 12 |
| *Scarna2* | 0 | 1.0926 | 0.217 | 0.009 | 0 | 24.1111 | 12 |
| *AABR07044420.2* | 0 | 1.0416 | 0.216 | 0.009 | 0 | 24 | 12 |
| *Atg9b* | 0 | 1.9217 | 0.408 | 0.018 | 0 | 22.6667 | 12 |
| *Gm25541* | 0 | 2.1939 | 0.429 | 0.019 | 0 | 22.5789 | 12 |
| *Rpph1* | 0 | 1.6183 | 0.329 | 0.015 | 0 | 21.9333 | 12 |
| *Sowahb* | 0 | 0.6184 | 0.129 | 0.006 | 0 | 21.5 | 12 |
| *LOC100912564.4* | 0 | 0.6854 | 0.118 | 0.006 | 0 | 19.6667 | 12 |
| *RGD1562378* | 0 | 1.8892 | 0.391 | 0.02 | 0 | 19.55 | 12 |

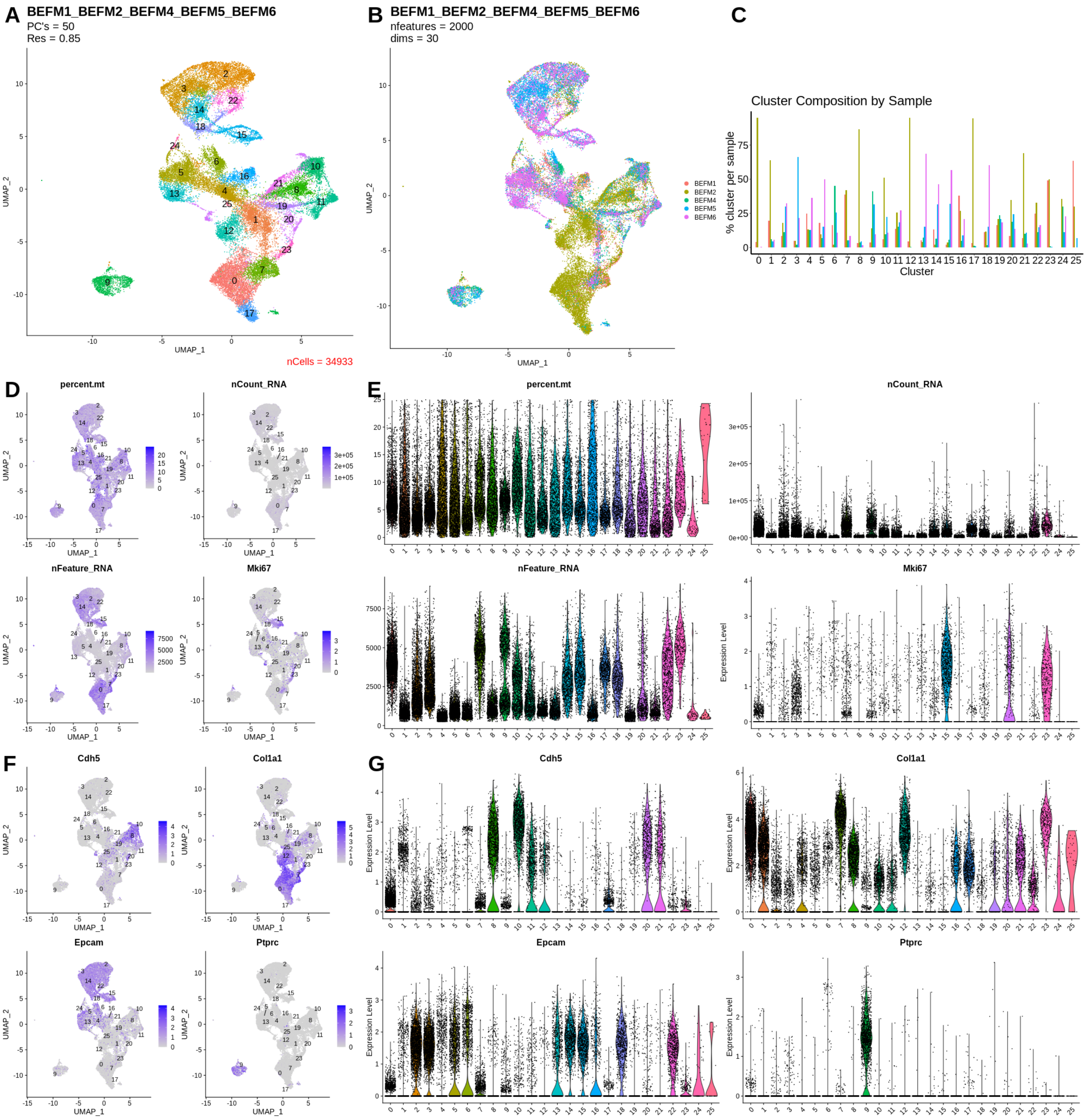


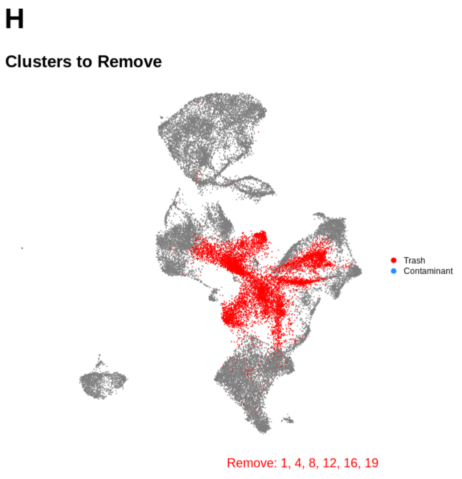


**I**

| | p_val | avg_log2FC | pct.1 | pct.2 | p_val_adj | ratio | cluster |
|---|---|---|---|---|---|---|---|
| *Tagln* | 0 | 1.5824 | 0.391 | 0.216 | 0 | 1.8102 | 1 |
| *Lgals1* | 0 | 1.591 | 0.832 | 0.463 | 0 | 1.797 | 1 |
| *Mgp* | 0 | 1.4649 | 0.864 | 0.513 | 0 | 1.6842 | 1 |
| *Col3a1* | 0 | 0.5508 | 0.739 | 0.441 | 0 | 1.6757 | 1 |
| *Cryab* | 0 | 1.3056 | 0.147 | 0.088 | 0 | 1.6705 | 1 |
| *Bgn* | 0 | 0.8542 | 0.642 | 0.386 | 0 | 1.6632 | 1 |
| *Col1a1* | 0 | 0.4168 | 0.778 | 0.478 | 0 | 1.6276 | 1 |
| *Gpx3* | 0 | 1.0295 | 0.524 | 0.345 | 0 | 1.5188 | 1 |

**J**

| | p_val | avg_log2FC | pct.1 | pct.2 | p_val_adj | ratio | cluster |
|---|---|---|---|---|---|---|---|
| *AY172581.21* | 0 | 0.3988 | 0.135 | 0.042 | 0 | 3.2143 | 4 |
| *AY172581.5* | 0 | 0.3932 | 0.123 | 0.039 | 0 | 3.1538 | 4 |
| *AY172581.3* | 0 | 0.6221 | 0.186 | 0.084 | 0 | 2.2143 | 4 |
| *AY172581.18* | 0 | 0.4453 | 0.148 | 0.074 | 0 | 2 | 4 |
| *AY172581.2* | 0 | 0.6005 | 0.167 | 0.09 | 0 | 1.8556 | 4 |
| *AY172581.12* | 0 | 0.335 | 0.101 | 0.058 | 0 | 1.7414 | 4 |
| *AY172581.23* | 0 | 0.4583 | 0.184 | 0.15 | 0 | 1.2267 | 4 |
| *Cxcl3* | 0 | 0.4081 | 0.774 | 0.631 | 0 | 1.2266 | 4 |

**K**

| | p_val | avg_log2FC | pct.1 | pct.2 | p_val_adj | ratio | cluster |
|---|---|---|---|---|---|---|---|
| *Hes2* | 0 | 0.58 | 0.109 | 0.003 | 0 | 36.3333 | 8 |
| *Arap3* | 0 | 1.3109 | 0.283 | 0.013 | 0 | 21.7692 | 8 |
| *7SK.113* | 0 | 0.6093 | 0.135 | 0.007 | 0 | 19.2857 | 8 |
| *Ildr2* | 0 | 0.5316 | 0.11 | 0.006 | 0 | 18.3333 | 8 |
| *7SK.405* | 0 | 0.7777 | 0.164 | 0.009 | 0 | 18.2222 | 8 |
| *7SK.35* | 0 | 0.4929 | 0.108 | 0.006 | 0 | 18 | 8 |
| *7SK.6* | 0 | 0.6224 | 0.14 | 0.008 | 0 | 17.5 | 8 |
| *7SK.321* | 0 | 0.9094 | 0.154 | 0.009 | 0 | 17.1111 | 8 |

**L**

| | p_val | avg_log2FC | pct.1 | pct.2 | p_val_adj | ratio | cluster |
|---|---|---|---|---|---|---|---|
| *7SK.184* | 0 | 0.6968 | 0.143 | 0.015 | 0 | 9.5333 | 12 |
| *7SK.76* | 0 | 0.6325 | 0.132 | 0.014 | 0 | 9.4286 | 12 |
| *7SK.51* | 0 | 0.601 | 0.132 | 0.014 | 0 | 9.4286 | 12 |
| *7SK.225* | 0 | 1.0204 | 0.217 | 0.025 | 0 | 8.68 | 12 |
| *7SK.72* | 0 | 0.8237 | 0.173 | 0.02 | 0 | 8.65 | 12 |
| *7SK.223* | 0 | 1.0534 | 0.224 | 0.026 | 0 | 8.6154 | 12 |
| *7SK.279* | 0 | 1.0633 | 0.232 | 0.027 | 0 | 8.5926 | 12 |
| *7SK.253* | 0 | 0.968 | 0.214 | 0.025 | 0 | 8.56 | 12 |

**M**

| | p_val | avg_log2FC | pct.1 | pct.2 | p_val_adj | ratio | cluster |
|---|---|---|---|---|---|---|---|
| *Atg9b* | 0 | 1.4996 | 0.284 | 0.019 | 0 | 14.9474 | 16 |
| *Terc* | 0 | 0.5125 | 0.103 | 0.016 | 0 | 6.4375 | 16 |
| *AABR07044420.2* | 0 | 0.6906 | 0.131 | 0.022 | 0 | 5.9545 | 16 |
| *Card10* | 0 | 0.6183 | 0.128 | 0.023 | 0 | 5.5652 | 16 |
| *Kcnk5* | 0 | 0.7597 | 0.129 | 0.024 | 0 | 5.375 | 16 |
| *Rap2b* | 0 | 0.4684 | 0.1 | 0.019 | 0 | 5.2632 | 16 |
| *Scarna2* | 0 | 0.846 | 0.168 | 0.033 | 0 | 5.0909 | 16 |
| *AABR07050487.1* | 0 | 1.0691 | 0.242 | 0.049 | 0 | 4.9388 | 16 |

**N**

| | p_val | avg_log2FC | pct.1 | pct.2 | p_val_adj | ratio | cluster |
|---|---|---|---|---|---|---|---|
| *Sncg* | 0 | 1.3526 | 0.206 | 0.025 | 0 | 8.24 | 19 |
| *Esm1* | 0 | 2.8525 | 0.545 | 0.139 | 0 | 3.9209 | 19 |
| *Ptprq* | 0 | 1.0266 | 0.246 | 0.09 | 0 | 2.7333 | 19 |
| *Srgn* | 0 | 1.667 | 0.362 | 0.148 | 0 | 2.4459 | 19 |
| *Gsta1* | 0 | 1.552 | 0.35 | 0.151 | 0 | 2.3179 | 19 |
| *Procr* | 0 | 1.8766 | 0.42 | 0.195 | 0 | 2.1538 | 19 |
| *Ccn3* | 0 | 1.6175 | 0.424 | 0.209 | 0 | 2.0287 | 19 |
| *Ecscr* | 0 | 0.8066 | 0.169 | 0.094 | 0 | 1.7979 | 19 |

### III. Cleaning sample objects individually

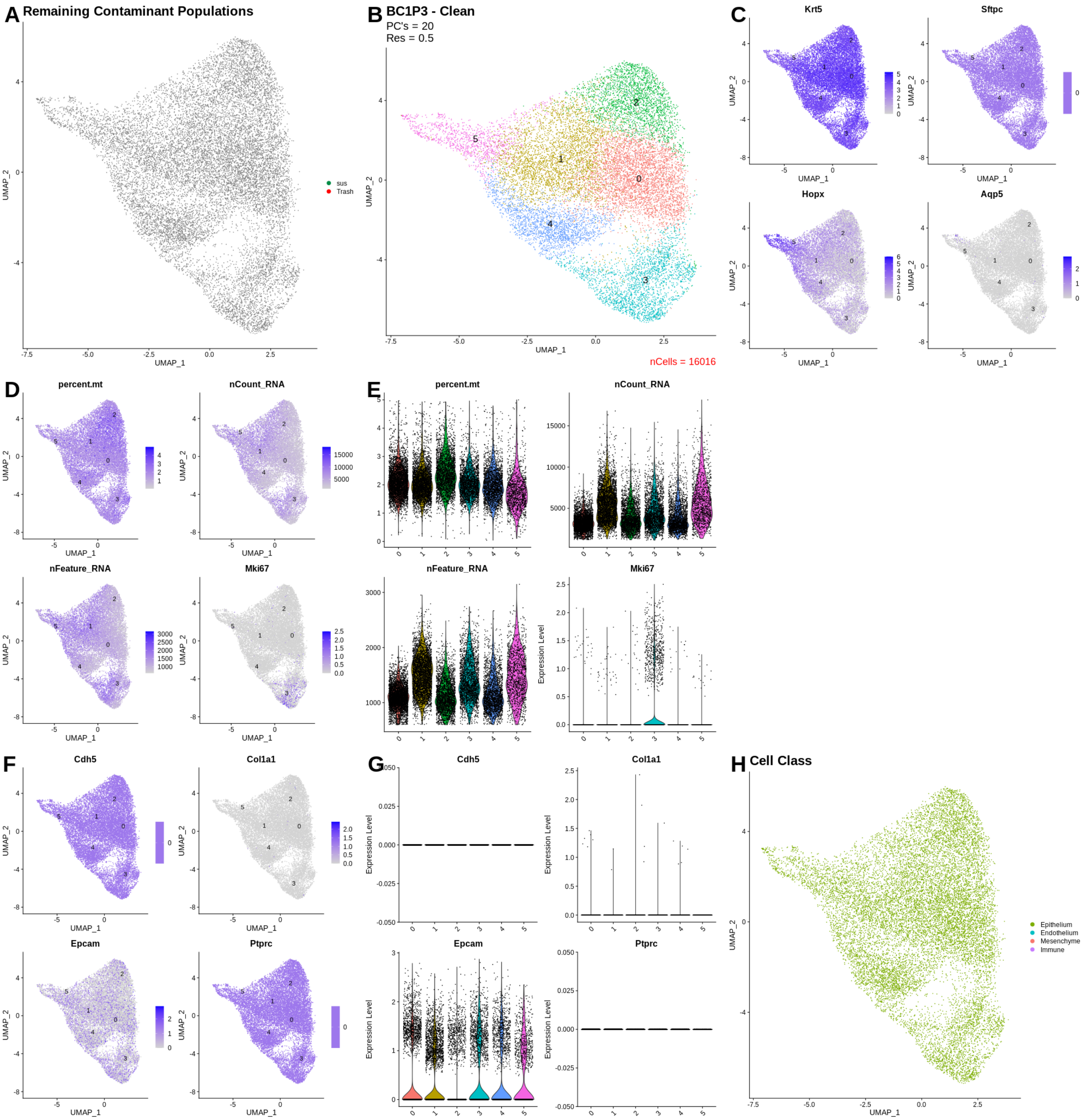
A Remaining Contaminant Populations
sus
Trash
B BC1P3 - Clean
PC's = 20
Res = 0.5
nCells = 16016
C
Krt5
Sftpc
Hopx
Aqp5
D
percent.mt
nCount_RNA
nFeature_RNA
Mki67
E
F
Cdh5
Col1a1
Epcam
Ptprc
G
H Cell Class
Epithelium
Endothelium
Mesenchyme
Immune
UMAP_1
UMAP_2
Expression Level

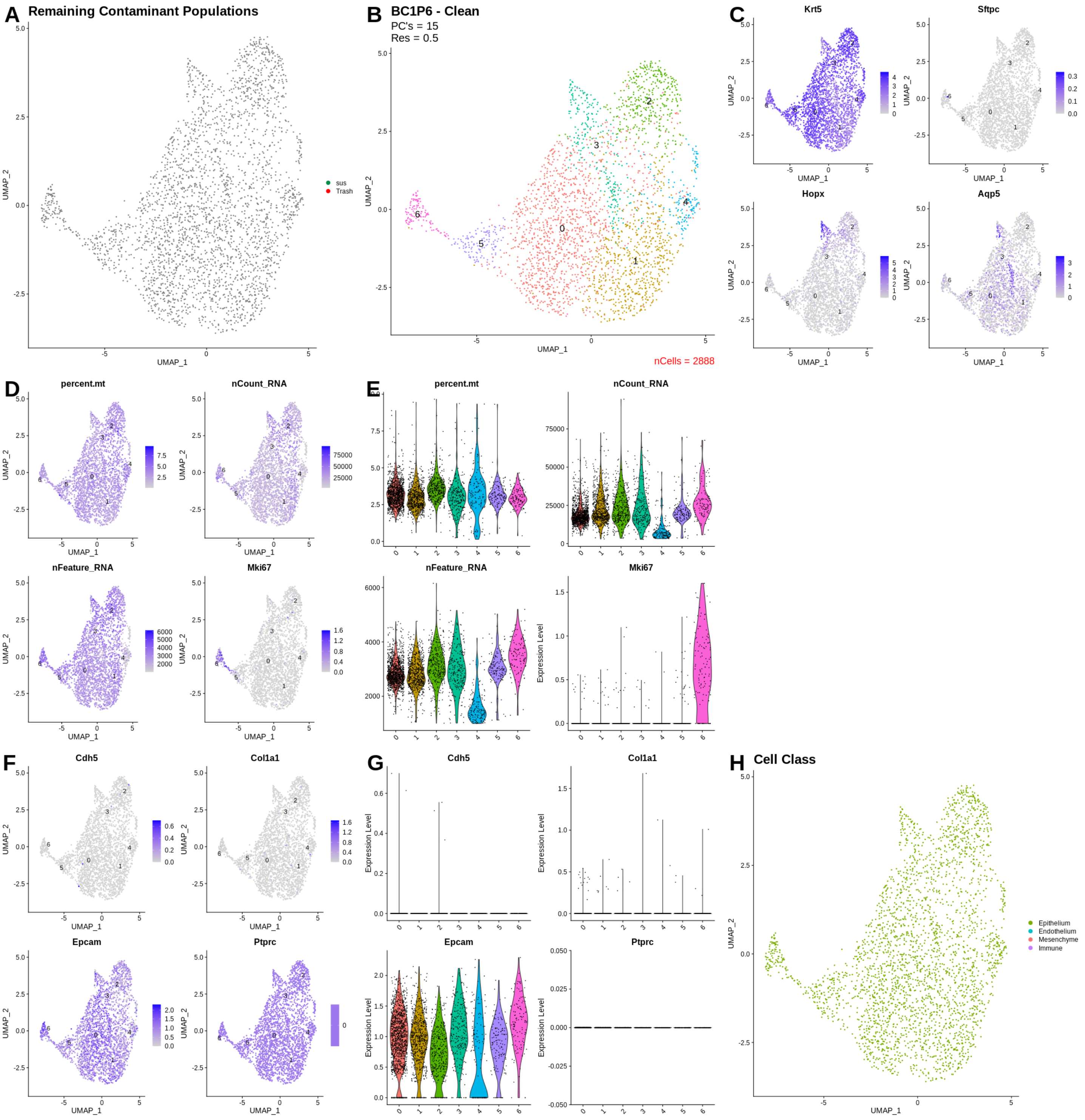
A Remaining Contaminant Populations
B BC1P6 - Clean
PC's = 15
Res = 0.5
nCells = 2888
C
Krt5
Sftpc
Hopx
Aqp5
D
percent.mt
nCount_RNA
nFeature_RNA
Mki67
E
F
Cdh5
Col1a1
Epcam
Ptprc
G
H Cell Class
Epithelium
Endothelium
Mesenchyme
Immune
sus
Trash
UMAP_1
UMAP_2
Expression Level

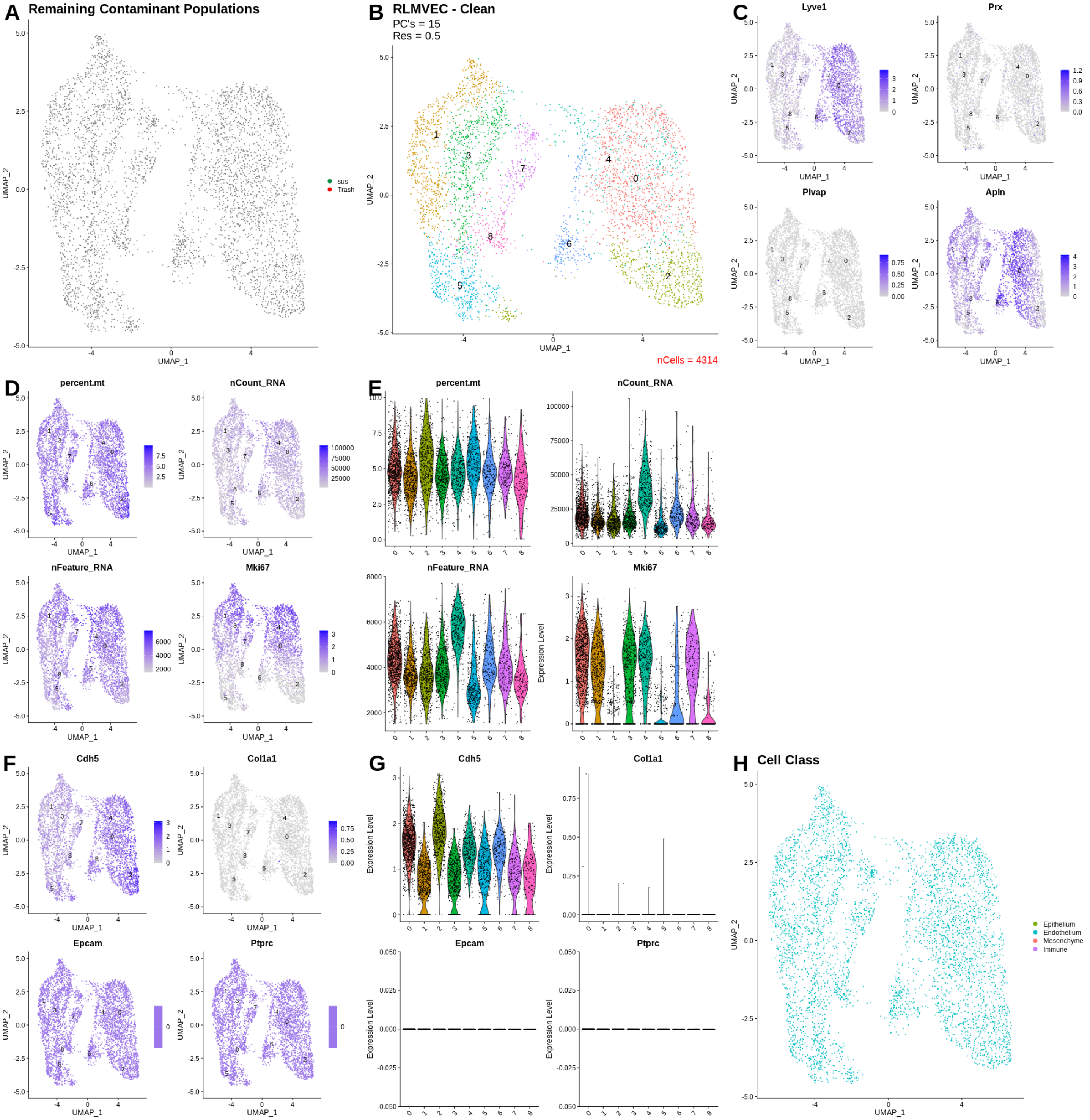
A Remaining Contaminant Populations
sus
Trash
B RLMVEC - Clean
PC's = 15
Res = 0.5
nCells = 4314
C
Lyve1
Prx
Plvap
Apln
D
percent.mt
nCount_RNA
nFeature_RNA
Mki67
E
percent.mt
nCount_RNA
nFeature_RNA
Mki67
F
Cdh5
Col1a1
Epcam
Ptprc
G
Cdh5
Col1a1
Epcam
Ptprc
H Cell Class
Epithelium
Endothelium
Mesenchyme
Immune
UMAP_1
UMAP_2
Expression Level

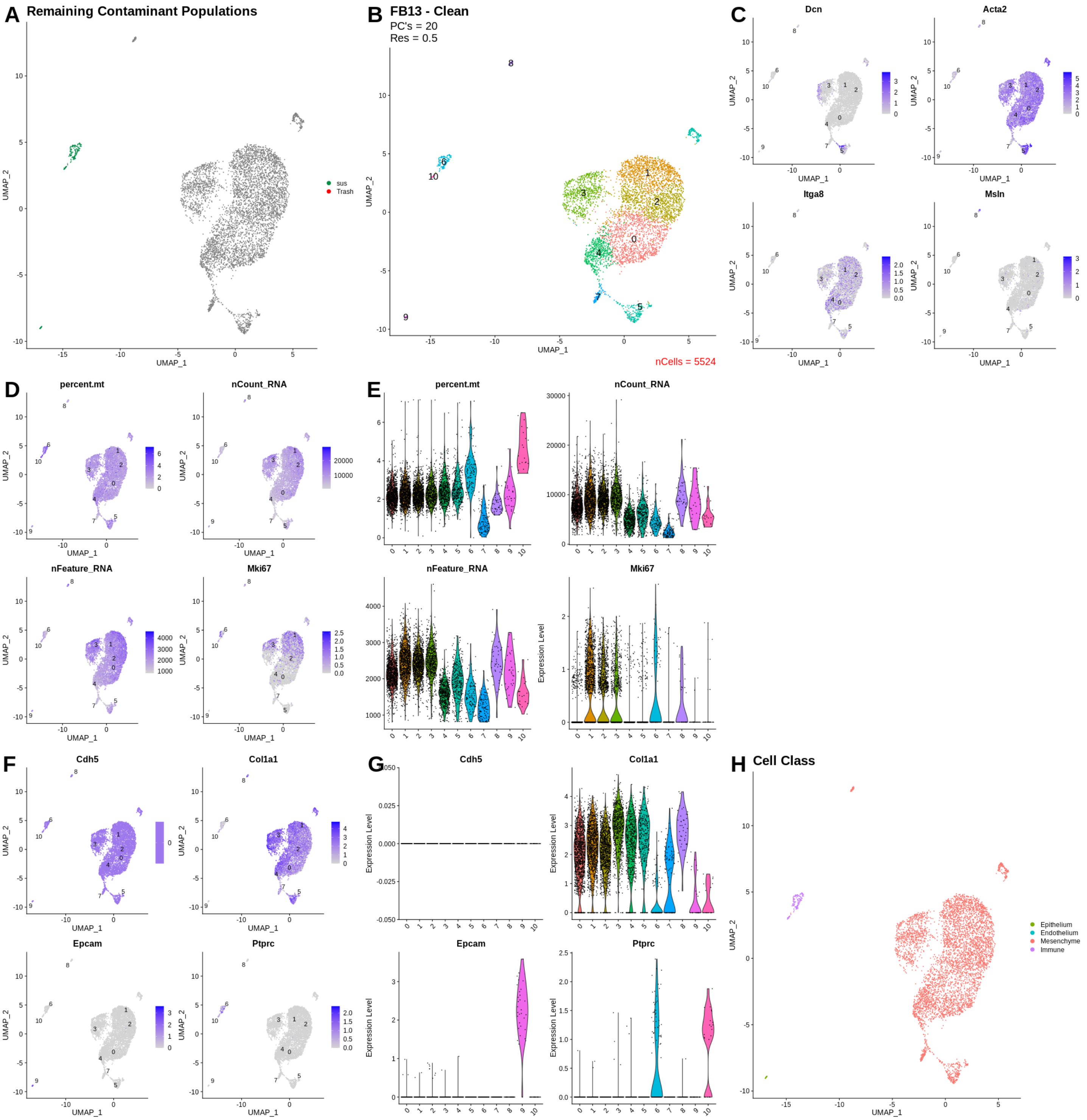

A Remaining Contaminant Populations
sus
Trash
UMAP_1
UMAP_2
B FB13 - Clean
PC's = 20
Res = 0.5
nCells = 5524
C
Dcn
Acta2
Itga8
Msln
D
percent.mt
nCount_RNA
nFeature_RNA
Mki67
E
F
Cdh5
Col1a1
Epcam
Ptprc
G
Expression Level
H Cell Class
Epithelium
Endothelium
Mesenchyme
Immune

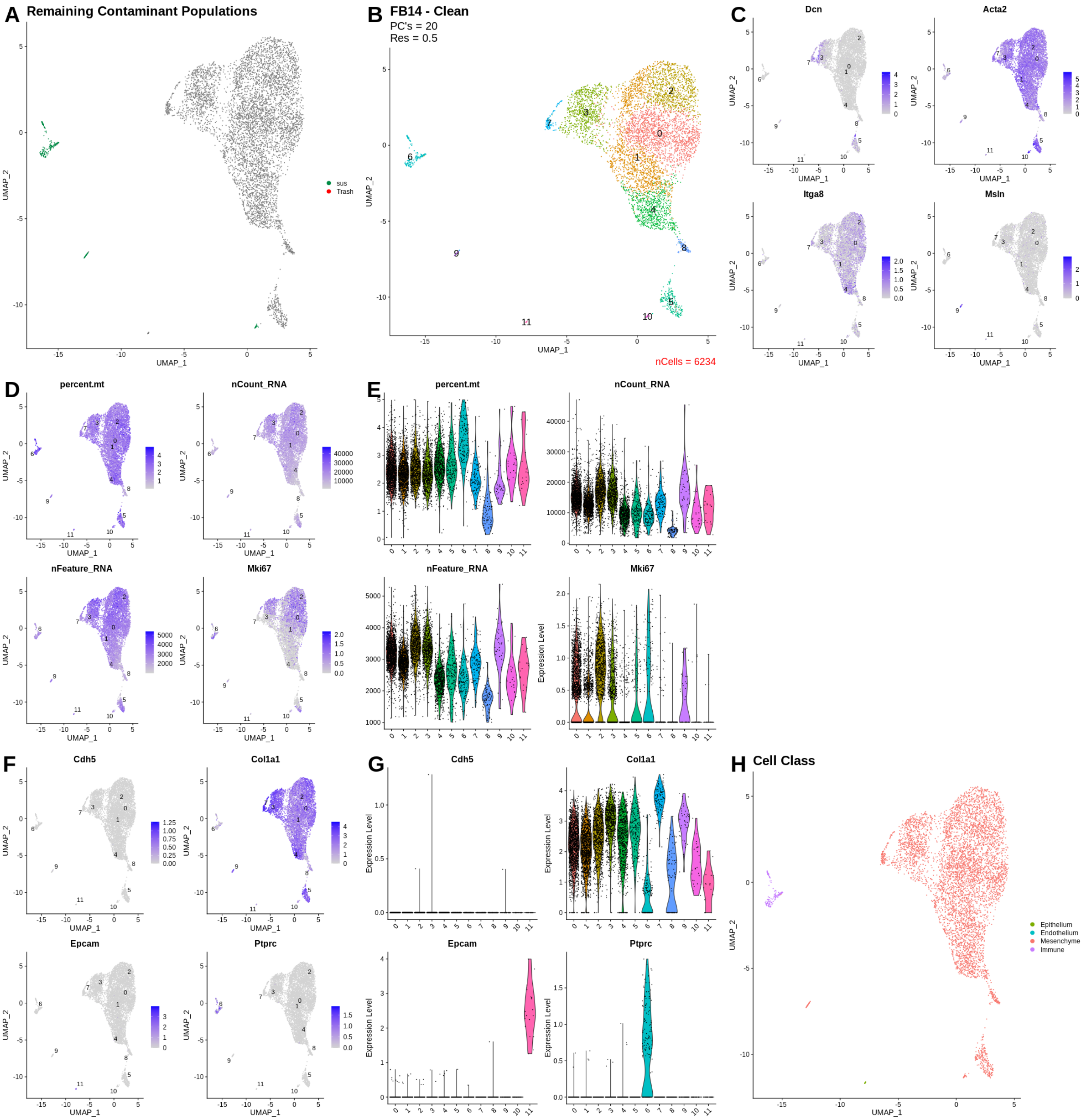
A Remaining Contaminant Populations
sus
Trash
B FB14 - Clean
PC's = 20
Res = 0.5
nCells = 6234
C
Dcn
Acta2
Itga8
Msln
D
percent.mt
nCount_RNA
nFeature_RNA
Mki67
E
F
Cdh5
Col1a1
Epcam
Ptprc
G
H Cell Class
Epithelium
Endothelium
Mesenchyme
Immune
UMAP_1
UMAP_2
Expression Level

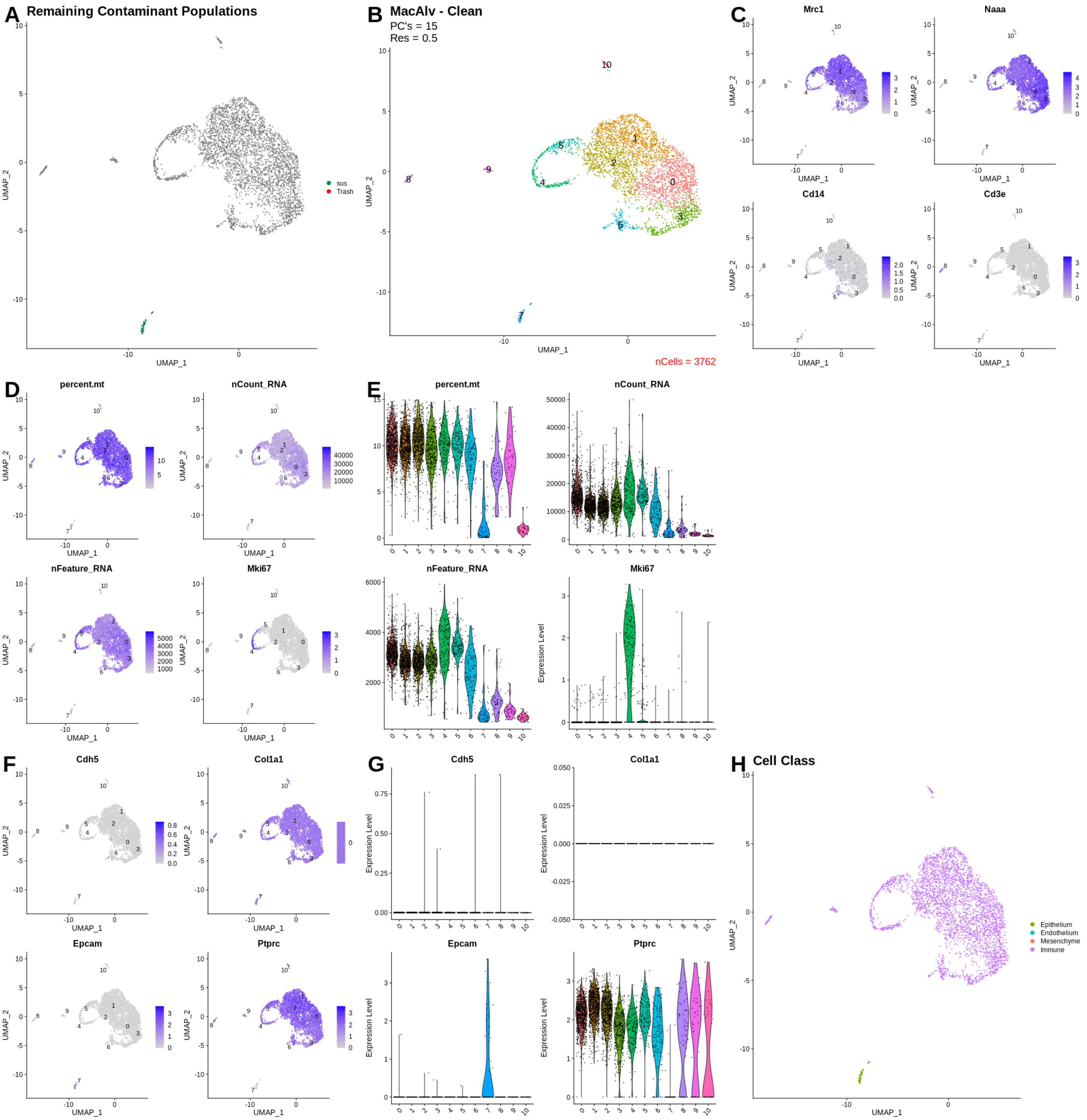

A Remaining Contaminant Populations
sus
Trash
B MacAlv - Clean
PC's = 15
Res = 0.5
nCells = 3762
C
Mrc1
Naaa
Cd14
Cd3e
D
percent.mt
nCount_RNA
nFeature_RNA
Mki67
E
percent.mt
nCount_RNA
nFeature_RNA
Mki67
F
Cdh5
Col1a1
Epcam
Ptprc
G
Cdh5
Col1a1
Epcam
Ptprc
H Cell Class
Epithelium
Endothelium
Mesenchyme
Immune
UMAP_1
UMAP_2
Expression Level

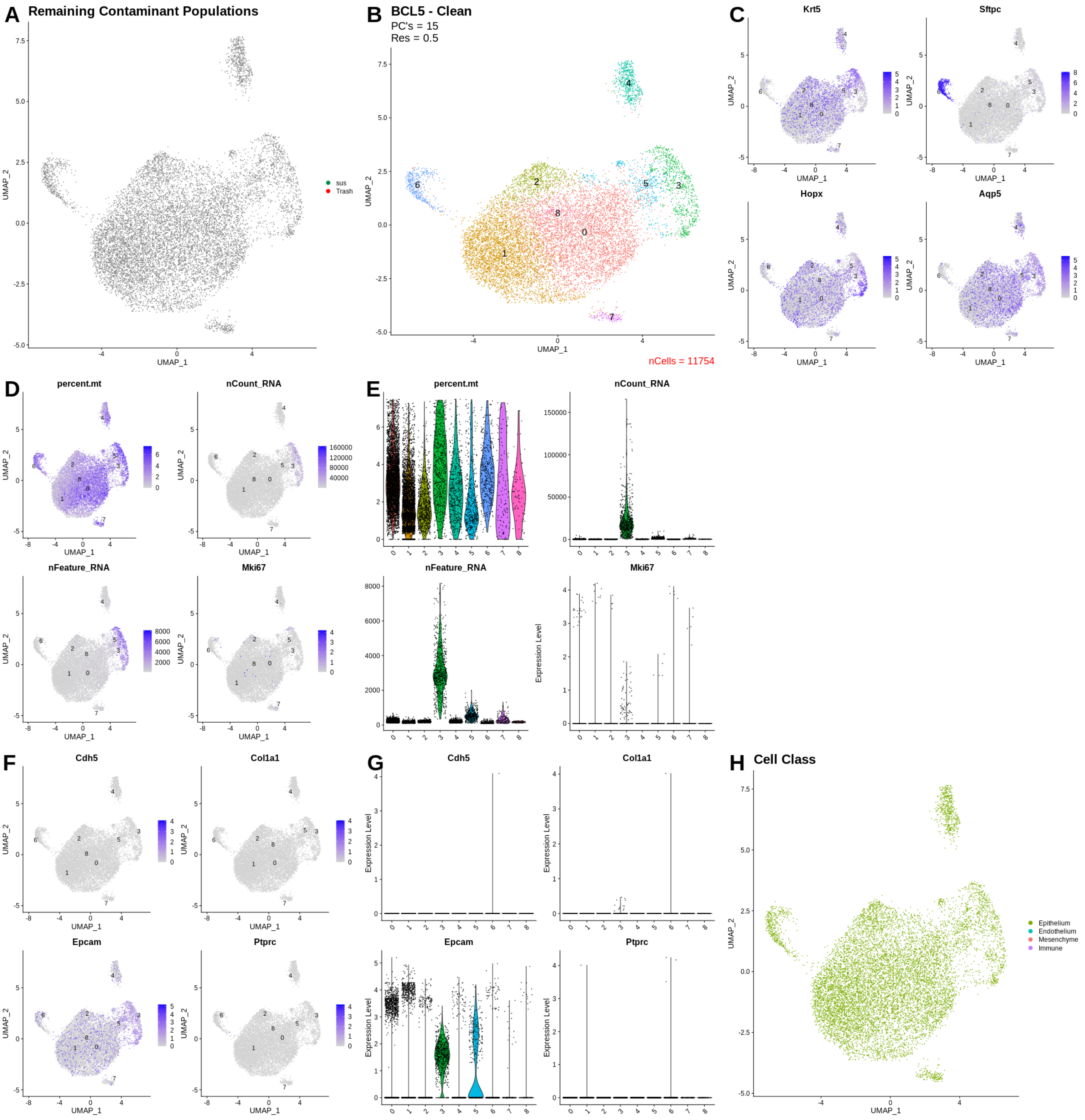
A
Remaining Contaminant Populations
sus
Trash
UMAP_1
UMAP_2
B
BCL5 - Clean
PC's = 15
Res = 0.5
nCells = 11754
C
Krt5
Sftpc
Hopx
Aqp5
D
percent.mt
nCount_RNA
nFeature_RNA
Mki67
E
percent.mt
nCount_RNA
nFeature_RNA
Mki67
Expression Level
F
Cdh5
Col1a1
Epcam
Ptprc
G
Cdh5
Col1a1
Epcam
Ptprc
H
Cell Class
Epithelium
Endothelium
Mesenchyme
Immune

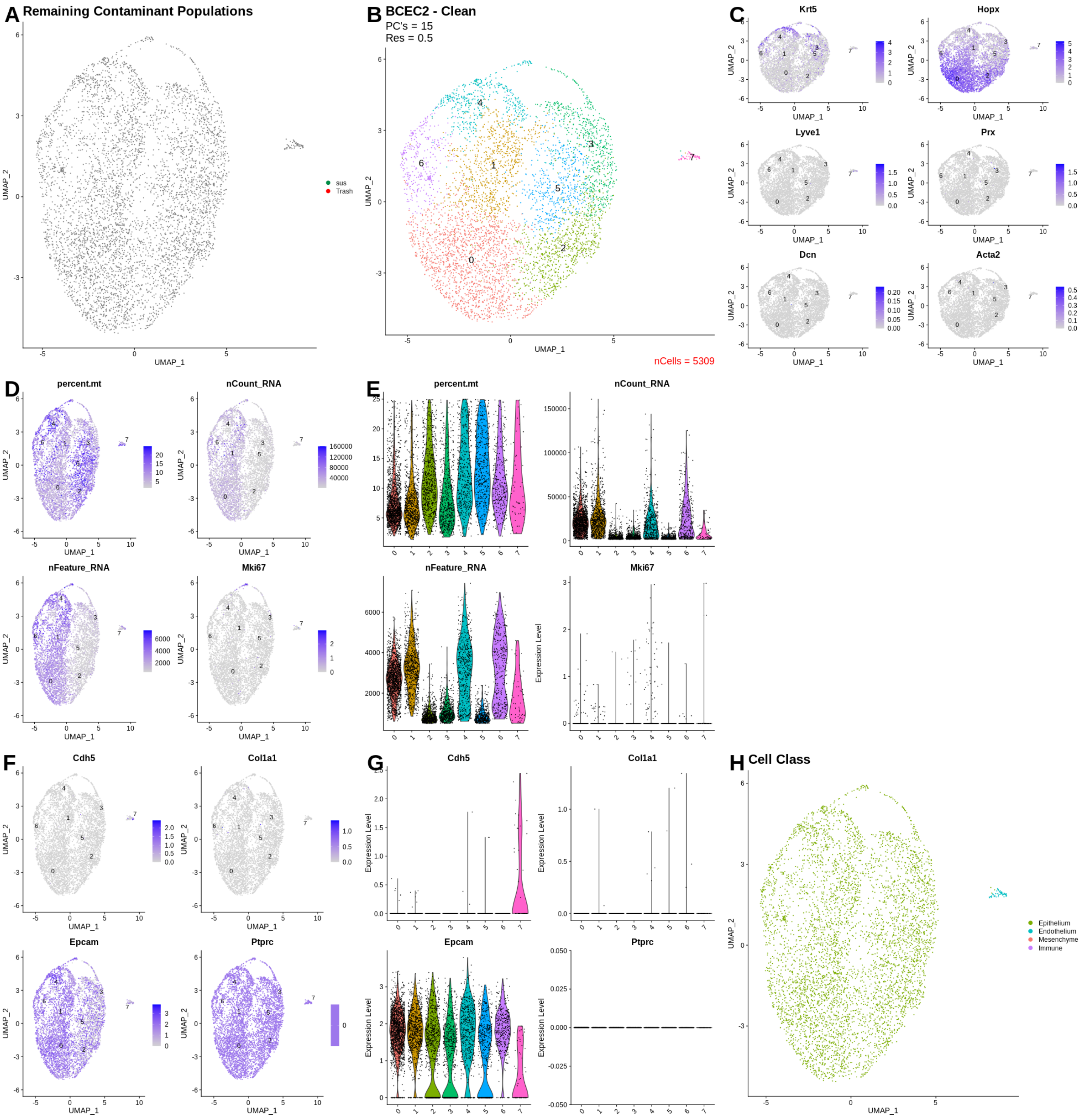
A Remaining Contaminant Populations
sus
Trash
UMAP_1
UMAP_2
B BCEC2 - Clean
PC's = 15
Res = 0.5
nCells = 5309
C
Krt5
Hopx
Lyve1
Prx
Dcn
Acta2
D
percent.mt
nCount_RNA
nFeature_RNA
Mki67
E
Expression Level
F
Cdh5
Col1a1
Epcam
Ptprc
G
H Cell Class
Epithelium
Endothelium
Mesenchyme
Immune

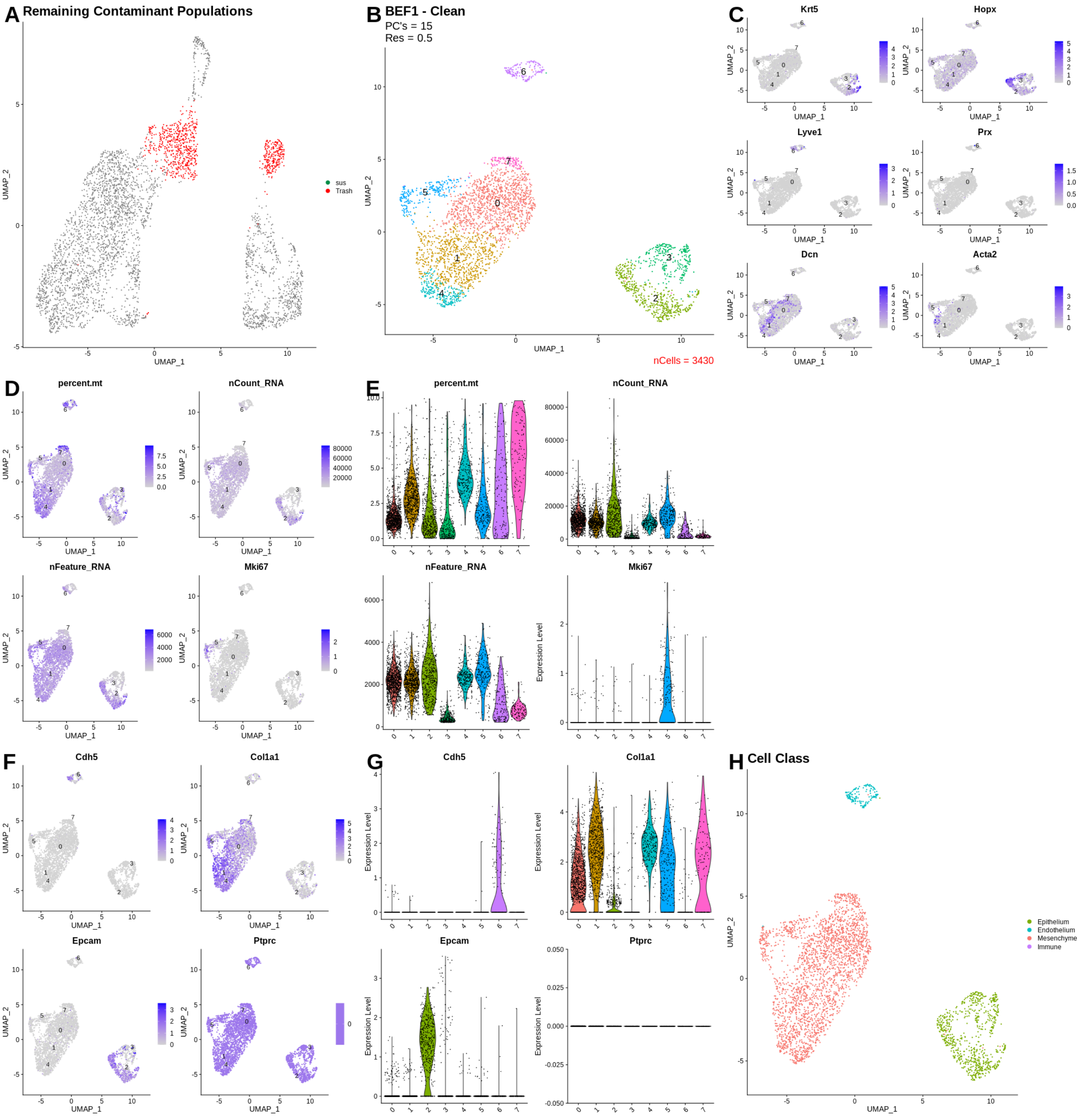
A
Remaining Contaminant Populations
sus
Trash
UMAP_1
UMAP_2
B
BEF1 - Clean
PC's = 15
Res = 0.5
nCells = 3430
C
Krt5
Hopx
Lyve1
Prx
Dcn
Acta2
D
percent.mt
nCount_RNA
nFeature_RNA
Mki67
E
Expression Level
F
Cdh5
Col1a1
Epcam
Ptprc
G
H
Cell Class
Epithelium
Endothelium
Mesenchyme
Immune

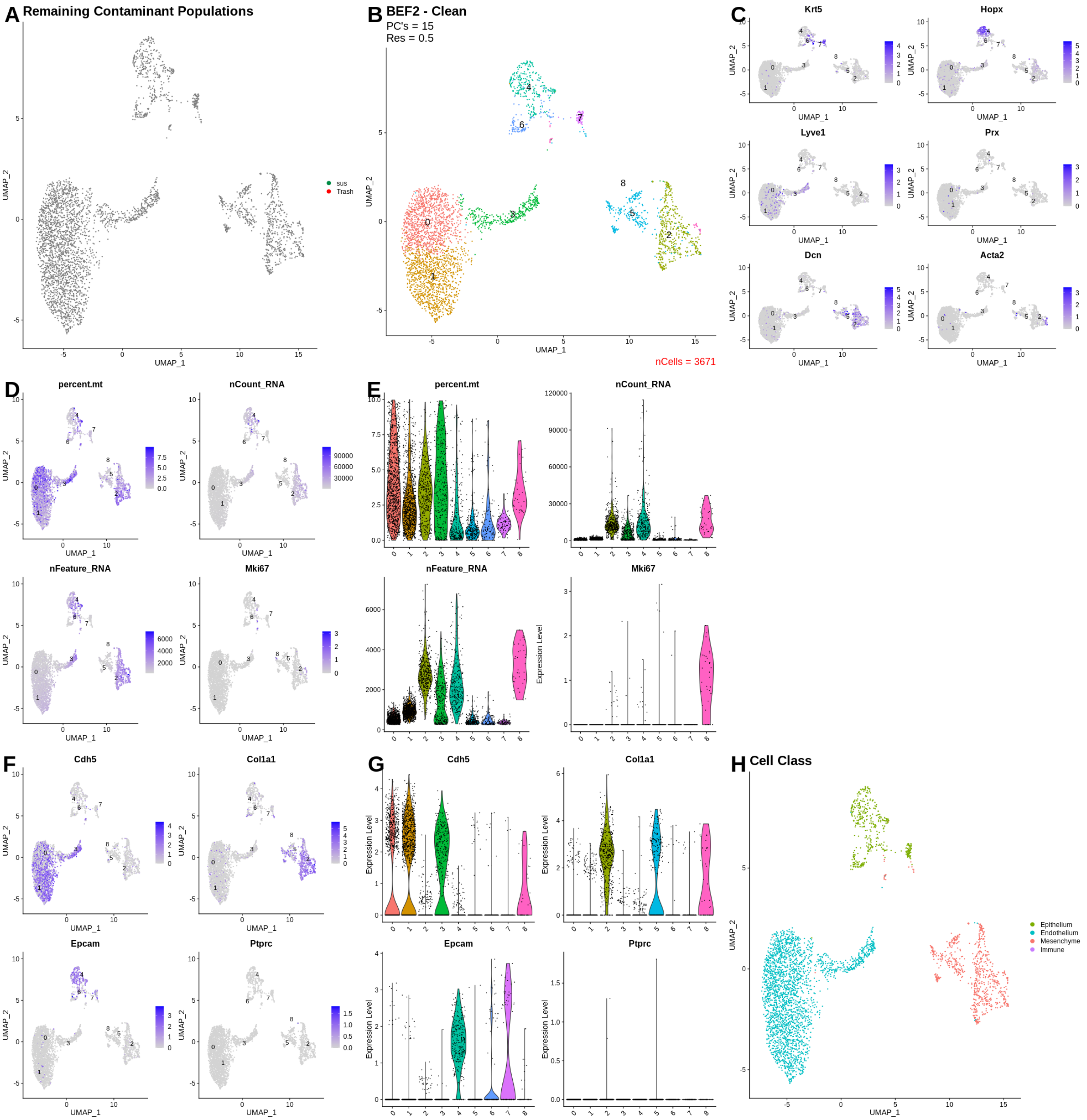

A
Remaining Contaminant Populations
sus
Trash
UMAP_1
UMAP_2
B
BEF2 - Clean
PC's = 15
Res = 0.5
nCells = 3671
C
Krt5
Hopx
Lyve1
Prx
Dcn
Acta2
D
percent.mt
nCount_RNA
nFeature_RNA
Mki67
E
percent.mt
nCount_RNA
nFeature_RNA
Mki67
Expression Level
F
Cdh5
Col1a1
Epcam
Ptprc
G
Cdh5
Col1a1
Epcam
Ptprc
Expression Level
H
Cell Class
Epithelium
Endothelium
Mesenchyme
Immune

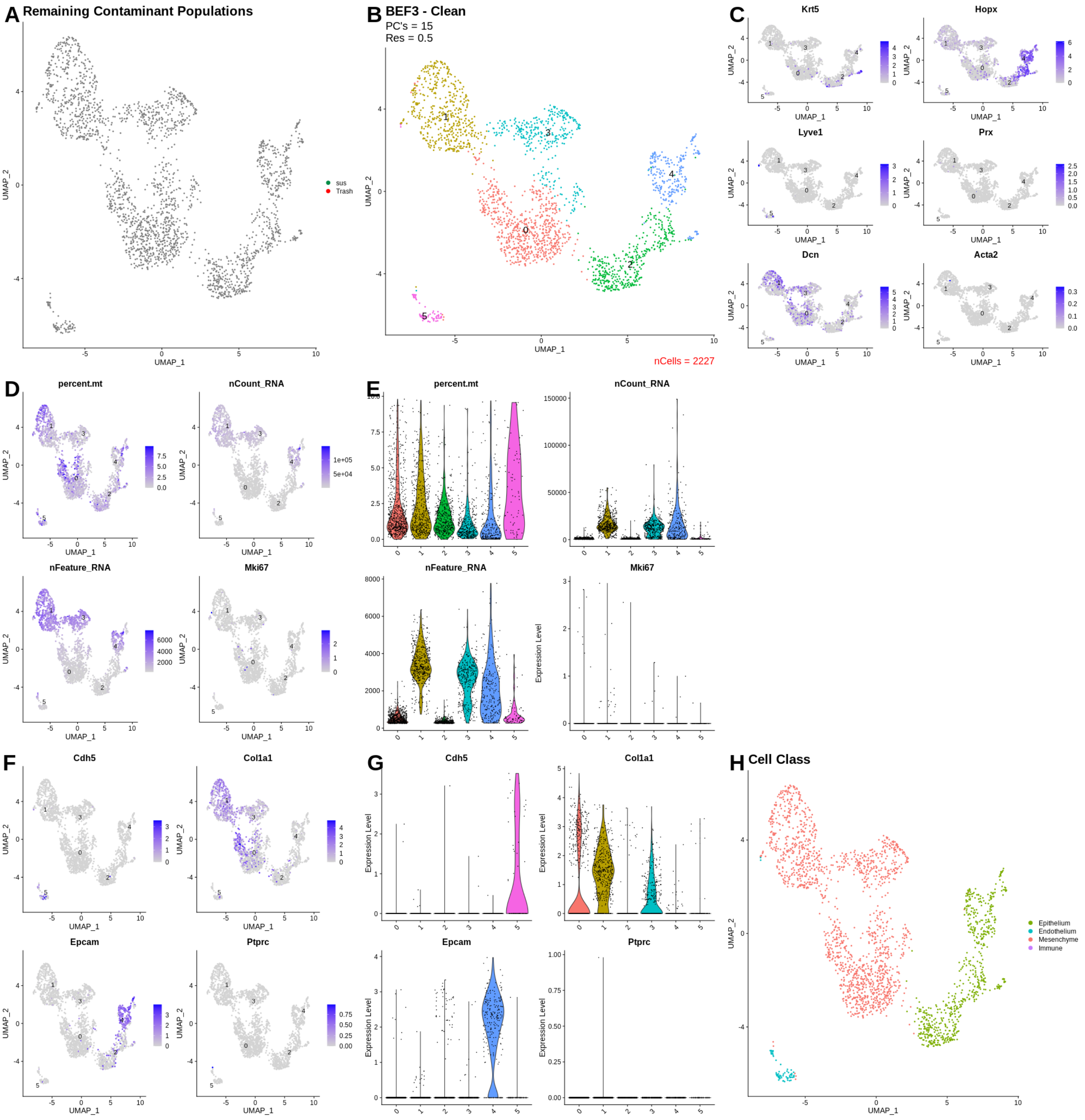
A
Remaining Contaminant Populations
sus
Trash
UMAP_1
UMAP_2
B
BEF3 - Clean
PC's = 15
Res = 0.5
nCells = 2227
C
Krt5
Hopx
Lyve1
Prx
Dcn
Acta2
D
percent.mt
nCount_RNA
nFeature_RNA
Mki67
E
percent.mt
nCount_RNA
nFeature_RNA
Mki67
Expression Level
F
Cdh5
Col1a1
Epcam
Ptprc
G
Cdh5
Col1a1
Epcam
Ptprc
H
Cell Class
Epithelium
Endothelium
Mesenchyme
Immune

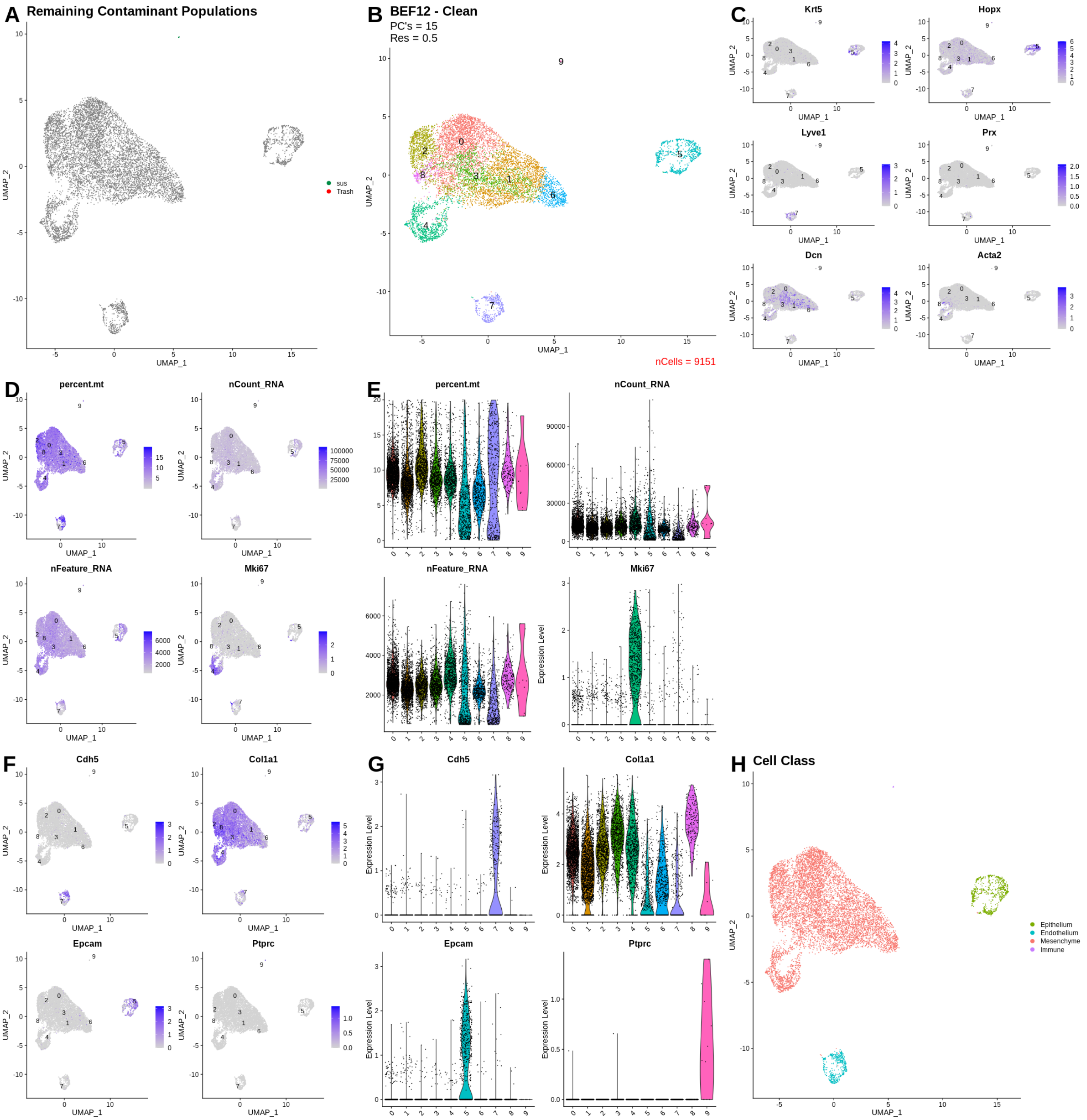
A Remaining Contaminant Populations
sus
Trash
UMAP_1
UMAP_2
B BEF12 - Clean
PC's = 15
Res = 0.5
nCells = 9151
C
Krt5
Hopx
Lyve1
Prx
Dcn
Acta2
D
percent.mt
nCount_RNA
nFeature_RNA
Mki67
E
Expression Level
F
Cdh5
Col1a1
Epcam
Ptprc
G
H Cell Class
Epithelium
Endothelium
Mesenchyme
Immune

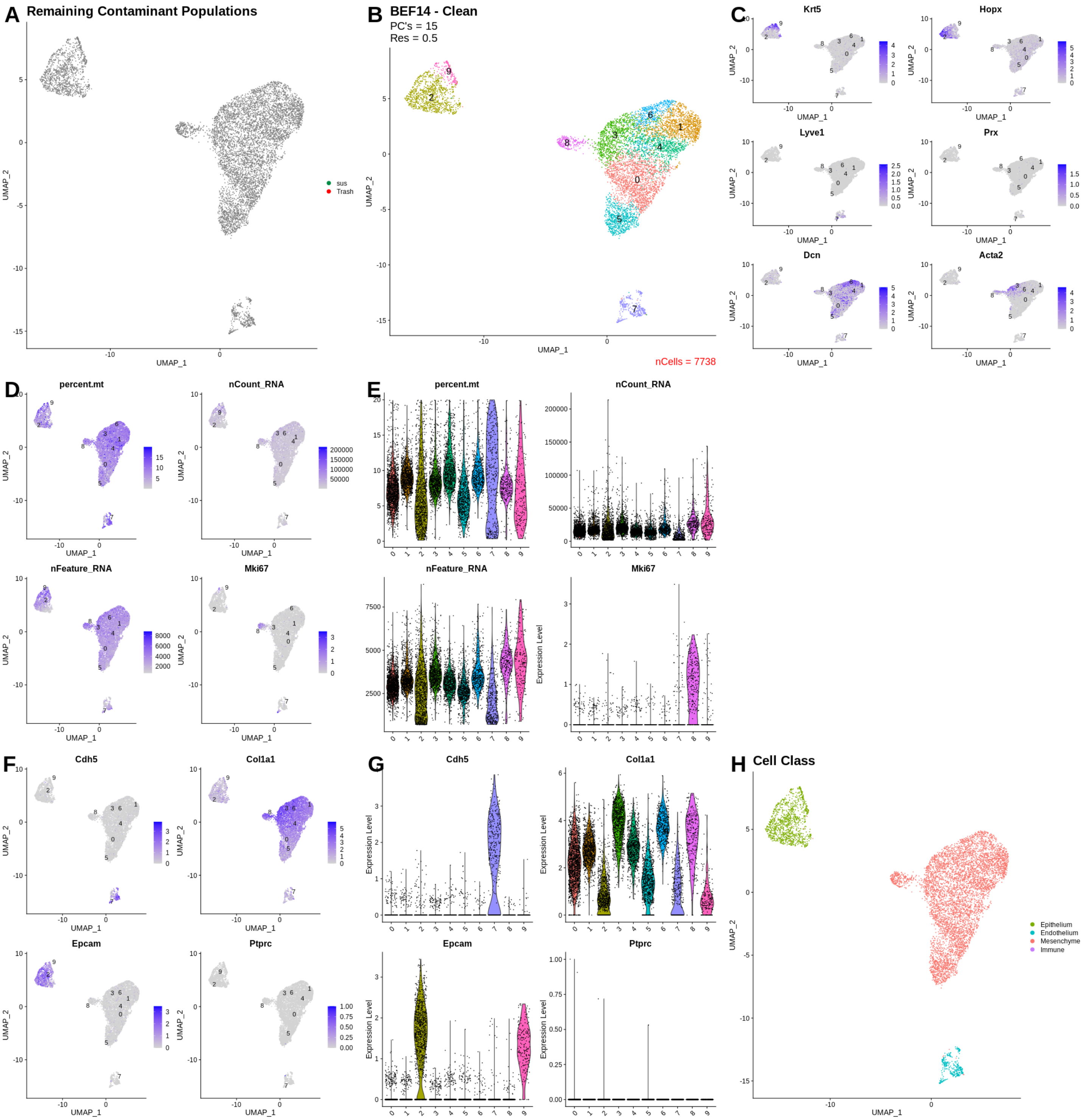
A
Remaining Contaminant Populations
sus
Trash
B
BEF14 - Clean
PC's = 15
Res = 0.5
nCells = 7738
C
Krt5
Hopx
Lyve1
Prx
Dcn
Acta2
D
percent.mt
nCount_RNA
nFeature_RNA
Mki67
E
percent.mt
nCount_RNA
nFeature_RNA
Mki67
F
Cdh5
Col1a1
Epcam
Ptprc
G
Cdh5
Col1a1
Epcam
Ptprc
H
Cell Class
Epithelium
Endothelium
Mesenchyme
Immune
UMAP_1
UMAP_2
Expression Level

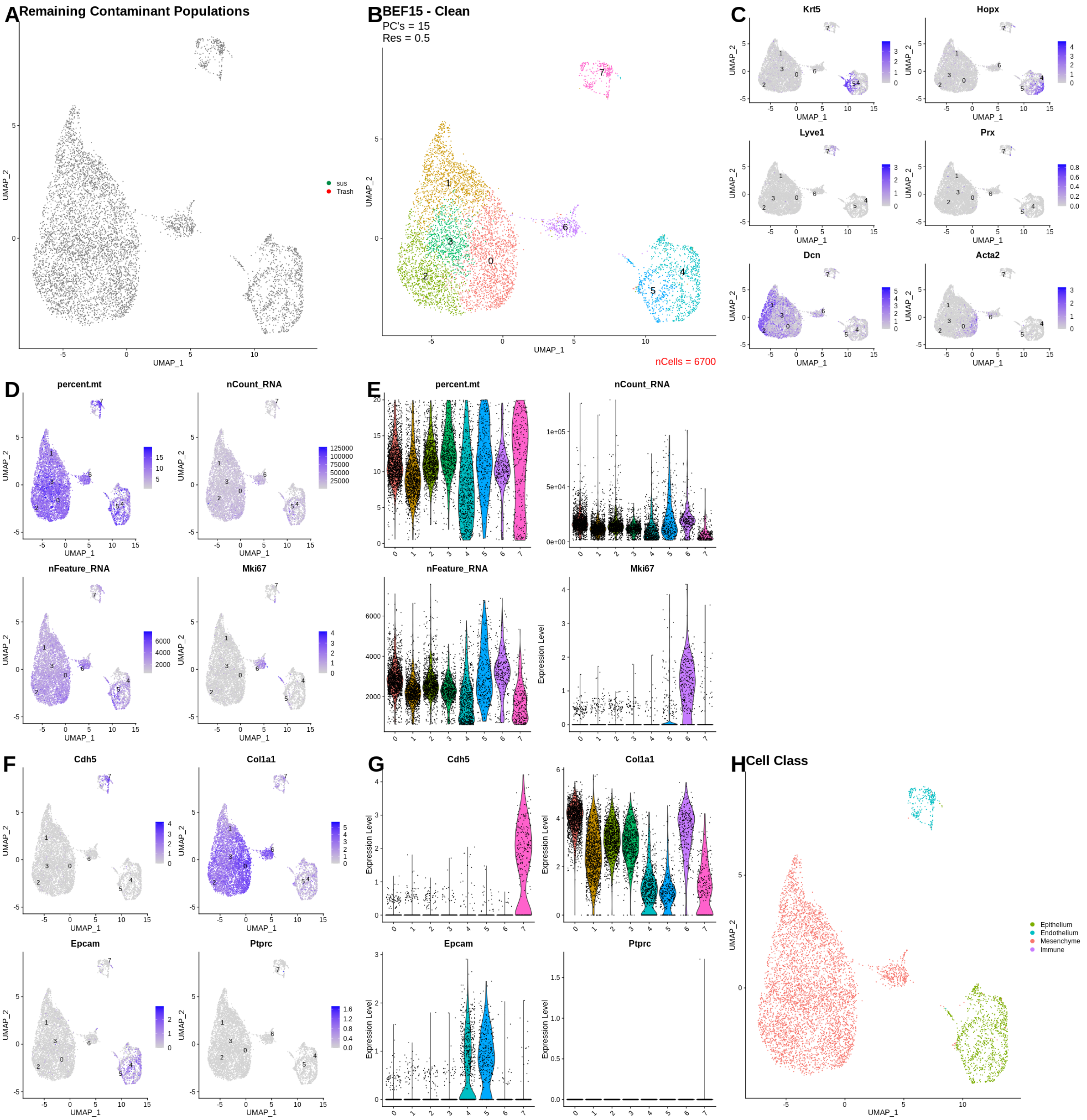

A Remaining Contaminant Populations
sus
Trash
B BEF15 - Clean
PC's = 15
Res = 0.5
nCells = 6700
C
Krt5
Hopx
Lyve1
Prx
Dcn
Acta2
D
percent.mt
nCount_RNA
nFeature_RNA
Mki67
E
F
Cdh5
Col1a1
Epcam
Ptprc
G
H Cell Class
Epithelium
Endothelium
Mesenchyme
Immune
UMAP_1
UMAP_2
Expression Level

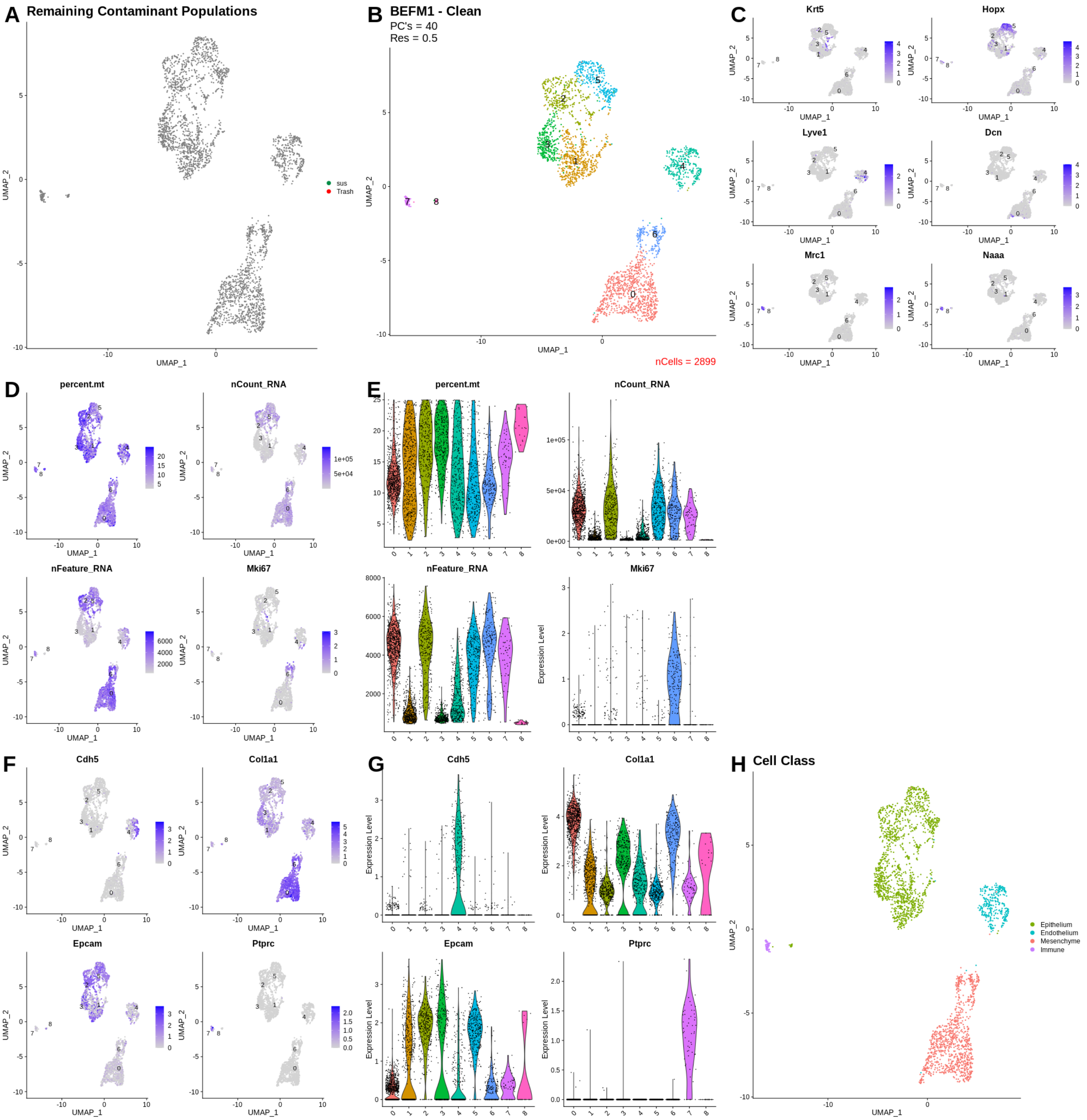
A
Remaining Contaminant Populations
UMAP_1
UMAP_2
sus
Trash
B
BEFM1 - Clean
PC's = 40
Res = 0.5
nCells = 2899
C
Krt5
Hopx
Lyve1
Dcn
Mrc1
Naaa
D
percent.mt
nCount_RNA
nFeature_RNA
Mki67
E
percent.mt
nCount_RNA
nFeature_RNA
Mki67
Expression Level
F
Cdh5
Col1a1
Epcam
Ptprc
G
Cdh5
Col1a1
Epcam
Ptprc
H
Cell Class
Epithelium
Endothelium
Mesenchyme
Immune

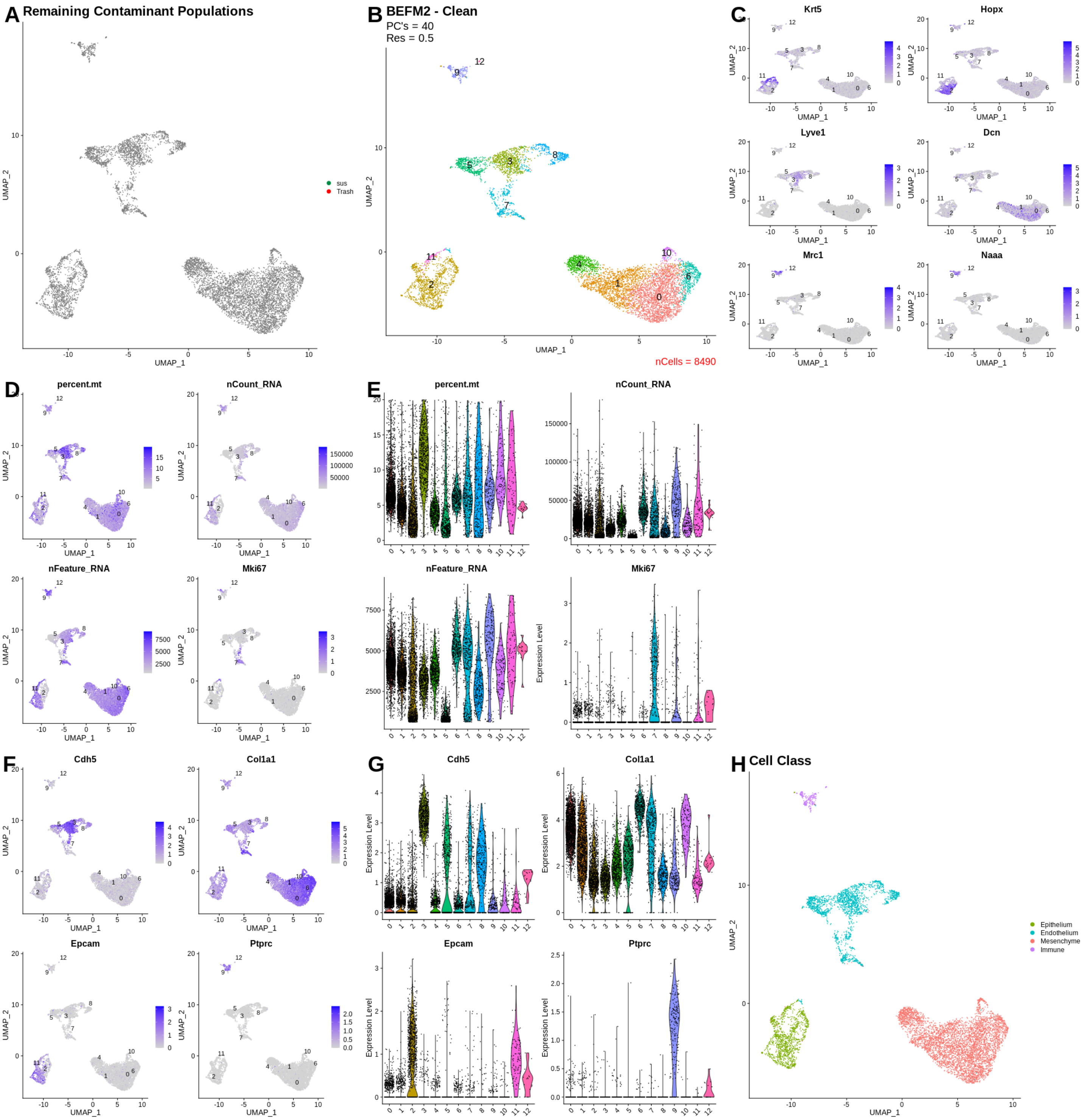

A Remaining Contaminant Populations
sus
Trash
B BEFM2 - Clean
PC's = 40
Res = 0.5
nCells = 8490
C
Krt5
Hopx
Lyve1
Dcn
Mrc1
Naaa
D
percent.mt
nCount_RNA
nFeature_RNA
Mki67
E
percent.mt
nCount_RNA
nFeature_RNA
Mki67
Expression Level
F
Cdh5
Col1a1
Epcam
Ptprc
G
Cdh5
Col1a1
Epcam
Ptprc
H Cell Class
Epithelium
Endothelium
Mesenchyme
Immune
UMAP_1
UMAP_2

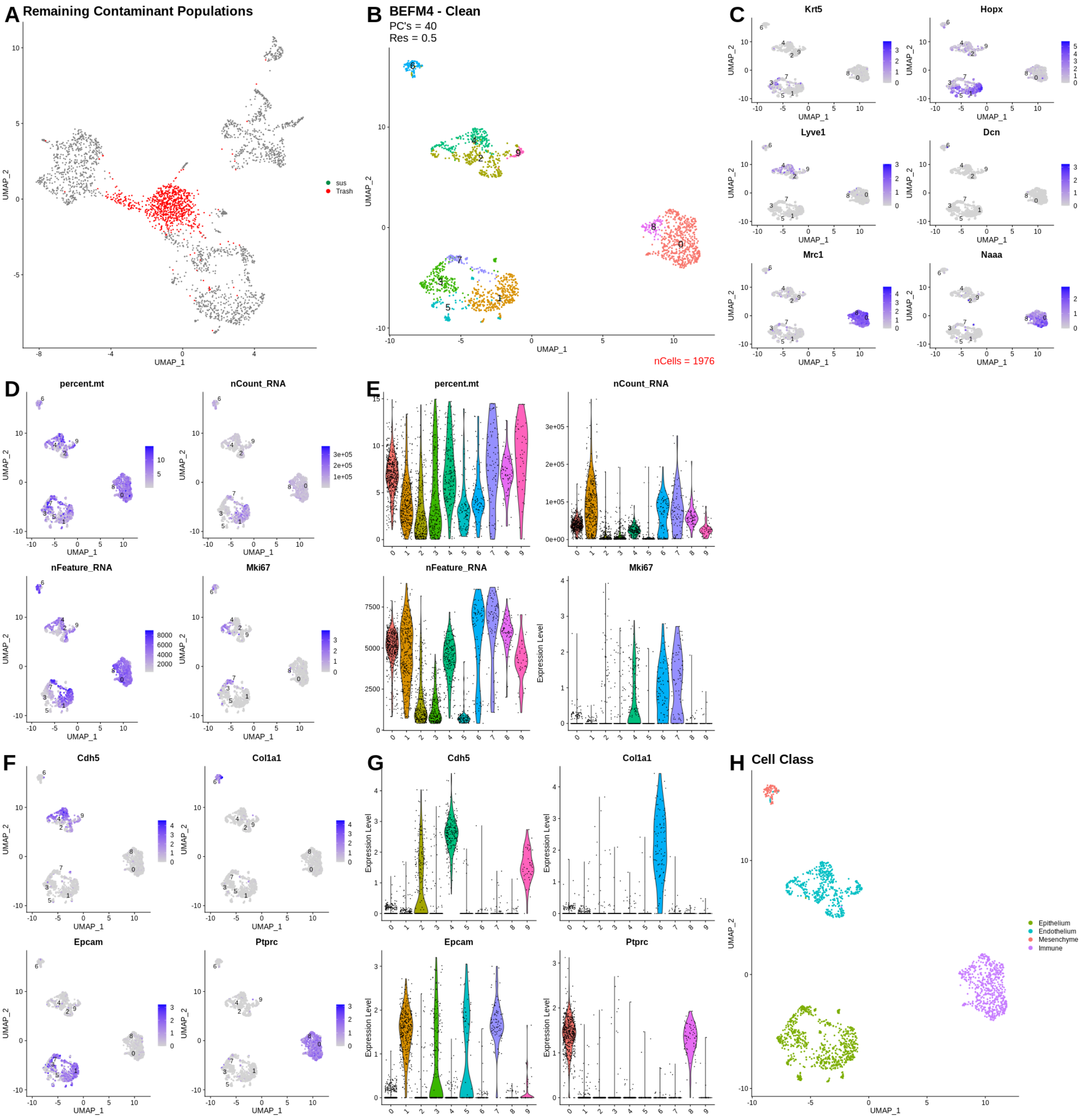
A
Remaining Contaminant Populations
sus
Trash
UMAP_1
UMAP_2
B
BEFM4 - Clean
PC's = 40
Res = 0.5
nCells = 1976
C
Krt5
Hopx
Lyve1
Dcn
Mrc1
Naaa
D
percent.mt
nCount_RNA
nFeature_RNA
Mki67
E
Expression Level
F
Cdh5
Col1a1
Epcam
Ptprc
G
H
Cell Class
Epithelium
Endothelium
Mesenchyme
Immune

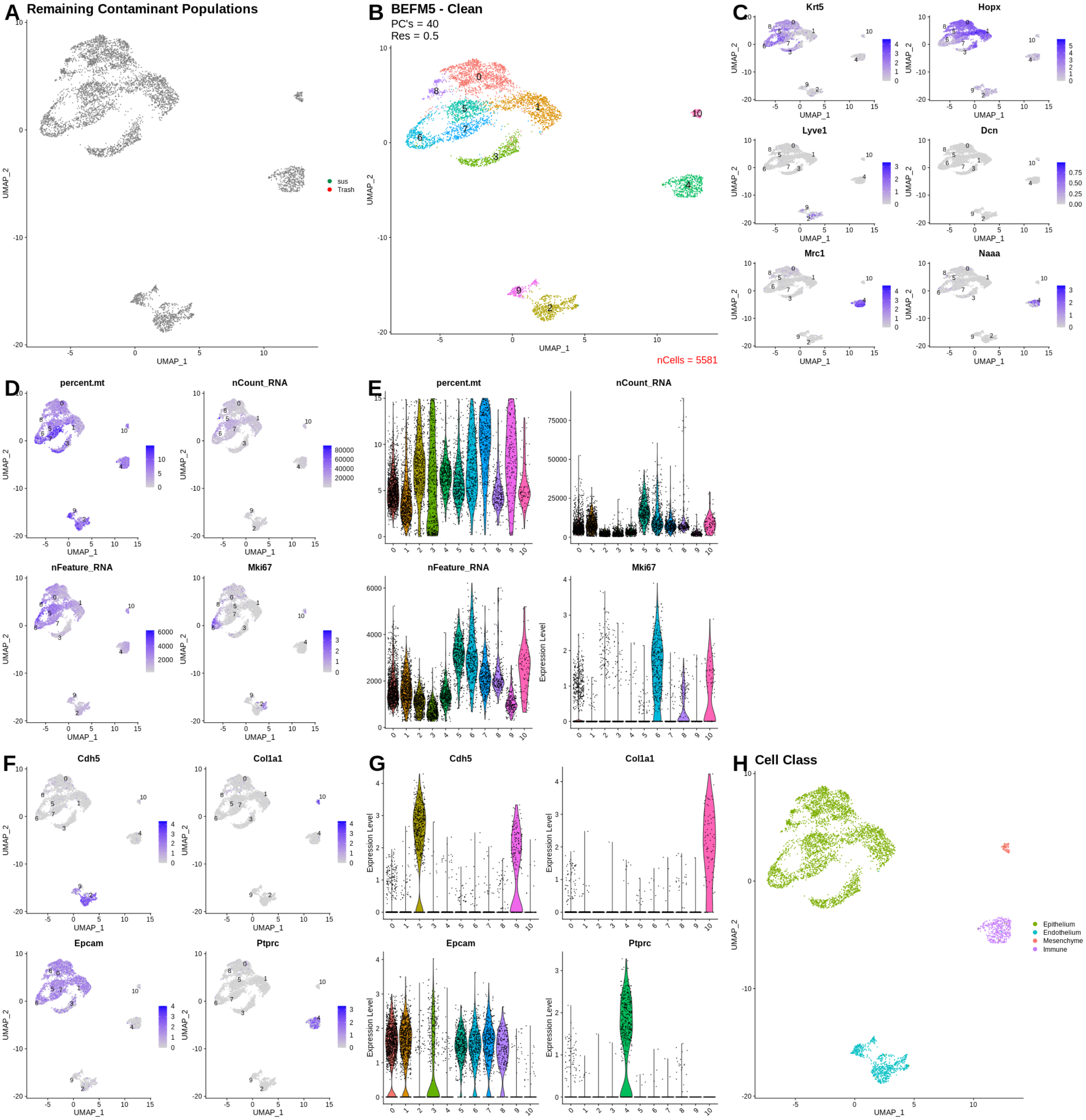

A Remaining Contaminant Populations
sus
Trash
B BEFM5 - Clean
PC's = 40
Res = 0.5
nCells = 5581
C
Krt5
Hopx
Lyve1
Dcn
Mrc1
Naaa
D
percent.mt
nCount_RNA
nFeature_RNA
Mki67
E
F
Cdh5
Col1a1
Epcam
Ptprc
G
H Cell Class
Epithelium
Endothelium
Mesenchyme
Immune
UMAP_1
UMAP_2
Expression Level

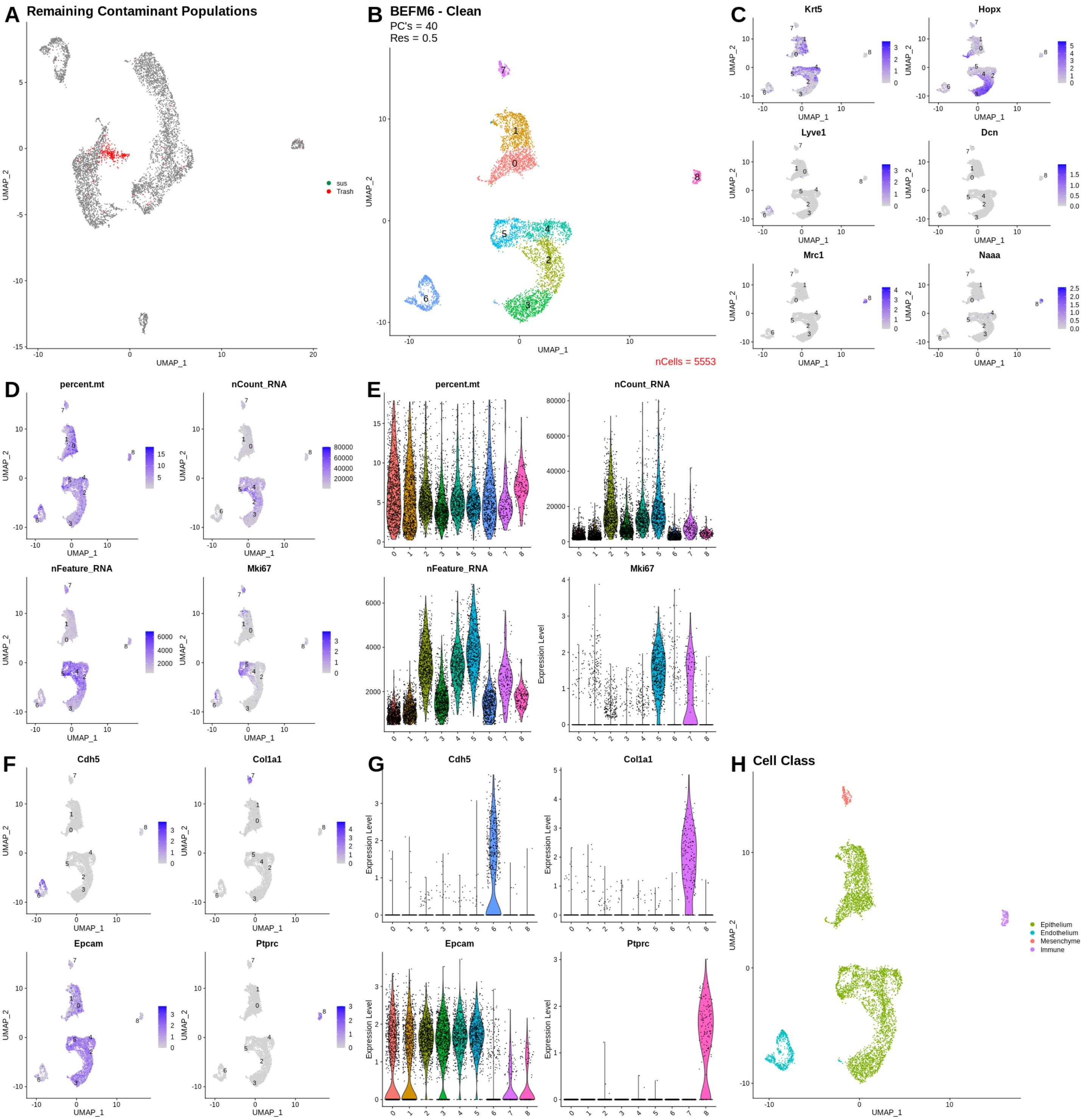
A Remaining Contaminant Populations
sus
Trash
B BEFM6 - Clean
PC's = 40
Res = 0.5
nCells = 5553
C
Krt5
Hopx
Lyve1
Dcn
Mrc1
Naaa
D
percent.mt
nCount_RNA
nFeature_RNA
Mki67
E
percent.mt
nCount_RNA
nFeature_RNA
Mki67
Expression Level
F
Cdh5
Col1a1
Epcam
Ptprc
G
Cdh5
Col1a1
Epcam
Ptprc
Expression Level
H Cell Class
Epithelium
Endothelium
Mesenchyme
Immune
UMAP_1
UMAP_2
